# The origins and functional effects of postzygotic mutations throughout the human lifespan

**DOI:** 10.1101/2021.12.20.473199

**Authors:** Nicole B. Rockweiler, Avinash Ramu, Liina Nagirnaja, Wing H. Wong, Michiel J. Noordam, Casey W. Drubin, Ni Huang, Brian Miller, Ellen Z. Todres, Katinka A. Vigh-Conrad, Antonino Zito, Kerrin S. Small, Kristin G. Ardlie, Barak A. Cohen, Donald F. Conrad

## Abstract

Postzygotic mutations (PZMs) begin to accrue in the human genome immediately after fertilization, but how and when PZMs affect development and lifetime health remains unclear. To study the origins and functional consequences of PZMs, we generated a multi-tissue atlas of PZMs from 948 donors using the final major release of the Genotype-Tissue Expression (GTEx) project. Nearly half the variation in mutation burden among tissue samples can be explained by measured technical and biological effects, while 9% can be attributed to donor-specific effects. Through phylogenetic reconstruction of PZMs, we find that their type and predicted functional impact varies during prenatal development, across tissues, and the germ cell lifecycle. Remarkably, a class of prenatal mutations was predicted to be more deleterious than any other category of genetic variation investigated and under positive selection as strong as somatic mutations in cancers. In total, the data indicate that PZMs can contribute to phenotypic variation throughout the human lifespan, and, to better understand the relationship between genotype and phenotype, we must broaden the long-held assumption of one genome per individual to multiple, dynamic genomes per individual.

**One-Sentence Summary:** The predicted rates, functional effects and selection pressure of postzygotic mutations vary through the human lifecycle.

## Main text

The effects of age ravage all tissues of the body, but the pace and consequences of age-related decay varies among tissues and people. The accumulation of DNA damage is thought to be a primary agent of age-related disease(*1*), and surveys of postzygotic mutations (PZMs) in normal tissues (e.g., blood(*2–4*), brain(*5*), and skin(*6, 7*)), and across the body (*8–10*), have found PZMs to be pervasive across the genome and individuals. However, beyond cancer there are few conditions where PZMs are known to have a causal role. Due to the high cost and technological challenges of PZM studies, a general understanding of how and when mutation affects the function of specific cell and tissue types is essential for defining research priorities. One way to prioritize hypotheses about mutation and disease is to systematically characterize the fitness consequences of PZMs across a broad range of tissues. Surveys of normal tissues have found that PZMs appear to accrue neutrally(*9, 11*), but positive and negative selection do occur in specific genes and cellular contexts, suggesting PZMs affect cellular function.

Another fundamental question is how the timing of mutation modulates risk for diseases. As clearly demonstrated in oncology, it is possible to detect disease-causing PZMs and augment clinical care years before clinical disease is recognized(*12, 13*). If PZMs that confer risk for disease accrue across the lifespan, the PZM profile in a healthy individual could contain actionable prognostic information. While the relative contributions of prenatal and postnatal PZMs to disease risk are unclear, due to the massive cell proliferation during development, prenatal PZMs have the potential to affect many cells, and thus, play an important role in disease.

The vast majority of PZM research has been single-tissue studies largely focused on tissues that are easily accessible, such as blood, liver, skin and colon. An exciting next generation of PZM studies now examine PZMs across multiple tissues within an individual(*8*–*10, 14*). However, the relatively small number of individuals and tissue types used in such studies have limited the ability to ascribe sources of mutation variation among individuals or provide detailed descriptions of embryonic mutations that occur after the first few cell divisions. To expand our knowledge of PZMs in normal tissues, we developed a suite of methods called Lachesis to identify single-nucleotide PZMs from bulk RNA-seq data and predict when the mutations occurred during development and aging (**Fig. S1**). We ran the algorithm on the final major release of GTEx, a collection of RNA-seq data from 17,382 samples derived from 948 donors across 54 diverse tissues and cell types, to generate one of the most comprehensive databases of PZMs in normal tissues (**tables S1** and **S2**). We used this atlas, and the rich metadata on GTEx donors, to characterize sources of variation in PZM burden among individuals and unveil the spatial, temporal, and functional variation of PZMs in normal development and aging. This reference of normal PZM variation will be instrumental to identifying abnormal variation associated with disease.

### DNA PZMs are accurately detected in bulk tissue RNA-seq

We evaluated the accuracy of the algorithm using several *in silico* and experimental methods (**fig. S4**, **tables S3**-**S6**). For experimental validation, we obtained four independent DNA- and RNA-based validation datasets generated from the same tissue samples as the primary data covering 296 unique genomic sites across 95 samples. The original RNA-seq VAF was highly correlated with the validation DNA VAF (Spearman’s ρ = 0.82, *P*-value = 2.3E-25) suggesting RNA-seq based VAFs are representative of true mutant cell frequencies. PZMs with VAFs as low as 0.16% and PZMs found in multiple tissues and multiple donors were validated. The average FDR across all validation datasets was 27% and is lower than published methods for detecting PZMs from RNA-seq (e.g., 34% - 82%(*8, 10, 15*)) (**Fig. S1E** and **table S3**). Since mutations may fail to validate due to spatial variation in mosaicism, the FDRs may be overestimates. A small subset of samples (~5%) had an extraordinarily high PZM burden; we used additional experimental validation data from 1,509 putative PZMs in these samples to demonstrate that these outliers were likely technical artifacts, and not hypermutated tissues (**Supplementary result 2.1**).

We used power simulations to estimate the algorithm’s sensitivity. As expected, simulated PZMs with larger VAFs and higher coverage had higher PZM detection power. At the middle quintile of coverage ([673, 1395) fold coverage), PZMs with VAFs as low as 0.66% could be detected in at least 90% of simulations, suggesting the method has reasonable sensitivity (**Fig. S1F**).

### PZMs are pervasive and highly variable among donors and tissues

Following sample and PZM quality control, 56,585 PZMs were detected with variant allele frequencies (VAFs) as low as 0.04% and a median VAF of 0.5% (**table S7**). 100% of the donors and 77% of the tissue samples had detectable mosaicism (**table S2**). The median mutation burden per tissue ranged from 0.03 PZMs/Mb in cerebellar hemisphere to 0.47 PZMs/Mb in liver (**Fig. 1A**). Strikingly, the observed mutation burden was more variable within a tissue than between tissues (mean median absolute deviation (MAD) within a tissue = 0.07 PZMs/expressed Mb; MAD across tissues = 0.02 PZMs/expressed Mb). This observation suggests that processes generating detectable PZMs may be more variable across donor than across tissue types.

**Fig. 1.**
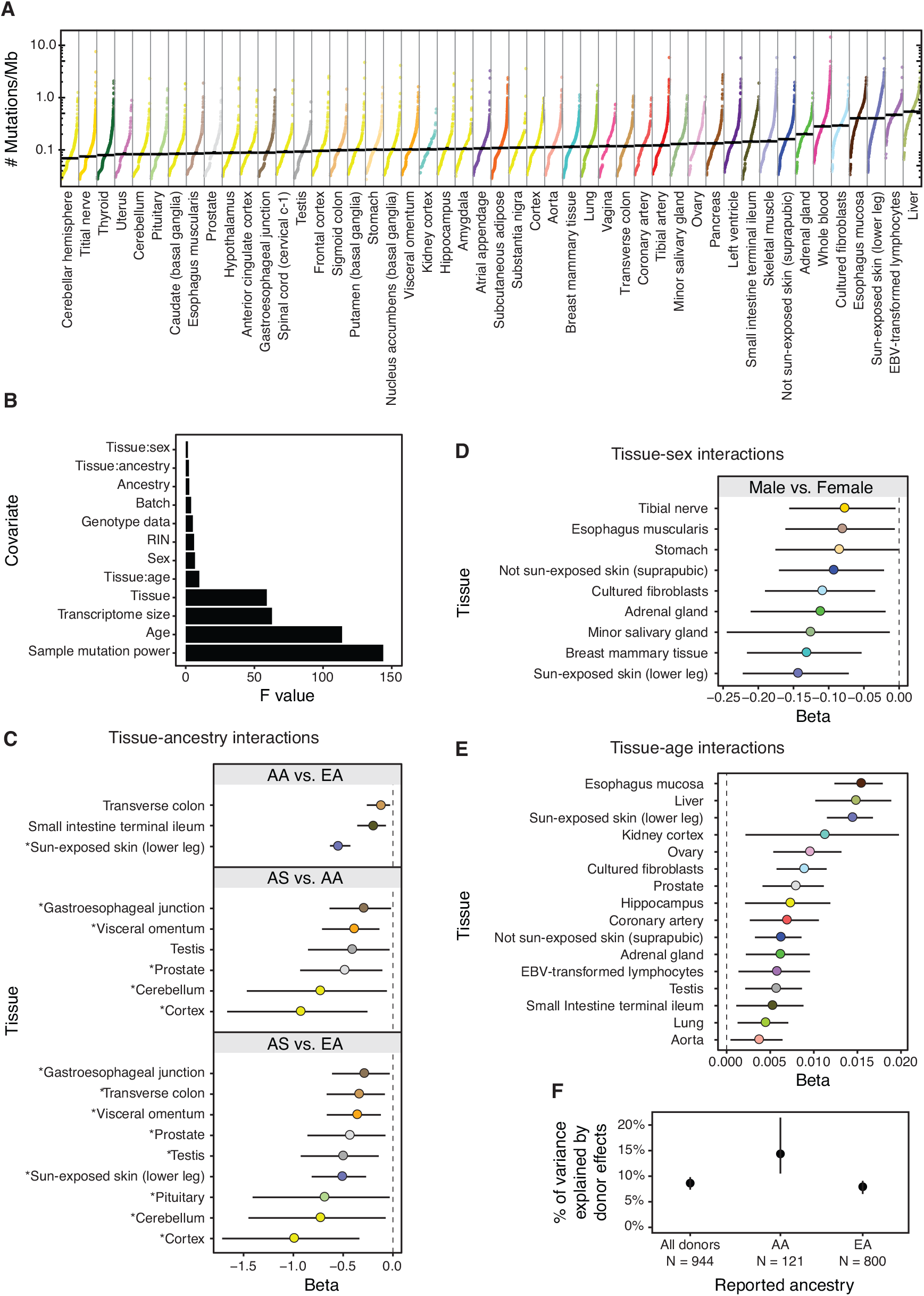
PZM burden is correlated with biological and technical variables. Each datapoint represents a single tissue sample and is colored by tissue. Median PZM burden in a tissue denoted by horizontal black line. Tissues are sorted by increasing median PZM burden. A pseudocount of 1 mutation was added to each sample before normalization and log transformation for visualization. (**B**) We fit a regression model for single-tissue PZM burden using 12 covariates and 48 tissues. ANOVA F statistics for each covariate in the model. Larger F statistics correspond to greater explanatory power of the covariate. (**C**) Regression coefficients of tissue-ancestry interactions and (**D**) tissue-sex interactions (female used for reference level) indicate strong effects of ancestry and sex on PZM burden. AA = African American. AS = Asian American. EA = European American. * in C denote trends that are consistent with cancer incididence trends (**E**) Significant positive tissue-age interaction effects were detected for 16/48 (33%) tissues. (**F**) Variance component estimates of donor-specific random effects on PZM burden indicate that 8%-15% of variation among tissues can be ascribed to donor effects, which could be genetic and environmental. Dashed vertical lines at beta = 0 in interaction plots denote no association between mutation burden and interaction. Error bars represent 95% CIs. Tissues are colored using the GTEx coloring convention (see **table S8** for a complete legend).

#### PZM burden is correlated with biological and technical variables

To partition and quantify potential sources of single-tissue PZM burden, we fit linear models relating technical and biological metadata to single-tissue PZM burdens and selected the best fitting model identified from detailed model comparisons (**Methods section 1.3**). The model contained twelve covariates and explained 48% of the variation in mutation burden. Significant sources of variation were biological (age, tissue, and interactions of tissue with age, sex, and reported ancestry) and technical (e.g., mutation detection power and RNA extraction batch) (**Fig. 1B-E**). 20.8% (10/48) of tissues showed significant associations with reported ancestry, including, as expected, a much lower burden of mutation in sun-exposed skin in African Americans and Asian Americans compared to European Americans (*8*). The incidence rates of cancer types affecting these tissues have ancestry associations that are consistent (i.e. in the same direction as) with the mutation burden associations in 83% (15/18) of comparisons(*16*), suggesting that variation in PZM burden in normal tissues may contribute to differences in cancer risk among ancestries (**Fig. 1C**, **table S9)**. Unexpectedly, males had lower burden in all three skin-related sample types compared to females. Age was positively associated with 33% (16/48) of tissues and was the strongest for esophagus – mucosa, liver, and sun-exposed skin. We note that power may have been too low to detect some associations, e.g., there were few young GTEx brain donors.

Extending this model to include a random donor effect, we estimated that 8.8% of variation in PZM burden can be attributed to systematic properties of donors that extend across some or all tissues of a donor, even after controlling for metadata such as age and sex. We found that this donor variance component estimate was larger in African Americans (14.1%; 95% confidence interval (CI): 10.5-21.5%) than in European Americans (8%; 95% CI: 6.5-9.1%) (**Fig. 1F**). These unexplained donor-specific effects could have both genetic and environmental bases. Notably, a recent study estimated that 5.2% of variance in germline mutation rate could be attributed to family-specific effects(*17*). In total, our results indicate that variation in PZM rate among individuals is less constrained than variation in germline mutation rate and that there is considerable scope for heritable variation in observable PZM burden. The inability of the models to explain all variation imply there are additional factors associated with detectable mutation burden and/or stochasticity plays a major role in mosaicism(*18, 19*).

### Mutation spectra is variable across tissues and reflect known biological processes

Diverse processes mutate the human genome with characteristic mutational signatures(*20*). Thus, the observed mutation spectra can provide insight on the types and relative activities of the unobserved mutation processes that occurred. We estimated the contribution of canonical mutation signatures for each tissue. Due to the relatively low number of detected mutations, mutation spectra were reliably deconstructed for only four tissues/cell types (**fig. S10A**). Consistent with expectations and previous studies(*3, 6, 7*), the mutations were resolved into mutational signatures associated with age in all tissues and ultraviolet light exposure in skin-related tissues (**fig. S11**).

For a higher powered, but coarser-grained analysis of mutation spectra, we assessed the frequency of the six base substitutions across all tissues (**fig. S10B**). Mutation spectra were highly variable across tissues suggesting that mutational mechanisms and their relative activity may vary across the human body. C>T was the most common mutation type across tissues whereas C>G and T>A were the least common. Hierarchical clustering with bootstrap resampling of the mutation types revealed two significant large clusters: cluster A (marked by depleted T>G) and cluster B (marked by elevated T>G). Cluster membership was associated with mutation burden suggesting the underlying mutation mechanisms may be coupled to the frequency of mutagenic events (*P*-value = 3.8E-2, Mann-Whitney *U* test). Additionally, Cluster B was enriched with neural ectoderm tissues compared to cluster A (*P*-value = 7.7E-6, Fisher’s exact test). These clusters could not be attributed to differences in sample processing (**Supplementary result 2.3**, **fig. S12**). We speculated that these clusters may reflect differences in the relative contributions of mutations acquired during prenatal development and mutations that accrue during age-related tissue renewal (e.g., during mitotic divisions in epithelial tissues). To further study the properties of prenatal and postnatal PZMs, we developed methods to define the developmental origin of each PZM.

### The developmental origins of prenatal PZMs

#### Multi-tissue PZMs exhibit prenatal properties

We defined a multi-tissue PZM as a PZM that was detected in at least two tissues from the same donor. Since the PZM mutation burden was relatively low across tissues (**Fig. 1**) and PZMs are predominantly under neutral selection(*11*), we hypothesized that a multi-tissue PZM was the result of a single PZM that occurred in a common ancestor of the mutated tissues. Since the common ancestors of any set of GTEx tissues (excluding cell lines) occurred before the end of organogenesis, multi-tissue PZMs may have occurred prenatally. Consistent with this hypothesis, we found several lines of evidence suggesting the multi-tissue PZMs occurred prenatally (**Supplementary result 2.4**, **figs. S13** and **S14**). We denote these mutations as prenatal PZMs, while all other mutations are called postnatal PZMs.

#### PZM burden and spectra vary throughout prenatal development with most mutations occurring during early embryogenesis

To determine when and where PZMs occur in prenatal development, we developed a method called LachesisMap to map the origin of 1,864 prenatal mutation events (**Fig 2A**, **figs. S5-S7**, **Methods section 1.5**). Briefly, the method takes as input a directed rooted tree representing the developmental relationships among the tissues and a list of multi-tissue PZMs and maps the PZMs to the tree that accounts for differential mutation detection power across the genome, human body, and developmental tree. The algorithm outputs a list of edge weights that represent the estimated fraction of PZMs that occurred in that spatiotemporal window of development.

**Fig. 2.**
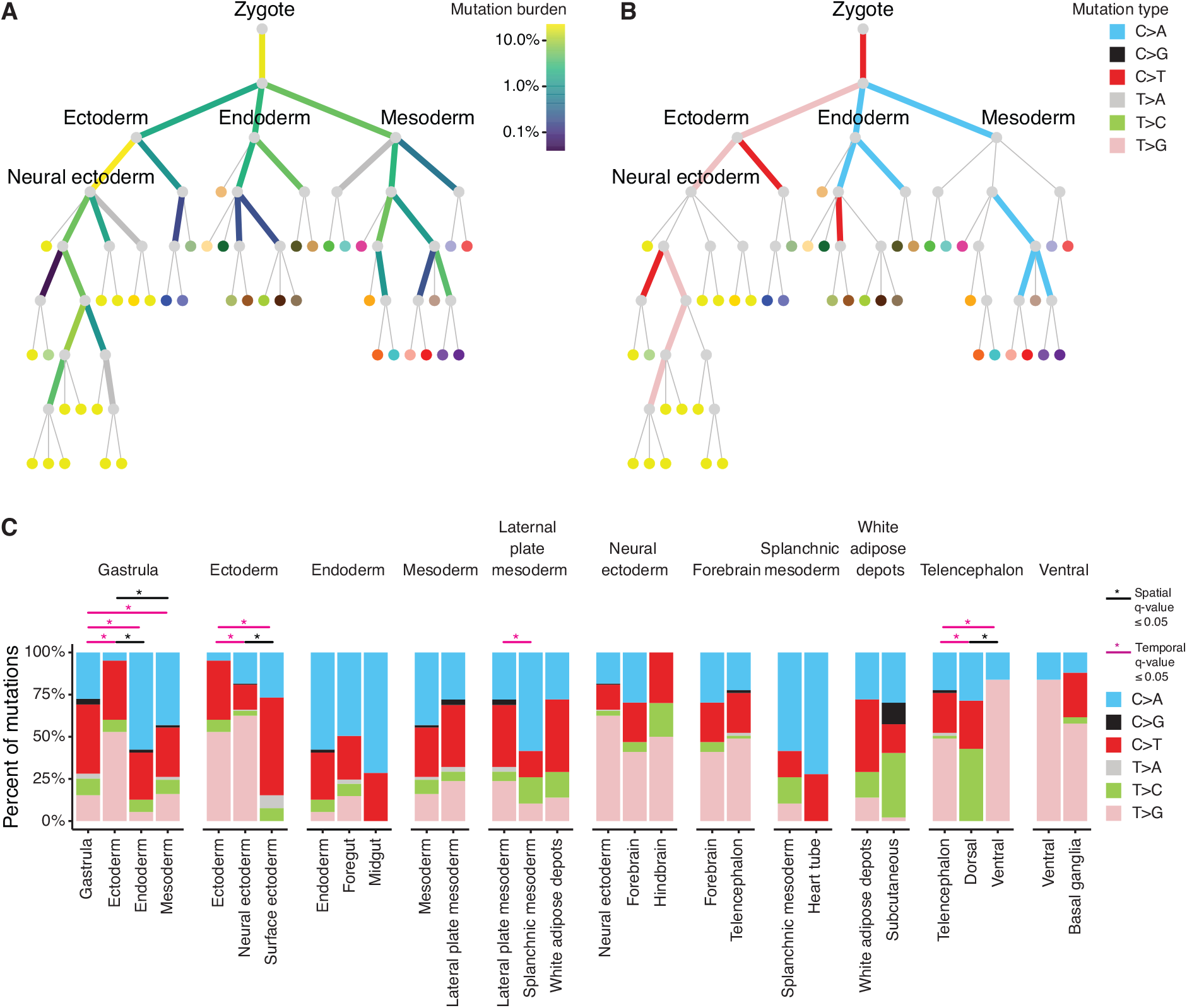
Mutation burden and spectra of prenatal PZMs across time and space. (**A**) Prenatal PZM mutation burden. Edge color represents the percent of prenatal PZMs mapped to that period in development. Thick gray edges are edges with limited mutation detection power. (**B**) Edge color represents the predominant mutation type of mutations mapped to that edge, as established by binomial testing. Thin gray edges are edges with no predominant mutation type. See **fig. S5A** for the full set of vertex labels. Adult tissues (leaves of tree) are colored using the GTEx coloring convention (see **table S8** for a complete legend). (**C**) Local variation in mutation spectra across developmental space and time. Each facet represents the mutation spectra observed in a parent edge (leftmost barplot) and its children’s edges. Statistically significant differences in mutation spectra are annotated with “*”.

The mutation burdens across developmental time and space were highly variable, with edge weights ranging from 0.04% to 23%, and appeared compatible with an exponential distribution (*P*-value = 0.56, Kruskal-Wallis test). The top two edge weights, representing 41% of prenatal mutation events, were the zygote to gastrula transition and the ectoderm to neural ectoderm transition, suggesting that most detectable prenatal mutations occur during early embryogenesis(*14, 21*). Of critical note, the edge mutation burdens were not explained by differential edge mapping power across the developmental tissue tree (**Supplementary results 2.5 and 2.6**, **fig. S15** and **S16**).

We next asked if the mutational processes, as proxied by their mutation spectra, varied over development, using binomial tests to establish the “predominant” mutation type on each edge. There was a strong dichotomy between ectoderm lineages, which tended to have T>G mutations, and endoderm and mesoderm lineages which tended to have C>A mutations (**Fig. 2B**). These observations could not be attributed to differences in sample processing (**Supplementary result 2.3**, **fig. S12**).

In addition to global changes in mutation across the tree, we also examined local changes by comparing mutation spectra between sibling edges (i.e., local spatial differences) and parentchild edges (i.e., local temporal differences) (**Fig. 2C**). Significant spatial and temporal variation was detected during gastrulation and in ectodermal lineages. Differences in mutation spectra across developmental space (N = 4/8 (50%) sibling edge comparisons) occurred at similar rates as differences along developmental time (N = 8/18 (44%) parent-child comparisonsmethod for the study of somatic) (*P*-value = 1.00, Fisher’s exact test).

Together, these results suggest that the mutational mechanisms that operate during development may vary across space and time. Although published data is limited, others have also detected variation in mutation spectra in fetal stem cells in humans(*22*) and during early embryogenesis and gametogenesis in mice(*23*).

We repeated these analyses using a simplified germ layer tree and observed similar results as the full developmental tissue tree (**fig. S17**), suggesting that the development tree definition does not substantially affect the results.

### The functional consequences of PZMs across the human lifespan

The GTEx PZM atlas provides a unique opportunity to compare the quality and fitness consequences of mutations that arise at different stages of the human life cycle. First, we annotated the PZM atlas with CADD, a widely used machine learning classifier of genetic variation(*24*). The CADD score of a genetic variant is a quantitative prediction of deleteriousness, measured on an evolutionary timescale. We performed a series of systematic comparisons of PZM CADD scores to identify differences across mutation VAF, developmental time, developmental location, and tissue type.

Strikingly, when comparing prenatal and postnatal PZMs, we found a major effect of VAF on the distribution of CADD scores (**Fig. 3A** and **fig. S18A**). For prenatal PZMs, low VAF PZMs were much more deleterious than high VAF PZMs (odds ratio = 1.9, *P*-value = 2.6E-7, Fisher’s exact test), while no such difference was observed for postnatal PZMs (*P*-value-value = 0.24, Wald Test). Furthermore, we found that for low VAF PZMs, deleteriousness decreased over time (odds ratio = 0.58, *P*-value = 1.4E-9) but remained constant for high VAF PZMs (*P*-value = 0.15). This result suggests that mutations that appear deleterious on an evolutionary timescale may be benign or even beneficial to a growing fetus so long as the mutation remains in a small fraction of cells.

**Fig. 3.**
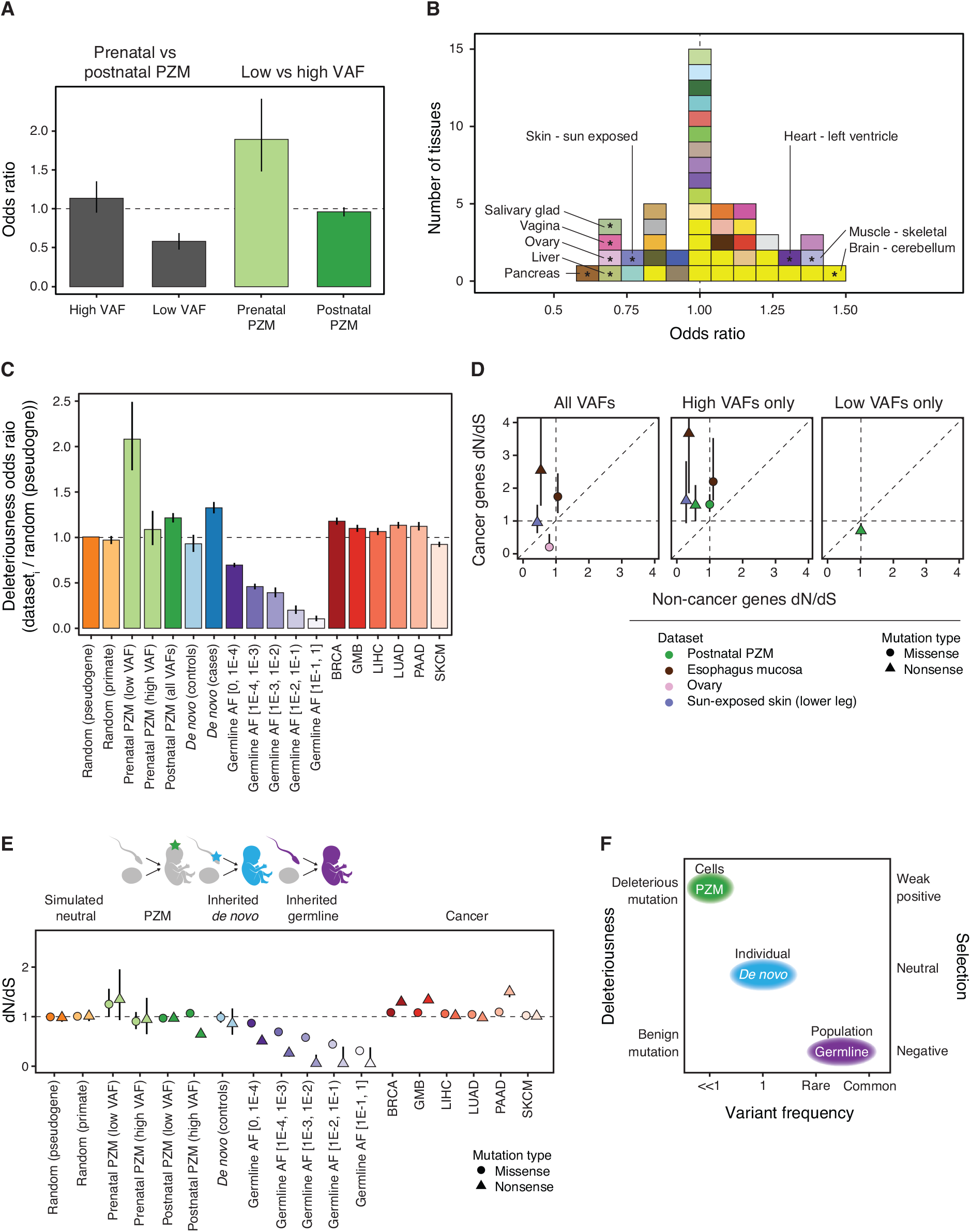
Deleteriousness and selective pressure changes as a function of VAF, space, time, and classes of genetic variation. (**A**) Relative odds of detecting deleterious mutations across developmental time (gray bars) and VAF bins (green bars). (**B**) Histogram of the odds of detecting deleterious postnatal PZMs in each tissue compared to the average tissue. Tissues are colored using the GTEx coloring convention (see **table S8** for a complete legend). Tissues with significant odds ratios (at q-value ≤ 0.05) are marked with “*” and labeled with their name. Vertical dashed line at odds ratio = 1 indicates no difference in odds. (**C**) Relative odds of detecting deleterious PZM mutations compared to different classes of genetic variation. Lines at odds ratio = 1 indicates no difference in odds of detecting deleterious mutations compared to reference group. Error bars represent 95% CIs. (**D**) Comparison of postnatal PZM selection pressure in cancer and non-cancer genes. For clarity, only PZM datasets that had different selection pressure between cancer and non-cancer genes are shown. Left: PZM datasets that had variable selection when using all mutations; middle: high VAF mutations; right: low VAF mutations. (**E**) dN/dS values for classes of genetic variation, as in (C). CIs are plotted behind each datapoint and are sometimes smaller than the datapoint size. (**F**) Model for the association between variant frequency and deleteriousness and selection pressure. The data suggests negative correlations between mutation frequency in a population and deleteriousness and selection. Variation in deleteriousness and selection may be driven by differential forces that act on cells, individuals, and populations. dN/dS = 1 indicates neutral expectation. AF = allele frequency. BRCA = breast invasive carcinoma. GBM = glioblastoma multiforme. LIHC = liver hepatocellular carcinoma. PAAD = pancreatic adenocarcinoma. SKCM = skin cutaneous melanoma.

Next, we asked if deleteriousness varied across the adult human body by comparing postnatal PZMs in each adult tissue. PZM deleteriousness was similar across tissues; however, there were a few exceptions (**Fig. 3B**). PZMs in 6/48 (13%) tissues were significantly less deleterious than the average tissue and 3/48 (6%) tissues were more deleterious. When analyzed together, the PZMs from all brain regions were also more deleterious than average (*P*-value = 0.02, Fisher’s exact test). Interestingly, the tissues that were enriched for deleterious mutations were tissues composed of a large fraction of postmitotic cells (odds ratio CI: [1.5, ∞), *P*-value = 0.01, Fisher’s exact test). In contrast, tissues that rely on renewal from differentiated cells were depleted of deleterious mutations(*25*). This data suggests that the maintenance of genome integrity in a tissue may be coupled with its tissue homeostasis strategy and thus may provide a way for tissues to suppress tumor formation(*26*).

Finally, to provide context for our results, we compared the deleteriousness of GTEx PZMs to other classes of single-nucleotide genetic variation: 1) random mutations (simulated from two different models of neutral evolution), 2) standing germline variation (from gnomAD, a comprehensive database of germline genetic variation(*27*)), 3) inherited *de novo* mutations from cases and control probands (from denovo-db, a curated database of *de novo* mutations(*28*)) and 4) somatic mutations observed in cancer (from TCGA, a comprehensive database of cancer somatic mutations)(*29*).

Surprisingly, the low VAF prenatal PZMs were the most deleterious class of genetic variation investigated (**Fig. 3C** and **fig. S18**). Using the simulated neutral mutations as a reference, we found that postnatal PZMs, *de novo* mutations in cases, and somatic cancer mutations to be significantly enriched for deleterious mutation. *De novo* mutations in controls and high VAF prenatal PZMs were not statistically different from simulated neutral mutations. Inherited germline variants were significantly depleted of deleterious mutations, with the extent of depletion increasing with population frequency. These observations were recapitulated in 3 validation datasets that used a variety of nucleic acid sources and variant calling methods (**Supplementary result 2.8**).

Similar results were observed using alternative metrics for deleteriousness (SIFT(*30*) and PolyPhen(*31*)) (data not shown).

### The selective constraint on the transcribed exome varies throughout the human lifespan

The deleteriousness results suggest that selection pressure may be different across classes of genetic variation. We investigated this hypothesis by estimating the selection pressure on PZMs and other classes of genetic variation using dN/dS, a normalized rate of nonsynonymous to synonymous mutations(*32*). dN/dS values greater than one are interpreted as evidence for positive selection, while negative selection can lead to dN/dS values less than one. Using *dNdScv,* a method for the study of somatic evolution(*11*), we assessed dN/dS across VAF, developmental time, developmental location, and tissue type, and contextualized the results by comparing selection pressures on PZMs to other classes of genetic variation.

To determine if differences in selective pressure may explain differences in CADD scores of prenatal and postnatal PZMs, we compared dN/dS for prenatal and postnatal PZMs as a function of VAF. For high VAF PZMs, prenatal and postnatal mutations both fit a model of neutral accumulation. However, for low VAF PZMs, the remarkable prenatal PZM class was under nominal positive selection (missense dN/dS = 1.25, *P*-value = 0.047) whereas postnatal PZMs were under nominal negative selection (missense dN/dS = 0.97, *P*-value = 0.045), suggesting selection pressure on low VAF PZMs may vary across the human lifespan. These results are consistent with the idea that high CADD score PZMs in early development may confer a survival advantage that is missing from or irrelevant to postnatal cells.

We next determined if selection pressure of postnatal PZMs varied across adult tissues. For most tissues of the body, single-tissue dN/dS was not significantly different than 1, consistent with previous work(*11*). However, dN/dS was higher for high VAF PZMs compared to low VAF PZMs for all tissues *en masse* and for three tissues/cell types individually (whole blood, EBV-transformed lymphocytes, and adrenal gland). Additionally, dN/dS estimates for high VAF postnatal PZMs were higher in cancer driver genes than non-cancer driver genes for all tissues *en masse,* sun-exposed skin and esophagus mucosa, tissues where the action of adaptive evolution has already been documented (*7, 33*) (**Fig. 3D**). These observations are consistent with the expectation that positive selection on a mutation may result in clonal growth, and indeed, we detected mutations associated with clonal hematopoiesis of indeterminant potential in the blood of individuals without apparent hematological malignancies (**fig. S23** and **table S11**).

#### Selection pressure varies across different classes of genetic variation

To contextualize the selection on PZMs, we compared PZM missense dN/dS estimates to other classes of genetic variation (**Fig. 3E**). Selection of mutations *within an individual* (in both normal and diseased states) was characterized by neutral to weak positive selection whereas selection on variants *within a population* was characterized by negative selection. Altogether, the deleteriousness and selection results suggest a stark dichotomy between growth within an individual versus growth within a population: the mutations that are selected for within an individual may be detrimental to the population (**Fig. 3F**).

### Characterization of germ cell PZMs

#### Construction of a catalog of germ cell PZMs throughout the germ cell life cycle

While a great deal is known about germline variation and *de novo* mutations, much less is known about the PZMs that seed these forms of inherited genetic variation. To better understand PZMs in germ cells, we characterized and contrasted the mutation burden, spectra, and deleteriousness of germ cell PZMs across the germ cell life cycle.

Due to cell composition differences between male and female gonads, PZMs in testes samples could be confidently mapped to germ cells but PZMs in ovary samples could not (**Supplementary result 2.10**, **fig. S24**, and **table S12**). Therefore, only testicular germ cell PZMs were analyzed further. Germ cell PZMs were classified into “gonosomal” (present in somatic and germ cells) and “germ cell-specific”. 571 germ cell PZMs were identified in bulk testis from 281 testis donors of which 12% were putative gonosomal PZMs and the remaining 88% were putative germ cell-specific PZMs.

Testicular germ cell PZMs represent the full reservoir of mutations that can be passed to progeny. We hypothesized that the selection pressures on spermatogenesis, fertilization, and prenatal development may alter the types of mutations that pass through each of these bottlenecks of life. To examine germ cell PZMs that passed the spermatogenesis bottleneck, we generated whole exome sequencing data on small 200-cell pools of ejaculated sperm and identified and validated 75 PZMs in the same genomic regions that we assessed in the GTEx RNA-seq samples (defined as the “allowable transcriptome”) (**Methods section 1.10.2** and **fig. S8**). To examine germ cell PZMs that completed prenatal development, we used ~17,000 *de novo* mutations in the allowable transcriptome from denovo-db(*28*).

The mutation spectra for each germ cell mutation dataset were statistically different from the others (**Fig. 4A** and **table S13**, Chi-square test). While C>T was the most common mutation type in all datasets, C>A was the most variable. Interestingly, hierarchical clustering of the spectra nested the classes in developmental order, indicating that bottlenecks may leave incremental changes to the mutation spectra (**Fig. 4A inset**).

**Fig. 4.**
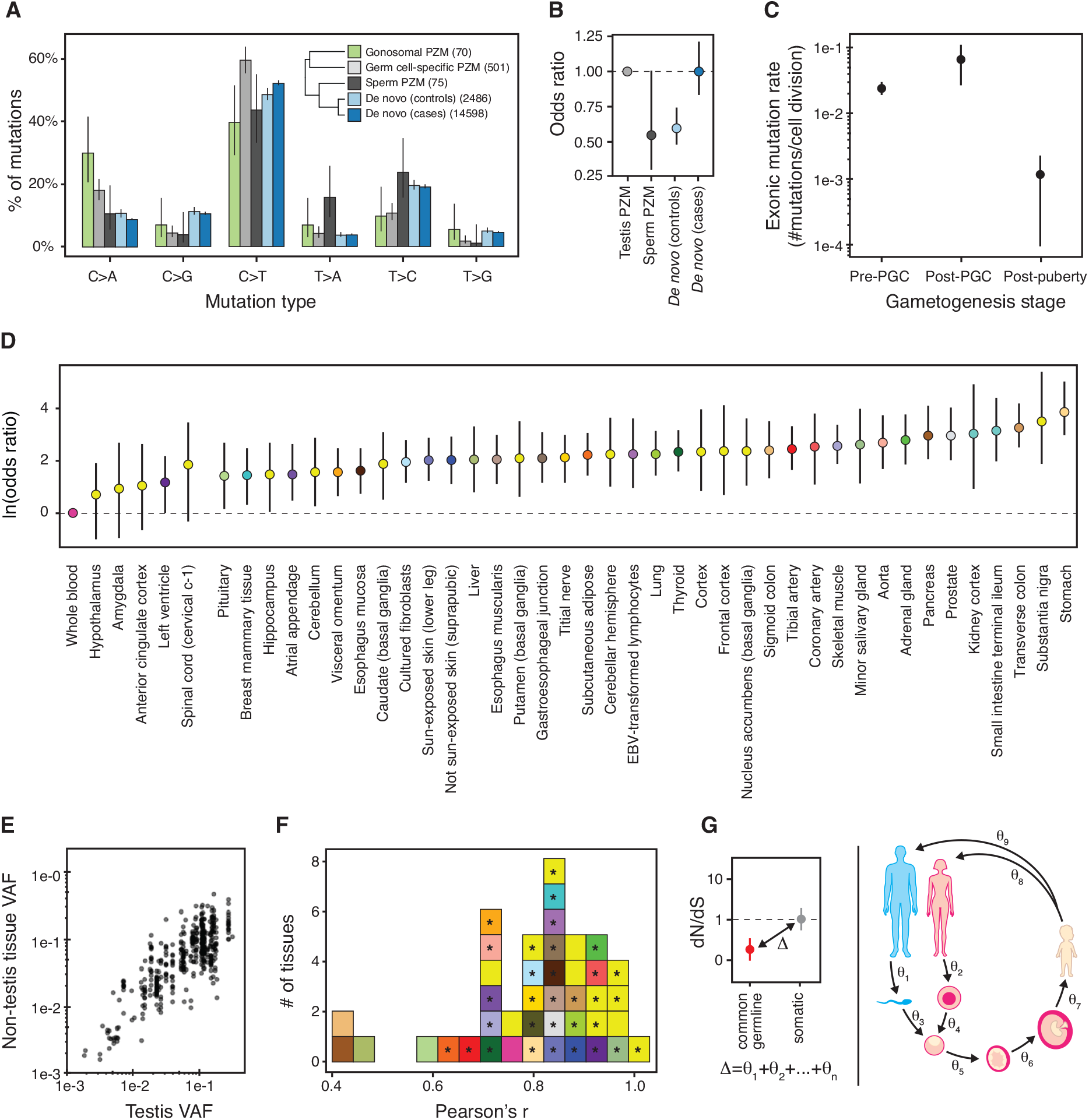
Germ cell PZM characteristics. (**A**) Mutation spectra of different germ cell mutation classesNumber of mutations used in each dataset is listed in the legend. **Inset**: Hierarchical clustering of germ cell mutation spectra. (**B**) Relative odds of detecting deleterious mutations across germ cell datasets compared to testis PZMs. Bars colored by dataset. Horizontal black line at odds ratio = 1 denotes no difference in odds. (**C**) Germ cell mutation rate varies during gametogenesis in males. (**D**) Majority of somatic tissues have a significantly higher odds of detecting a gonosomal PZM than blood. Natural log odds ratio for detecting a gonosomal PZM in each somatic tissue compared to blood. Dashed line at Y = 0 denotes no difference in odds. (**E**) Gonosomal PZM VAF in somatic tissue versus germ cell VAF. (**F**) Distribution of tissue-specific Pearson correlations of log10-transformed gonosomal PZM VAFs in each somatic tissue and testis. Significant correlations at q-value ≤ 0.05 marked with “*”. (**G**) Schematic of the difference in selective constraint between germline and somatic genetic variation partitioned into discrete stages of the life cycle. Error bars denote 95% CIs.

#### Deleterious mutations are likely purged during the germ cell life cycle

Consistent with the action of purifying selection on male germ cells, we found that mutation deleteriousness decreased over the germ cell life cycle when comparing testicular germ cell PZMs and *de novo* mutations in controls (**Fig. 4B**). In contrast, *de novo* mutations from cases were just as likely to be deleterious as testis PZMs. To replicate these observations, we performed a similar analysis using only DNA-based measurements from published datasets (**Supplementary result 2.10.3**). Encouragingly, we found that both the fraction of coding mutations and the odds of detecting a deleterious mutation decreased over the germ cell life cycle in the independent datasets (**fig. S25**). Donor age was not associated with PZM deleteriousness in each dataset.

#### The mutation rate during male gametogenesis is dynamic

We estimated the mutation rate (i.e., number of mutations in the transcriptome per cell division) for each of three major stages during male gametogenesis (**Methods section 1.11**). Consistent with previous work(*34*), the observed mutation rate was higher in prenatal timepoints than the postnatal timepoint (**Fig. 4C**). The observed lower mutation rate during adulthood may be a strategy to limit the number of deleterious mutations that are passed to the next generation. Unlike (*34*) and other studies that use transmitted *de novo* mutations to measure mutation rates(*35–38*), these estimates reflect mutation rates in germ cells in the testis and thus offer insight on a novel perspective of germ cell mutagenesis.

#### Blood is a poor surrogate for measuring mosaicism of gonosomal PZMs

Motivated by the fact that only a small subset of tissue types is easily and ethically accessible in antemortem human subjects research, we hypothesized that more accessible tissues may be useful surrogates for examining prenatal PZMs in less accessible tissues. The results of such analyses may shed light on the cellular dynamics of human development and implications for preconception genetic counseling and *de novo* mutation discovery.

We fit a mixed-effects model to predict whether a gonosomal PZM was detected in a somatic tissue while controlling for technical effects. Surprisingly, 88% (38/43) of tissues had significantly higher odds of detecting gonosomal PZMs than in blood (**Fig. 4D**), suggesting that blood is a poor surrogate for detecting gonosomal PZMs. Additionally, 88% (38/43) of somatic tissues had a significant linear correlation between the somatic VAF and the germ cell VAF (**Fig. 4E** and **F**; Pearson’s correlation test), suggesting that somatic tissues may offer a faithful representation of gonosomal PZMs in germ cells. These observations were not an artificial result of germline variant filtering or our cross-sample mutation calling strategy (**Supplementary Result 2.10.4**, **fig. S26**).

Based on these results, for trio studies, we recommend sperm (a direct readout of germ cells) should be profiled in males and skin (which is predicted to be over 5× more likely than blood to contain a gonosomal PZM) should be profiled in females.

### Conclusion

Here, we present one of the most comprehensive and diverse surveys of PZM variation in normal individuals, which should prove a valuable resource for understanding the causes and consequences of PZMs across the body. By linking these mutation calls to the vast data and tissue resources of the GTEx project, there are a number of new analyses that could be attempted. First, our observations are consistent with a heritable component to PZM burden, which, if it exists, could be mapped using GWAS(*39, 40*). Second, the impact of PZMs on gene expression traits, both in *cis* and *trans,* can be directly assessed(*10, 41*). Third, the spatial and cell-type distribution of the mutations reported here could be mapped in banked tissue samples from the GTEx donors(*7, 42*), and the mutation type and burden of each sample associated with histology images collected by the GTEx project. We performed extensive validation of our PZM callset, and these validation data will be helpful in training new algorithms for PZM detection.

We observed a number of striking features regarding the developmental origins of mutations that deserve follow-up. Most intriguing is a class of low VAF prenatal mutations that appear to have the highest fraction of deleterious mutation across the human lifespan, even considering disease states. This observation, based on a definition of deleteriousness on an evolutionary timescale, suggests that the functional consequences of mutation can have opposite fitness effects at different stages of the life cycle of genomes and in different cellular contexts. One well established example of dramatic differences in fitness effects between somatic and germline cells is the RAS-MAP pathway, in which gain-of-function mutations provide a transmission advantage to male germ cells, but are often reproductively lethal for the resulting conceptus(*43, 44*). While some parallels have been noted between molecular mechanisms of carcinogenesis and normal embryogenesis(*45, 46*), there is essentially no data on the potential adaptive effects of PZMs on embryonic or fetal development in healthy individuals.

We reported a large difference in dN/dS inferred from PZMs and inherited germline variants, consistent with strong purifying selection reducing the transmission of deleterious mutation across generations (**Figs. 3E** and **4B**). An important future direction is to dissect and quantify the physiological basis of this purifying selection (**Fig. 4G**). With careful thought and experimental design, it should be possible to model the steps of the human lifecycle where purifying selection can occur, estimate the strength of selection at each step, and translate this data into life stagespecific measures of selective constraint for each gene in the genome. This would be of great benefit to human geneticists, who rely heavily on selective-constraint measures aggregated across the lifecycle (such as CADD) for interpretation of genetic variants in the context of disease(*47, 48*). Stage-specific constraint metrics could augment current methods for variant interpretation to be more relevant to the tissue and developmental time affected by a disease.

## Acknowledgements

We would like to thank the GTEx donors and families for their generous and selfless donation of tissues and organs for the advancement of science. We thank T. Lappalainen, the M. Griffiths lab, S. Montgomery lab, the Conrad lab and the Cohen lab for data sharing and helpful discussions. Sequencing for the validation experiments was performed by the Genome Technology Access Center (GTAC) in the Department of Genetics at Washington University School of Medicine.

We would also like to acknowledge that this manuscript includes several analyses that focus on GTEx donors with European and African American ancestry and less so on other ancestries in GTEx. This decision was made so that we’d have higher statistical power to detect potentially rare genetic signals that may vary across populations. We hope that in future studies, larger numbers of Black, Indigenous, and People of Color are included so that we can learn about all populations in an equitable manner.

## Funding

National Institutes of Health ***grant*** R01MH101810, R01HG007178, and R01HD078641 (DFC)

National Human Genome Research Institute grant T32HG000045 (NBR)

Medical Research Council MR/M004422/1 and MR/R023131/1 (KSS)

Wellcome Trust

Medical Research Council

European Union

Chronic Disease Research Foundation (CDRF)

the National Institute for Health Research (NIHR)-funded BioResource Clinical Research Facility and Biomedical Research Centre based at Guy’s and St Thomas’ NHS Foundation Trust in partnership with King’s College London.

## Authors contributions

D.F.C. conceived the study. N.B.R. designed and performed research and analyzed data. W.H.W. and E.Z.T. assisted with experimental design, sample procurement and generated sequencing libraries for validation experiments. M.J.N. and L.N. performed and analyzed data from the sperm experiment. B.M. and A.Z. contributed to analyses. A.R., C.D., and N.H. contributed to the study design. K.A.V. contributed to data visualization. D.F.C, K.A, and B.A.C. supervised the project. N.B.R. and D.F.C. wrote the manuscript in consultation with all authors.

## Data and materials availability

All GTEx protected data are available through the database of Genotypes and Phenotypes (dbGaP) (accession no. phs000424.v8). Access to the raw sequence data is now provided through the AnVIL platform (https://gtexportal.org/home/protectedDataAccess).

The cancer results are based upon data generated by the TCGA Research Network: https://www.cancer.gov/tcga. We appreciate obtaining access to de novo mutations on SFARI base (https://base.sfari.org).

