## Supplementary Information for "The origins and functional effects of postzygotic mutations throughout the human lifespan"

1.7.4 Simulated random mutations ... 46
1.8 PZM deleteriousness ... 47
1.8.1 PZM deleteriousness validation ... 49
1.9 Selection ... 52
1.9.1 Definition of cancer genes ... 54
1.10 Germ cell PZM analyses ... 54
1.10.1 PZM variant filtering ... 54
1.10.2 Sperm sequencing experiments ... 55
1.11 Calculation of mutation rates during gametogenesis ... 58
1.11.1 Germline surrogate tissue analysis ... 59
**2 Supplementary results ... 61**
2.1 PZMs in hypermutated samples are likely false positives ... 62
2.2 PZM features recapitulate known biology ... 63
2.3 Preservation method does not likely affect mutation burden and spectra ... 65
2.4 Evidence suggesting multi-tissue PZMs occurred prenatally ... 69
2.5 PZM phylogeny reconstruction power ... 73
2.6 PZM phylogeny validation ... 75
2.6.1 Mapped prenatal PZMs are explained by shared developmental history rather than random sharing patterns ... 75
2.6.2 Observed mutation edge weights are independent of edge mapping power ... 77
2.7 Relationship between VAF and deleteriousness score ... 79
2.8 The high fraction of deleterious PZM mutations is recapitulated in multiple validation datasets ... 80
2.8.1 Brazhnik et al. validation dataset ... 80
2.8.2 TwinsUK validation dataset ... 82
2.8.3 Yizhak et al. validation dataset ... 84
2.8.4 García-Nieto et al. validation dataset ... 85
2.9 Prevalence of expressed CHIP mutations in GTEx ... 87
2.9.1 Definition of CHIP mutations ... 88
2.9.2 Expressed CHIP mutations are observed at expected rates ... 88

### 1 Methods

#### *1.1 Algorithm for calling PZMs (LachesisDetect)*

LachesisDetect contains four basic steps. First, alignment files are filtered for extremely high quality alignments. Next, the algorithm leverages cohort-wide information by simultaneously analyzing all samples to estimate position-specific error models for over 115 Mb of the transcriptome. LachesisDetect uses these models to detect putative PZMs with single-sample calling. Third, the method removes sources of false positive PZMs such as RNA editing and allele-specific expression of germline variants using > 15 filters based on theoretical and experimental validation metrics. In the last step, the method leverages donor information by jointly analyzing all samples in a donor to detect mutations with low power and estimate empirical false positive rates (**Fig. S1**). As LachesisDetect uses statistical frameworks for many of the steps, the user has precise control on how conservative to make the mutation calling.

##### *1.1.1 Motivation and overview of method*

While there are a multitude of variant calling algorithms, at the time of starting this project, there was no appropriate method to detect DNA PZMs from bulk RNA-seq data. Therefore, we developed a novel method called LachesisDetect to identify DNA postzygotic point mutations from bulk RNA-seq data. As mutations can cause disease and eventually mortality, we named the algorithm after the Greek goddess, Lachesis, who measures the thread of life and chooses a person's destiny. While we designed LachesisDetect with GTEx data in mind, the method is generalizable to other medium-to-large cohort studies.

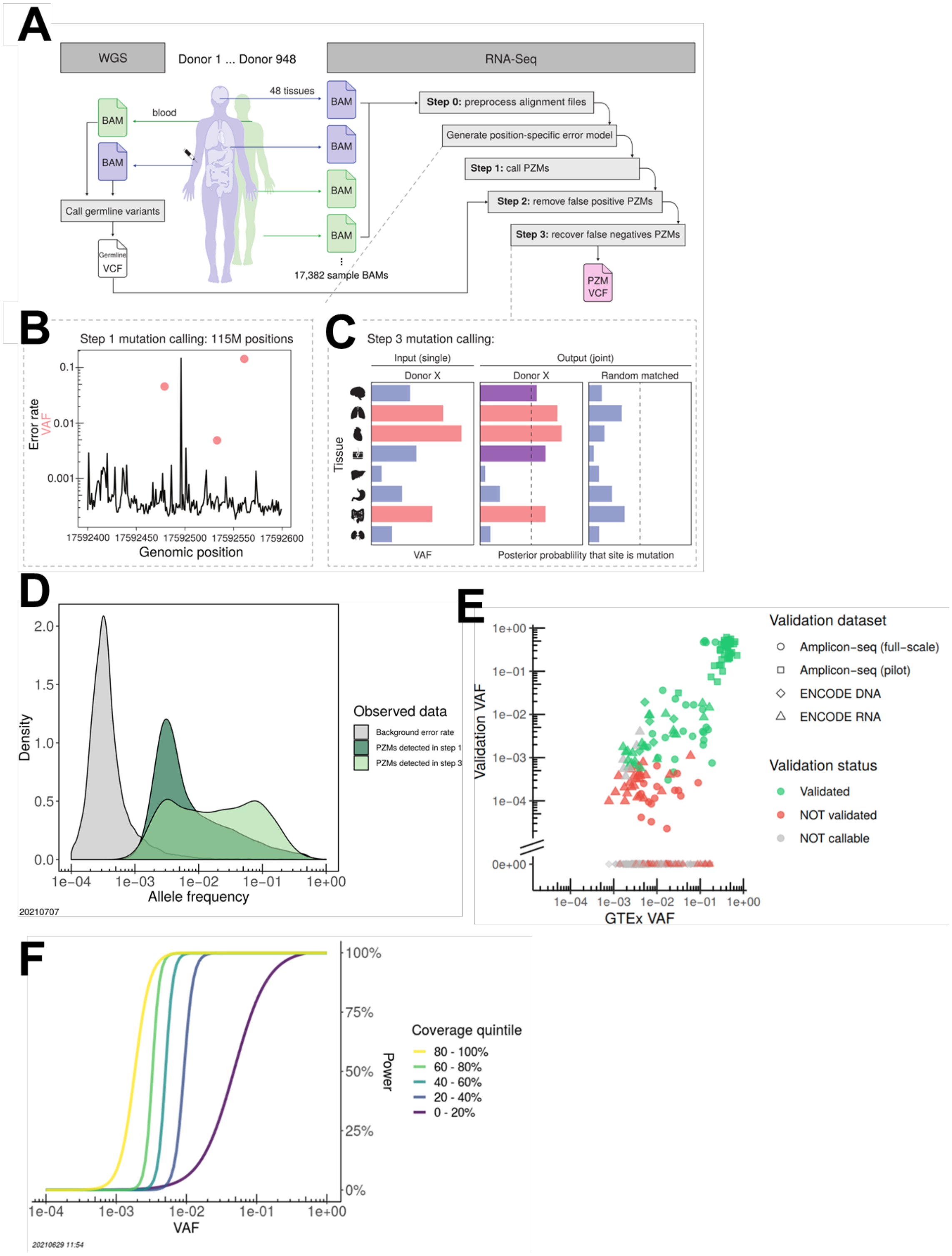

145 **Fig. S1. Overview of study design and mutation calling method.** (**A**) The input to LachesisDetect consists of WGS and RNA-seq BAMs and the output is a VCF of the putative

PZMs. (**B**) Overview of step 2. An error model is generated at each position in the transcriptome. PZMs are detectable when the VAF is greater than the error rate at a position. Error rate versus genomic position for a representative region on chr22. Three PZMs were detected in this region (marked with coral dots). Error rate is defined as the fraction of non-reference allele coverage versus total coverage across all samples. VAF is defined as the fraction of alternative allele coverage versus total coverage in a sample. Thus error rate and VAF are both unitless and are supported on [0,1]. A PZM is detectable when the VAF is significantly greater than the error rate at a given position. (**C**) Overview of step 3. Variant allele information for all samples in a given donor are used as input, including tissues with the PZM detected (red) and tissues with no PZM detected (blue) (**left**). Step 3 estimates the posterior probability that each donor tissue contains the PZM (**middle**). Tissues that did not previously have the mutation detected may have been false negatives (purple) that are now detected. Random tissue-matched controls are used for modelling and model evaluation (**right**). Data is from simulation. (**D**) Distribution of position-specific error rates at PZMs and distributions of PZM VAFs that were identified after step 1 (true positives) and step 3 (false negatives).For the former VAF distribution, only the subset PZMs that were retained after the completion of the pipeline were plotted, i.e., after false positive PZMs were removed in step 2. (**E**) Validation VAF versus GTEx RNA-seq VAF for PZMs where validation was attempted. (**F**) Step 1 PZM detection power as a function of VAF and RNA-seq coverage quintile. BAM = binary sequence alignment/map. PZM = postzygotic mutation. VAF = variant allele frequency. VCF = variant call format. WGS = whole-genome sequencing.

##### *1.1.2 Basic tenets of the method*

An RNA-seq alignment containing a non-reference allele is the result of one of two mutually exclusive processes: 1) a postzygotic mutation event (a true positive) or 2) a non-postzygotic mutation event (a false positive). False positives can be due to 1) an experimental error, 2) a germline variant (with or without allele-specific expression), or 3) an RNA-editing event. When viewed as a whole, the fraction of non-reference reads to all reads at a position represents the signal-to-noise ratio and is defined as the variant allele frequency.

In a bulk RNA-seq sample, the fraction of cells containing the PZM can be very small and therefore, the VAF can also be very small. To detect variants accurately, the signal must be greater than the noise. Of the false positive processes listed above, experimental error can be the most challenging to discriminate against because the PZM VAF and experimental error distributions can overlap considerably. In this regime, the PZM VAF is very low (e.g., VAF <

1%) and the error rate is high (error rate < 1%). Since many PZMs are rare in bulk tissue, we chose to focus our efforts in detecting rare PZMs and discriminating putative mutations from noise. Specifically, we first scanned the transcriptome for positions where the VAF was significantly larger than the experimental error rate, thereby removing false positives due to experimental error. We then ruled out false positives due to germline variants and RNA-editing events with conservative filtering. By eliminating all false positive processes, we are left to conclude that the position is a putative PZM.

###### *1.1.3 Step 0: Preprocess alignment files*

To minimize the number of false positive variant calls, we only included the highest quality data. For each sample's alignment file, we excluded non-uniquely mapped alignments, duplicate alignments, vendor failed alignments, alignments with > 20 soft-clipped bases, alignments with soft-clipping on both ends, alignments with > 2 mismatches, and alignment read pairs with > 3 mismatches. Additionally, for each base within the sequencing read, we excluded those with base qualities < 20. If read 1 and read 2 overlapped, the overlapping bases with the lower base qualities were excluded to avoid double-counting the same cDNA molecule in the coverage calculations. These thresholds were determined by evaluating the reproducibility of mutation call sets made from two different versions of the data (GTEx v6 and GTEx v8). The preprocessing pipeline used custom scripts, SAMtools v1.9(*1*), and a modified version of samclip (https://github.com/tseemann/samclip).

###### *1.1.4 Step 1: Identify putative PZMs*

There are several sources of experimental error that can contribute to a non-zero VAF: nucleic acid damage during sample preparation (e.g., from reactive oxygen species(*2*)), amplification errors during sample preparation and sequencing, and alignment errors(*3*)). These sources of error can be modeled *en masse* as a binomial random variable for each sample $s$ and position $i$ (*4*) as, $E_{s,i} \sim Binom(n_{s,i}, e_i)$, where $n_{s,i}$ (the number of trials) is the total coverage in sample $s$ at position $i$ and $e_i$ (the probability of success, i.e., an experimental error occurred) is the experimental error rate at site $i$.

Since the GTEx dataset contains thousands of samples, we hypothesized that using information from all samples would lead to a more accurate estimate of experimental error rate than what could be obtained from a single sample. We defined the experimental error rate at site $i$, $e_{s,i}$, as the ratio of non-reference coverage across all samples excluding sample $s$ to the total coverage across all samples excluding sample $s$ at position $i$. To maintain a high level of conservatism, we included all GTEx v8 RNA-seq samples, including those that did not pass quality control, in the background error model (N = 17,382). In total, we generated a position-specific error model for over 115 Mb of the transcriptome (**Fig. S1.B**).

For a given sample $s$ and position $i$, we defined the putative PZM alternative allele as the non-reference allele with the highest coverage and then tested if the alternative allele coverage was significantly higher than expected given $e_{s,i}$ (binomial test) (**Fig. S1.B**). *P*-values were corrected using the Benjamini-Hochberg procedure. Putative PZMs were defined as sites with $q \leq 0.05$. To reduce the number of hypothesis tests (and therefore increase our detection power), we corrected the *P*-values after several filters were applied (see Step 2: Remove false positive PZMs).

Of note, this approach assumes the PZM is rare in the population of samples (i.e., the non-reference coverage is assumed to primarily come from false positive events rather than PZM events). This assumption is appropriate because PZMs are expected to be rare events(*5*) and our sample set is large (17,382 samples across 948 donors).

This framework could be extended to model arbitrary types of correlation structures that might exist in the error processes, e.g., batch effects across sequencing runs and base- and strand-specific error models(*6*). Given the algorithm's reasonably high validation rate (see PZM validation), we concluded the relatively simple model was sufficient to accurately detect PZMs.

###### *1.1.5 Step 2: Remove false positive PZMs*

In the next step, false positive PZMs are removed using sample-, position-, and variant-specific filters. These filters are described in detail below. A list of all the filters applied and how many variants they removed is in **S1**.

###### 1.1.5.1 Remove low quality samples

Using GTEx recommendations, we removed any sample with a RNA integrity number (RIN) < 6 (N = 1,674 samples)(*7*) or was from a tissue type whose collection was discontinued during the GTEx pilot phase due to quality issues (Bladder, Cervix - Ectocervix, Cervix - Endocervix, Fallopian Tube, Kidney - Medulla, and Spleen) (N = 230 samples)(*7*, *8*). The remaining tissues are referred to as the **GTEx pass quality control (QC) tissues**. Additionally, we checked that no tissue sample was from a transplanted tissue/organ using the donor metadata. There were 15,478 tissue samples after filtering (**S2**).

###### 1.1.5.2 Remove genomic regions with potential mappability issues

Genomic regions prone to incorrect read alignments could lead to false positive variant calls. To minimize this issue, we removed several genomic regions. Immunoglobulin genes were removed since they are structurally diverse and polymorphic due to V(D)J recombination, class-switch recombination, and somatic hypermutation. Regions within 10bp of an exon boundary were removed since splice junctions can have inaccurate alignments(*9*)). Segmental duplications and repetitive sequences (annotations downloaded from the UCSC genome browser, http://genome.ucsc.edu/) were also removed since reads may map to an incorrect copy in the genome.

###### 1.1.5.3 Remove RNA editing sites

In RNA editing, the nucleotide sequence of an RNA is modified post- or co-transcriptionally. The most common type of RNA editing in mammals is the deamination of adenosine into inosine (A>I) by the ADAR family of enzymes(*10*). After library preparation and sequencing, the original editing event will be read as an A>G change. Another type of editing, although much rarer, is the conversion of cytosine to uracil via the family of cytidine deaminases (AID/APOBEC)(*11*).

To minimize the number of false positives from RNA editing events, we removed all genomic sites known to harbor RNA editing in humans using version 2 of the Rigorously Annotated Database of A-to-I RNA editing (RADAR) (http://rnaedit.com), a curated and comprehensive source of RNA-editing sites in humans(*12*). The RADAR database was converted from hg19 to GRCh38 coordinates using the liftover utility(*13*). Genomic sites that were not A or T in GRCh38 were excluded.

Additionally, since databases can be incomplete, we conservatively removed any putative A>G PZM on the coding strand that was seen in at least two donors. Our approach will most likely miss rare, donor-specific RNA editing events, to the extent such events exist. Since most sites have a low level of RNA editing (<1%)(*14*), we don't expect RNA editing events to be a major source of error for higher VAF PZMs.

###### 1.1.5.4 Remove germline variants

###### 1.1.5.4.1 Known germline variants filter

To remove false positive PZMs due to germline variants, we removed all genomic sites with a germline SNV or within 10bp of a germline indel in *any* of the donors' whole blood genotyping datasets. To increase our sensitivity of removing germline variants, we used all available genotyping datasets: v8 WGS data (838 donors), v6 WES (520 donors and 11 GTEx flagged donors), v6 WGS (148 donors), and v6 microarray data (191 donors for the pilot microarray and 275 donors from the midpoint microarray). v6 datasets were converted from hg19 to GRCh38 using the liftover utility(*13*). In our effort to be conservative, our variant calling method will not be able to call PZMs at sites that are polymorphic in the cohort.

Of note, only 853 of the 946 donors had any genotyping data. To minimize the number of false positive PZMs due to both rare germline variants in non-genotyped donors as well as missed germline variants in the genotyped donors, we used two additional germline filters which are described below.

###### 1.1.5.4.2 Putative germline variants filter

This filter removes all putative PZMs that appear to be a putative germline variant in at least one donor. To test if a putative PZM is actually a missed germline variant, we first calculated the cumulative alternative and cumulative total coverage in the donor with $S$ total samples at position $i$ using the following equations:

$$donor\ cumulative\ alt\ coverage_i = \sum_{s}^{S} alt\ coverage_{i,s} \tag{1-1}$$

$$donor\ cumulative\ total\ coverage_i = \sum_{s}^{S} alt\ coverage_{i,s} + ref\ coverage_{i,s} \tag{1-2}$$

In the null model (variant is a heterozygous germline variant), the cumulative alternative read coverage at position $i$ follows a binomial distribution with parameters $N$ (the donor cumulative total coverage), and $p$ (the expected VAF of true heterozygous germline variants in RNA-seq data). In the alternative model (variant is a PZM), the cumulative alternative coverage is less than expected of true heterozygous germline variants. The $p$ parameter was estimated from the RNA-seq data of all high-confidence heterozygous germline SNVs from 838 donors using all of the donors' GTEx pass QC tissue samples ($p = 0.45$). High-confidence heterozygous germline SNVs were defined as SNVs found in at least 1% of donors in the GTEx germline variant VCF.

For each donor variant, we tested if we could reject the null hypothesis using a binomial test. These *P*-values (denoted as **donor germline *P*-values**) were reused in the next filter. In an effort to be conserative, at each position, the largest *P*-value across all donors was retained. This intermediate list of *P*-values was corrected for multiple tests using the Benjamini-Hochberg procedure with a cutoff of q-value ≤ 0.05. Putative PZMs that overlapped a genomic site with a nonsignificant q-value were removed. Of note, this approach will also filter out homozygous germline variants because if the donor cumulative VAF is less than $p = 0.45$, it will also be less than $p \approx 1$, the expected VAF of homozygous germline variants.

###### 1.1.5.4.3 Allowable transcriptome definition

Of note, we defined the **allowable transcriptome** as the transcriptome after applying the genomic filters above. Ancillary datasets were filtered to the allowable transcriptome in order to make fair comparisons with the PZM dataset.

###### 1.1.5.4.4 Match donor mutation burden in genotyped and non-genotyped donors filter

We assumed the true mutation burden in a donor was independent of whether the donor was genotyped. However, prior to applying this filter, non-genotyped donors had higher mutation burdens than genotyped donors suggesting they may have an excess of false positive germline variants. This second germline variant filter specifically corrects for the fact that not all donors were genotyped.

We defined the donor mutation burden in donor $i$ ($d_{i,\alpha}$) as the total number of putative PZMs in all GTEx pass QC tissues from donor $i$ at a donor germline *P*-value cutoff of $\alpha$. $d_{i,\alpha}$ is a mixture of true PZMs and false positives, including false positive germline variants. Of critical note, $d_{i,\alpha}$ can be decreased/increased by using a more/less restrictive *P*-value cutoff ($\alpha$) on the donor germline *P*-values. (Recall that the donor germline *P*-values were calculated in the previous germline variants filter.)

We modeled the donor mutation burden as

$$\sqrt{d_{i,\alpha}} = \beta_{0,\alpha} + \beta_{1,\alpha} * N_i + \beta_{2,\alpha} * AGE_i + \beta_{3,\alpha} * self\_reported\_ancestry_i + \beta_{4,\alpha} * genotype\_status_i + e_{i,\alpha} \quad \text{1-3}$$

where $N_i$ is the number of GTEx pass QC tissues in donor $i$; $AGE_i$ is the age of donor $i$; $self\_reported\_ancestry_i$ is the self-reported ancestry of donor $i$; $genotype\_status_i$ is the genotyping status of donor $i$ (genotyped or not genotyped); $\beta_{\mathrm{i},\alpha}$'s are model coefficients at $P$-value cutoff $\alpha$; and $e_{i,\alpha}$ is the residual error. The square root of the donor mutation burden distribution was applied to transform residuals to a normal distribution.

We performed the linear regression on a large range of $P$-value cutoffs ($\alpha$) and selected the cutoff that corresponded to the least significant genotype status regression coefficient ($\beta_{4,\alpha}$). As expected, $AGE$ was significant across the germline variant significance cutoffs. The log10 $P$-value of $genotype\_status$ had a global maximum, suggesting cutoffs that are too small or too large could lead to over- or undercalling variants, respectively. The final donor germline $P$-value cutoff was $\alpha =$ 1E-30, corresponding to a non-significant $P$-value = 0.98 for the $\beta_4$ coefficient (**Fig. S2**). Thus, in the final model, donor mutation burden was independent of genotype status.

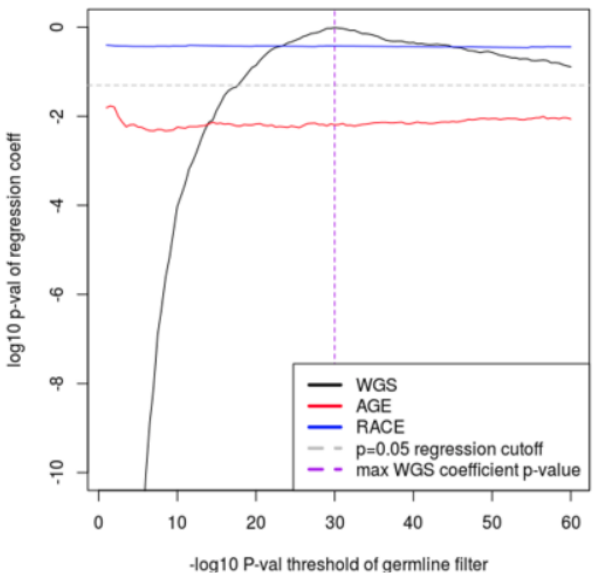

**Fig. S2. Selection of the appropriate donor germline *P*-value cutoff, $\alpha$.** $\alpha$ was chosen such that the $P$-value of $genotype\_status$ was minimized (log10 $P$-value maximized). The maximal log10 $P$-value of $genotype\_status$ is marked by dashed purple line $x =$ -$\log_{10}$ 1E-30. At this cutoff, $genotype\_status$ was not significantly associated with donor mutation burden (value above gray dashed line at $y = \log_{10} 0.05 = -1.3$). The effect of $\alpha$ on the significance of other covariates in the model is also shown.

###### 1.1.5.5 Remove variants with low coverage

We used the standard strategy in variant calling to remove mutations with low sequencing coverage. We removed mutations with ≤ 20× total coverage and ≤ 5× alternative coverage. In step 3 (below), this alternative coverage constraint was dropped, since less alternative coverage is more tolerable when there is strong evidence of a mutation in a different tissue from the same donor.

###### 1.1.5.6 Remove variants at genomic positions with high nucleotide diversity

Since we are calling mutations at non-germline variant sites, a genomic position with coverage for more than two alleles in a sample is an indication of experimental and/or bioinformatic error. Therefore, we removed mutations at genomic positions where there was at least 3× coverage for more than two alleles in a sample. This conservative filter should remove problematic genomic positions not caught by upstream filters.

###### 1.1.5.7 Remove variants with biased alignment metrics

When manually inspecting alignment files in a genome browser for false positive variants, a common heuristic to look for is whether the putative variant is supported by “diverse” alignments, e.g., alignments that map to both strands of the genome and alignments that contain the mutant allele at different positions relative to the start of the sequencing read. Since the RNA-seq protocol used to generate the data was unstranded, we conservatively removed variants that did not have at least 1× coverage on each of the Watson and Crick strands. Additionally, we removed variants that were only supported by alignments where the alternative allele was found near (within 25 bases) the start or end of the sequencing read.

###### 1.1.5.8 Remove hypermutated samples

We conservatively defined a sample as **hypermutated** if its mutation burden was greater than $Q_3 + 5 * IQR$, where $Q_3$is the third quartile and $IQR$ is the interquartile range of the sample mutation burden distribution (Tukey's fence method for outlier detection). This corresponded to a cutoff of 63 mutations and labeled 806 (5.2%) samples as hypermutated. Due to their extreme mutation burden, we removed these samples from the downstream analyses.

We used a variety of experimental validation methods to ascertain if this hypermutation phenomenon was biological or a technical artifact (see "PZMs in hypermutated samples are likely false positives"). Based on the low positive predictive value (PPV) in the hypermutated samples, the observed hypermutation is most likely a technical artifact and thus removal of these samples is well justified.

###### 1.1.5.9 Remove variants at sites with large background error rates

We observed a higher false discovery rate when the background error rate was high in all four independent sequencing-based validation datasets (**Fig. S3**) and *in silico* read-based phasing validation. (See PZM Validation for a description of the validation methods.) To improve the accuracy of the mutation call set, we calculated the maximum background error rate that maximized the validation rate of the four validation datasets (background error rate cutoff = 6E-4). Putative PZMs at positions with a background error rate greater than this cutoff were removed. In total, 13% (9.8 Mb) of the allowable transcriptome was removed by this filter.

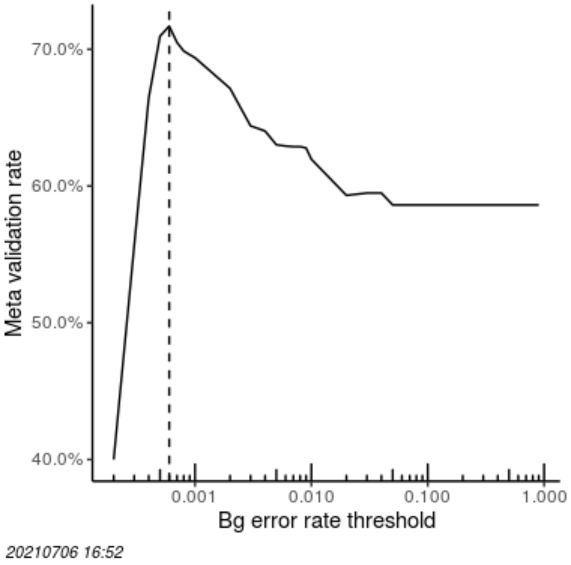

**Fig. S3. Validation rate as a function of background error filter threshold.** The threshold was chosen to maximize the validation rate (annotated as a dashed vertical line at y = 6E-4). Meta validation rate defined as the average positive predictive value across all validation datasets. Bg = background.

##### *1.1.6 Step 3: recover false negative PZMs with EM*

Step 1 may miss some PZMs, e.g., when there is low total coverage and/or low alternative allele coverage. Therefore, we implemented a more sensitive mutation calling strategy to identify false negative variants for the third step of the algorithm. The power to detect false negative mutations comes from jointly analyzing data from all tissue samples in a donor.

PZMs that were detected in step 1 of the algorithm have appreciable VAFs and thus represent either 1) a clone that rose to high frequency in a tissue or 2) a PZM that occurred early in development. In the later case, depending on the timing of the mutation, the mutation may exist in multiple tissues. Thus, if a PZM is found at position $i$ in tissue $t$ of donor $d$ (step 1), there is an increased likelihood that the PZM also exists in tissue $\tau$ of donor $d$. We refer to these PZMs as **multi-tissue PZMs**. Multi-donor PZMs are specified by a unique combination of genomic coordinates, alternative allele, and donor ID.

We modeled the distribution of alternative allele read coverage from each tissue in a donor at position $i$ as a mixture of two binomial distributions: one where the alternative reads are generated from a true PZM and the other where the alternative reads are generated from experimental errors. We used expectation maximization (EM) to calculate for each tissue, the posterior probability that the tissue contains the multi-tissue PZM. This process is described in more detail below. PZMs in tissues with a posterior probability of at least 0.95 were added to the mutation call set. PZMs in tissues that failed to meet this constraint were removed. Of note, this included removing a small fraction (1.5%) of PZMs that were originally detected in step 1 (**Fig. S1C**, **Table S1**).

Specifically, we parameterized the multi-tissue PZM and error distributions as $Binom(n_{i,s}, VAF_i)$ and $Binom(n_{i,s}, e_i)$, respectively, where $n_{i,s}$ is the total coverage of tissue $s$ at position $i$, $VAF_i$ is the (latent) PZM VAF at position $i$, and $e_i$ is the (latent) error rate at position $i$. To avoid local minima in the likelihood space, we used multiple initializations and selected the model with the highest likelihood. Additionally, to ensure that the set of tissues was indeed a mixture of both tissues with and without the variant, a set of matched tissues randomly selected from the remaining donors was added to the modeling (denoted as **negative samples**). Since donor samples were required to have at least 10× coverage, the same constraint was also used for the negative samples. For each tissue, 20 attempts were made to find a matched negative sample with sufficient coverage. Matched tissues were also used to control for tissue-specific issues, e.g., alignment errors due to tissue-specific isoforms.

###### 1.1.6.1 Empirical FPR and FDR estimates

The matched negative tissues were also used to estimate the FPR (false positive rate) and FDR (false discovery rate) of the mutation call set. FPR was defined as:

$$FPR = \frac{\sum_{d=1}^{D} \#\ of\ neg.samples\ with\ posterior\ prob. \geq p\ in\ multi\text{-}tissue\ PZM_d}{\sum_{d=1}^{D} \#\ of\ neg.samples\ in\ multi\text{-}tissue\ PZM_d} \quad \text{1-4}$$

where $D$ is the total number of multi-tissue PZMs and $p$ is posterior probability cutoff (set to 0.95).

FDR was defined as:

$$FDR = \frac{\sum_{d=1}^{D} Estimated\ \#\ of\ FPs\ in\ donor\ samples\ in\ multi\text{-}tissue\ PZM_d}{\sum_{d=1}^{D} \#\ of\ donor\ samples\ with\ posterior\ prob. \geq p\ in\ multi\text{-}tissue\ PZM_d} \quad \text{1-5}$$

where FPs = false positives.

The $Estimated\ \#\ of\ FPs\ in\ donor\ samples\ in\ multi\text{-}tissue\ PZM_d$ was defined as

$$Estimated\ \#\ of\ FPs\ in\ donor\ samples\ in\ multi\text{-}tissue\ PZM_d = \frac{\#\ of\ neg.samples\ with\ posterior\ prob. \geq\ p\ in\ multi\text{-}tissue\ PZM_d}{\#\ of\ neg.samples\ in\ multi\text{-}tissue\ PZM_d} * \#\ \text{of donor samples} \quad \text{1-6}$$

##### *1.1.7 Step 1 PZM detection power*

The power to detect a PZM in step 1 depends on the VAF, coverage, and background error rate. We estimated the power to detect a PZM mutation as a function of these variables through simulation.

For each simulated PZM, we randomly selected a tissue sample, genomic position, and VAF. Using the selected sample's coverage at that position (denoted as $N$), we simulated the alternative allele read coverage as $\lfloor N * VAF \rfloor$ and calculated if the simulated PZM would be detected given the alternative allele read coverage, $N$, and the background error rate at that position (binomial test). This process was repeated 240,000 times for each chromosome and preselected VAF (37 VAFs in the range [1E-4, 1].) Only male samples were used for simulated mutations on chrY. Mutations on chrX were originally simulated for males and females separately; however no sex-difference in power was detected, so the chrX data was combined. Since the total coverage is not known *a priori,* after all simulations were run, we split the simulations into quantiles of sample coverage bins.

**PZM mutation detection power** was defined as the fraction of simulations where the simulated mutation was correctly recovered at a given VAF and coverage bin. Power curves were fit to sigmoid functions using the Levenberg-Marquardt nonlinear least-squares algorithm from the minpack.lm R package (v 1.2-1) (https://CRAN.R-project.org/package=minpack.lm).

To estimate the average PZM mutation detection power in a given sample $s$ from tissue $t$ (denoted as $\boldsymbol{sample\_mutation\_detection\_power}$), we first estimated the joint distribution of VAF and coverage of all detected PZMs in tissue $t$ (denoted as the **tissue-VAF-coverage distribution**). (Here, we assumed that all samples in a given tissue have the same tissue-VAF-coverage distribution. This assumption allowed us to more accurately estimate the distribution due to the larger sample size.) Next, we estimated the distribution of coverage in the transcriptome in sample $s$ (denoted as the **sample-coverage distribution**). We simulated 5000 PZMs for each sample. For each simulated PZM, we randomly sampled a coverage bin from the sample's sample-coverage distribution and then randomly sampled a VAF from the tissue-VAF-coverage distribution corresponding to that coverage bin. We estimated the power to detect the PZM by using the PZM detection power curves for that coverage bin and VAF.

$sample\_mutation\_detection\_power$ in sample $s$ was defined as the average power of all simulated PZMs in that sample. Nine quantiles were used to define the coverage bins.

##### ***1.2 GTEx data***

We detected variants in the GTEx v8 dataset. To achieve a high quality dataset, we removed RNA-seq samples with RIN < 6, were derived from tissues with overall poor quality(*8*) or had an extremely high PZM mutation burden (see Remove hypermutated samples; **Table S2**). We also confirmed that none of the analyzed samples were from transplanted tissue. After our quality control, there were 14,672 samples from 944 donors from 48 diverse tissue and cell types. Library preparation, sequencing, alignment, and GTEx quality control are described in detail in (*15*). Briefly, RNA-seq samples were prepared using a polyA, unstranded RNA-seq protocol (Illumina TruSeq) and sequenced on Illumina HiSeq instruments (HiSeq 2000 or HiSeq 2500) with 76bp paired-end reads to a median coverage of ~83M total reads. Reads were aligned to the GRCh38 human reference genome using STAR v2.5.3a(*16*) and GENCODE v26 gene models. Duplicate reads were marked with Picard Tool's MarkDuplicates (v2.9.4) (https://broadinstitute.github.io/picard/).

###### *1.2.1 PZM variant calling*

We ran LachesisDetect on the GTEx RNA-seq BAMs to generate a list of putative PZMs in GTEx.

###### *1.2.2 Germline variant calling*

To increase the sensitivity to detect germline variants, we used all genotyping GTEx datasets that were available at the time of writing: v8 WGS data (838 donors)(*15*), v6 WES (520 donors and

11 GTEx flagged donors), v6 WGS (148 donors), and v6 microarray data (191 donors for the pilot microarray and 275 donors from the midpoint microarray)(*8*). Detailed descriptions of the sample preparation, sequencing, alignment, and quality control can be found in their respective publications.

Briefly for the v8 WGS data, for the majority of samples, DNA was isolated from whole blood. A non-blood source was used if blood was not available or the blood sample failed quality control. WGS was performed using 101bp or 151bp paired-end reads on Illumina HiSeq instruments (HiSeq 2000 or HiSeq X) to a median depth of 32×. Reads were aligned to the GRCh38 human reference genome using BWA-MEM (http://bio-bwa.sourceforge.net). Germline SNVs and indels were called using the GATK HaplotypeCaller (v3.5)(*17*).

###### *1.2.3 PZM Validation*

We performed several orthogonal validation experiments to quantify the FDR of the mutation calling algorithm. These efforts included both *in silico* and experimental approaches and involved analysis of both DNA and RNA from the tissues used for mutation detection. A summary of the validation results is in **Table S3**.

###### *1.2.4 In silico read-based phasing*

Read-based phasing is an *in silico* technique to estimate the FDR of a variant call set and has been used in somatic mutation validation(*18*, *19*). Under the assumption that PZMs are rare events, it is unlikely that two PZMs will occur at the same position on each of the maternal and paternal copies of the genome. Therefore, if a PZM does *not* segregate/phase with one parental haplotype, it is likely a false positive. This can be determined bioinformatically by analyzing sequencing reads that physically span both the putative PZM and a nearby heterozygous germline variant (**Fig. S4**).

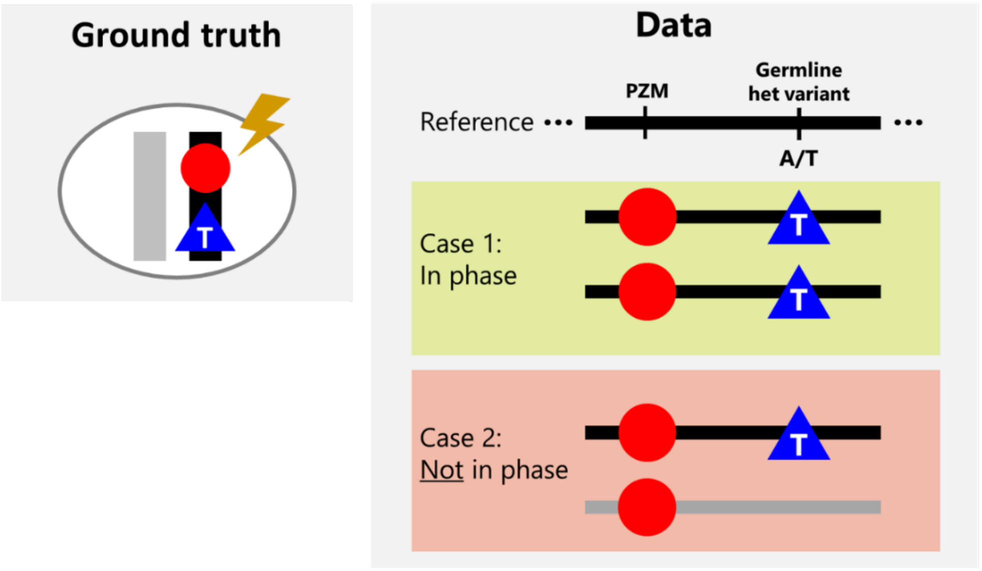

**Fig. S4. Overview of *in silico* read-based phasing.** A red PZM occurs on the black haplotype of the cell's diploid genome. The PZM is near a blue heterozygous germline which is T on the black haplotype. In the sequencing data, reads that contain the PZM and the T allele are in phase. However, reads that contain the PZM and reference allele (on the gray haplotype) are not in phase and likely represent a false positive PZM. Het = heterozygous. PZM = postzygotic mutation.

For each putative PZM, we scanned the corresponding sample's RNA-seq BAM for sequencing reads that overlapped the PZM and a heterozygous germline variant within 1,000,500 bp (the length of the largest human intron (~1 Mb) plus the length of the sequencing read). We defined these reads as **phasable sequencing reads**. We calculated the number of phasable sequencing reads that contained each combination of alleles at the PZM and germline variant loci. We used Fisher's exact test to test if the PZM segregated with one parental haplotype, i.e., whether the PZM variant allele was associated with one of the heterozygous alleles. To reduce noise due to low power, we ignored PZMs with ≤ 10 phasable sequencing reads. *P*-values were corrected using the Benjamini-Hochberg procedure. Using this approach, 41 PZMs had sufficient phasable sequencing reads.

Fisher's exact test is advantageous over simpler threshold-based methods because it uses a statistical framework allowing the researcher to tune significance cutoffs and allows for experimental errors at both the PZM and germline variant loci. Of note, false positive PZMs can still appear to segregate with one haplotype due to misalignments. For example, reads containing a germline variant can look like a phased PZM when aligned to the wrong copy of a segmental duplication(*20*). To minimize this issue, our variant calling method removed sequencing reads that mapped to multiple locations, PZMs that overlapped with repeat masked regions and segmental duplications from the call set (see Step 2: Remove false positive PZMs).

###### *1.2.5 ENCODE DNA and RNA validation*

The ENCODE consortium profiled several tissues from four GTEx donors using a variety of genomic, transcriptomic, and epigenomic assays. We hypothesized that we could use this rich, and importantly, independent dataset to validate our variant calls. The database of available ENCODE BAMs was queried in June 2019 and BAMs were downloaded at the same time.

We excluded DNA-based assays that used bisulfite treatment to avoid the increased complexity of aligning and calling mutations in bisulfite-treated DNA. We also excluded BAMs that failed ENCODE quality control and RNA-derived BAMs with RIN < 6. For a given sample, we excluded BAMs that had zero coverage at all PZMs in the sample. After filtering, 57 GTEx tissue samples were profiled by 245 ENCODE DNA assays and 20 GTEx tissue samples were profiled by 67 ENCODE RNA assays. The ENCODE data also included one hypermutated sample. After filtering, 1 hypermutated GTEx tissue sample was profiled by 7 ENCODE DNA assays and 3 ENCODE RNA assays (**Table S3**). ENCODE RNA and DNA BAMs were filtered using the BAM preprocessing pipeline (see Step 0: Preprocess alignment files).

To increase the power to detect mutations, we combined all BAMs from DNA-based assays that were generated from the same tissue and donor and combined all RNA-based assays that were generated from the same tissue and donor. By combining BAMs, the coverage at a given site may increase, and thus we may have higher power to detect mutations. An additional benefit of combining the data is that by analyzing data produced with many different technical variables (e.g., different assay protocols, different sequencing centers, different aligners, etc.) the noise in the data should increase. Thus, when a putative PZM is supported by the ENCODE data, we can be very confident that the mutation is a true positive and not the result of a systematic artifact that occurred in both the GTEx and ENCODE datasets.

###### 1.2.5.1 Validation method

For each putative PZM in GTEx, we calculated the coverage of each allele in the corresponding combined ENCODE BAM. If the PZM alternative allele had the highest non-reference allele coverage, we then tested if the alternative allele coverage was significantly higher than expected given the GTEx RNA-seq error rate at that position (binomial test) (see Note on the ENCODE validation error model). *P*-values were corrected using the Benjamini-Hochberg procedure. Variants with q-values $< 0.1$ were defined as validated. If the q-value $\geq 0.1$ and the PZM was defined as callable but not validated (see Callability definition). Furthermore, if the site was callable but the alternative allele did not have the highest non-reference allele coverage, the site was not validated.

###### 1.2.5.2 Callability definition

To calculate an accurate FDR, it is important to exclude variants where there is not enough power to detect the mutation in the validation data. We defined such variants as **not callable**. To determine if a variant was callable in the validation data, we simulated the distribution of alternative allele coverages that would be expected given the validation coverage and the putative PZM's VAF (estimated from the original GTEx data). We next calculated the fraction of simulations where we could detect the variant (N = 10,000 simulations). A simulated variant was detected if the simulated alternative allele coverage was significantly higher than expected given the GTEx RNA-seq error rate at that position (binomial test, $\alpha < 0.05$) (see Note on the ENCODE validation error model). If at least 95% of the simulations identified the variant, we concluded the PZM was callable.

###### 1.2.5.3 Note on the ENCODE validation error model

In the ideal case, we would have used the error model generated from the ENCODE data to validate PZMs. However, since the number of GTEx donors and samples profiled in ENCODE was small and the ENCODE datasets sometimes had orders of magnitude lower coverage than the GTEx dataset, we used the GTEx RNA-seq error model instead. Using ENCODE-based error models may lead to overestimated and/or inaccurate error rates. For example, since each sample can contribute a large percentage of the background error rate, a small number of poor quality samples could lead to an overestimate of the error rate. Similarly, regions with low coverage may lead to inaccurate error estimates due to sampling error. In the DNA-based amplicon-seq validation studies where we had many samples and had high coverage, the DNA-based error model and RNA-based error models were very similar; and thus we felt justified to use an error model derived from RNA to validate variants from DNA data.

##### *1.2.6 Pilot validation study*

We generated a pilot mutation call set from an earlier version of the GTEx data (v3 data freeze) using two variant calling methods distinct from the one used for v8 data(*21*). On the basis of the preliminary mutation call set, we designed a validation experiment using amplicon sequencing, and subsequently used these validation data for optimizing the final calling pipeline. Briefly, PCR primers were designed for 317 amplicons, targeting a total of 384 v3 mutations in 365 samples. Three control samples not related to the GTEx project were also included. Libraries were generated for all samples by multiplex PCR using the Fluidigm platform, and the resulting amplicons were pooled and sequenced on 4 lanes of MiSeq using a 2x150 bp protocol to an average read depth of approximately 200× per amplicon. A list of the amplified regions can be found in **Table S4**. In the final mutation call set, we attempted to validate 47 PZMs across 22 non-hypermutated samples and 14 PZMs across 9 hypermutated samples (**Table S3**).

##### *1.2.7 Large-scale validation study*

Due to improvements in our mutation calling algorithm and the large increase in the number of samples from the pilot study to the GTEx v8 data freeze, we performed another round of validation. The purpose of this validation was twofold: 1) to estimate the accuracy of the PZM calling algorithm and call set and 2) to clarify the source of the hypermutated phenotype observed in a small number of GTEx samples. We designed an amplicon-seq library to validate variants from a randomly selected set of donors (Cohort 1) and a random subset of variants from randomly selected donors with hypermutated samples (Cohort 2). GTEx samples were requested from the GTEx Biobank (https://gtexportal.org/home/samplesPage).

A list of the samples selected for validation is in (**Table S3**). A small number of samples were dropped from the initial study design due to sample quality or withdrawal from GTEx. For brevity, only samples that had PZMs in the final call set are reported in the table.

###### 1.2.7.1 AmpliSeq library preparation and sequencing

Primer pairs were designed to validate mutations using Illumina's Design Studio (https://designstudio.illumina.com/). The following parameters were used: Assay Technology = AmpliSeq for Illumina Hotspot; Species = Homo sapiens (grch38.p2); Sample Type = Regular; Max Amplicon Length = 275bp; Max Amplicon Length = 275bp; Stringency = High. Mitochondrial variants were excluded since Design Studio only allows regions in the nuclear genome. A list of the primer pairs is in

**Table S6** and a list of the locations targeted is located in **Table S5**.

The regions of interest were amplified using 100 ng of genomic DNA and an AmpliSeq for Illumina Custom DNA Panel kit. Sample libraries were sequenced by Illumina NovaSeq (S1 Flow Cell) using 2x150 bp reads to a median coverage of ~38,000×. 4/712 (0.6%) amplicons failed (defined as > 75% of samples with 0× coverage.)

Adapters were trimmed from sequencing reads using trimmomatic (v0.38)(*22*) with options PE -baseout. Sequencing reads were aligned to GRCh38 using BWA (v0.7.17)(*23*) with options MEM -t 4. BAMs were filtered using the same preprocessing pipeline in section Step 0: Preprocess alignment files with the following exceptions: duplicate alignments were not removed since the data was from an amplicon-seq library and alignments with alignment scores < 20 were removed.

In total, we attempted to validate 43 PZMs across 15 non-hypermutated samples and 1,628 PZMs across 5 hypermutated samples (**Table S3**).

###### 1.2.7.2 Validation method

Mutations were validated using the same validation method as the ENCODE datasets (see section ENCODE DNA and RNA validation) except that the error model was generated from the GTEx DNA amplicon-seq dataset since this validation study had high coverage and a large number of donors.

###### 1.2.7.3 Sensitivity of method

We designed a dilution experiment to calculate the sensitivity of the validation method across a range of VAFs (0.0005 - 0.01). Two genomic DNA samples, unrelated to GTEx, were genotyped by WES. The dilution samples were created by diluting one of the samples (the **foreground sample**) into the other sample (the **background sample**) at various ratios. The sensitivity of the validation method was estimated at genomic positions where the foreground sample had a germline variant (heterozygous or homozygous) and the background sample had a homozygous reference genotype.

The dilution samples were prepped with the same amplicon-seq kit used for the large-scale validation study, sequenced on the MiSeq platform (2x150bp reads) to a median amplicon ~2,800× coverage, and processed using the same analysis pipeline.

##### *1.3 Mutation burden modelling*

After an initial phase of data exploration and model selection, we fit the following linear model for mutation burden with nine main effects and three interactions:

$$\begin{aligned} sample\_mutation\_burden \sim\ & tissue + AGE + self\_reported\_ancestry\ + sex \\ & + RIN\ + batch + sample\_mutation\_detection\_power \\ & + genotype\_data\ + transcriptome\_size\ +\ age{:}tissue \\ & + self\_reported\_ancestry{:}tissue\ + sex{:}tissue \end{aligned} \tag{1-7}$$

where $batch$ was the batch ID for when the RNA was isolated and extracted from a sample; $sample\_mutation\_detection\_power$ was the average mutation detection power of simulated mutations in the sample; *transcriptome_size* was the number of "callable bases" present in the sequencing library, and $genotype\_data$ was the list of genotyping platforms used to call germline variants in the donor. The R package car (v 3.0-6)(*24*) was used to calculate type II ANOVA tables.

To estimate the variation explained by donors, we augmented the model in equation **1-7** to a mixed effects model by dropping $batch$ as a fixed covariate and adding $batch$ and $subject$ as random effects. The R package rptR (v0.9.22)(*25*) was used to fit the model with restricted maximum likelihood and calculate bootstrapped CIs for the variation explained by donors. The number of parametric bootstraps (nboot) was set to 500.

##### *1.4 Mutation signatures*

SigProfilerSingleSample (v0.0.0.27) (https://github.com/AlexandrovLab/SigProfilerSingleSample) was used to calculate the percent of mutations in a tissue that are attributable to a given set of mutation signatures. The following parameters were used: ref="GRCh38", exome=True. COSMIC substitution mutation signatures from version 3.2 were used. To increase the power to detect mutational signatures, PZMs from all samples in a tissue were combined. To achieve accurate mutation signature decompositions, only tissues with reconstruction accuracy ≥ 95% were retained. Putative mutation signature etiologies were taken from (*26*).

#### *1.5 Algorithm for reconstructing PZM phylogenies (LachesisMap)*

##### *1.5.1 Phylogenetic reconstruction assumptions and their justifications*

Since a PZM is inherited by all descendent cells, PZMs offer a natural lineage tracing experiment. We hypothesized that the PZMs in our atlas could be resolved spatiotemporally through phylogenetic reconstruction. To ensure the highest quality of data, we only used PZMs that were found in at least two tissues in the same donor for phylogenetic reconstruction as these have the highest probability of occurring prenatally (see section Evidence suggesting multi-tissue PZMs occurred prenatally). We use the phrases **multi-tissue PZMs** and **prenatal PZMs** to describe this class of PZMs. We also confirmed that we had adequate power for detecting prenatal PZM events (see section Tree mutation detection power).

Since our dataset and study design had complex patterns of missing and uncertain data due to differential mutation detection power across the body, genome, and developmental spacetime, we deemed off-the-shelf tools for phylogenetic reconstruction to be inappropriate for this dataset (see section Challenges in PZM phylogenetic reconstruction). Therefore, we developed a novel method to reconstruct the evolutionary history of the PZMs that specifically accounted for these complexities.

Unlike other applications of phylogenetic reconstruction where the underlying topology is unknown and is thus the outcome of interest, for this application, we knew *a priori* what the topology should approximately look like from developmental biology literature. Therefore, we were interested in mapping the PZMs on this scaffold. The choice to use a mapping method rather than a building method was cemented by the fact that developmental relationships among tissues are well supported in the literature whereas using PZMs for the foundation of the tree would require using a dataset with complex patterns of uncertainty and limited size. We derived two developmental tissue trees that represent the phylogenetic relationships among GTEx tissues during human development from the literature: the **full tree**, and the simplified **germ layer tree** (see section Derivation of developmental tissue trees).

The method is based on Kimura's infinite sites model(*27*) which is commonly used in molecular and cancer evolution fields. In this parsimonious model, if a PZM is observed in more than one tissue, then it is assumed there was one mutagenic event in a common ancestor of the tissues. The common ancestor of any two or more GTEx tissues existed before the end of organogenesis, and thus represents a time point during prenatal development. The infinite sites model is a reasonable assumption since the PZM mutation rate was low (**fig. 1A**) and PZMs are expected to be under neutral selection(*28*).

We excluded multi-tissue PZMs that were found in just lung and blood as these may be blood-specific variants that were erroneously identified in the lung due to the high vascularization of the lungs. Non-lung tissues were less likely contaminated with blood since blood vessels were avoided during biospecimen collection (BBRB ID PR-0004-W1)(*29*). Therefore, we did not remove other tissue-whole blood doubletons. Cell lines were also ignored as cell line PZMs may have occurred postnatally, e.g., a multi-tissue PZM in blood and lymphoblasts may have occurred during aging.

###### 1.5.1.1 Derivation of developmental tissue trees

Since mammalian development is immensely complicated with many aspects still unknown, we used several assumptions to simplify the modelling of mutational dynamics during development. Akin to Sulston *et al*.'s seminal cell lineage map of *C. elegans*(*30*), we assumed development of tissues could be modelled as a rooted tree with directed edges (i.e., an **arborescence**). In this arborescence, each node represents a spatial-temporal region of an organism. The root node is the zygote and each directed edge represents the developmental relationship between a parent node and a child node. By definition, there is only one directed path from the root to any other node. Thus, this model assumes each primordial tissue is created from a single parental primordial tissue. While this assumption simplifies the complexities of real development, in the face of limited data and additional restrictive assumptions required for more sophisticated modeling, we deemed this simplified model to be sufficient for capturing high-level summaries of prenatal PZMs.

We defined an arborescence with the zygote as the root node, primordial tissues as the internal nodes, and the GTEx tissues as the leaves. To focus on strictly prenatal mutations, we excluded cell lines since the MRCA of a cell line and the tissue it was derived from may have occurred postnatally. Sex-specific tissues were also omitted so that data from both females and males could be easily combined together. (Of note, gonosomal PZMs (i.e., PZMs detected in both somatic and germ cells) were analyzed separately (see section Characterization of germ cell

PZMs).) A variety of literature sources were used to define the internal nodes and edges of the developmental tissue tree(*31–35*).

Parent nodes with fewer children have higher spatial-temporal resolution than nodes with more children but are more reliant on the simplifying assumption that development can be modelled as an arborescence. We strove to balance these opposing forces by defining the arborescence to have mostly bifurcating nodes. The arborescence was pruned to remove internal nodes that only had one child node since mutations cannot be mapped to such edges without additional data or additional modelling assumptions. For example, the directed path from zygote → 2-cell embryo → 4-cell embryo → … → gastrula was simplified to zygote → gastrula. The final tree (referred to as the full tree) is shown in **Fig. S5A**.

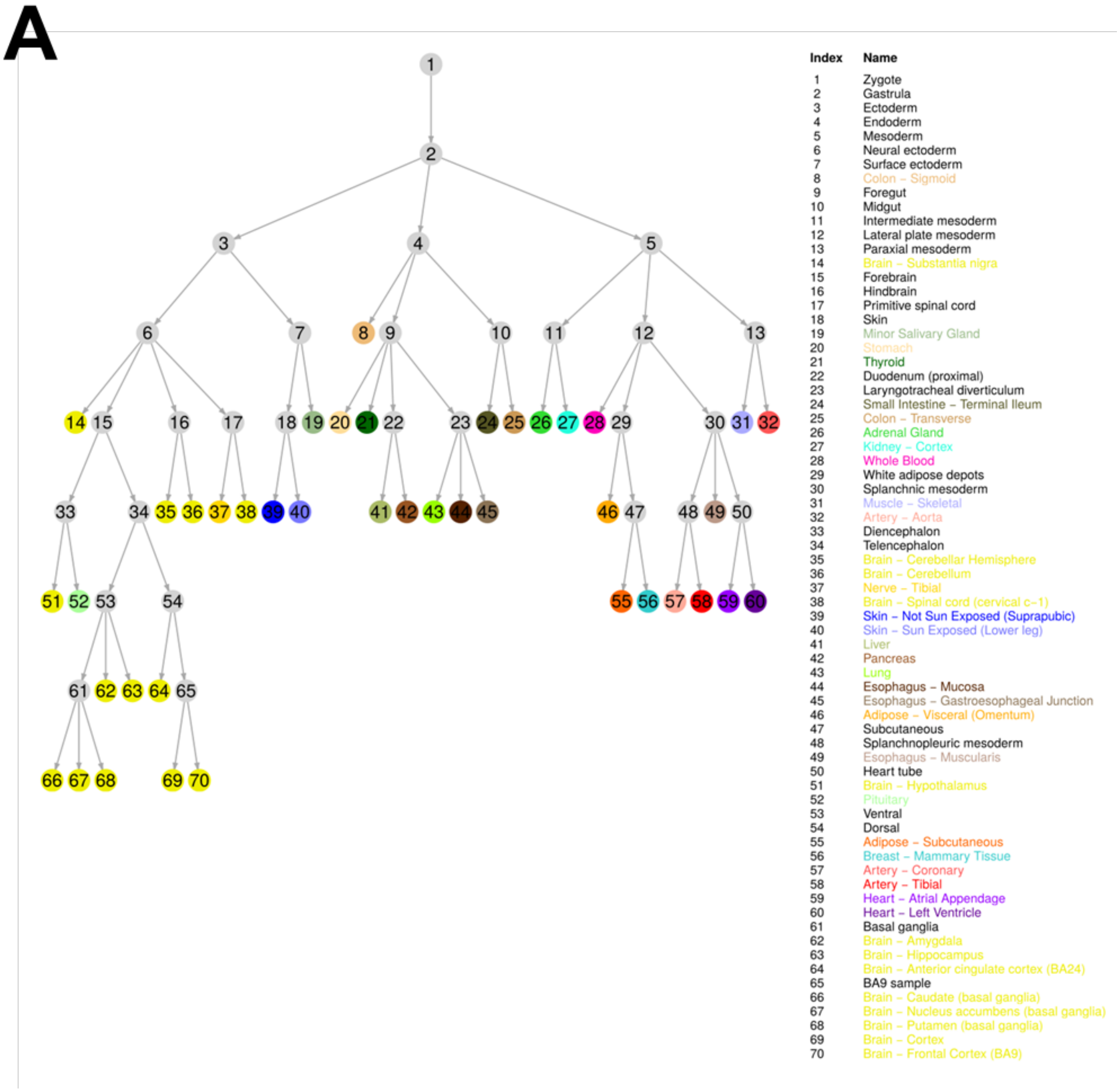
A
Index Name
1 Zygote
2 Gastrula
3 Ectoderm
4 Endoderm
5 Mesoderm
6 Neural ectoderm
7 Surface ectoderm
8 Colon – Sigmoid
9 Foregut
10 Midgut
11 Intermediate mesoderm
12 Lateral plate mesoderm
13 Paraxial mesoderm
14 Brain – Substantia nigra
15 Forebrain
16 Hindbrain
17 Primitive spinal cord
18 Skin
19 Minor Salivary Gland
20 Stomach
21 Thyroid
22 Duodenum (proximal)
23 Laryngotracheal diverticulum
24 Small Intestine – Terminal Ileum
25 Colon – Transverse
26 Adrenal Gland
27 Kidney – Cortex
28 Whole Blood
29 White adipose depots
30 Splanchnic mesoderm
31 Muscle – Skeletal
32 Artery – Aorta
33 Diencephalon
34 Telencephalon
35 Brain – Cerebellar Hemisphere
36 Brain – Cerebellum
37 Nerve – Tibial
38 Brain – Spinal cord (cervical c–1)
39 Skin – Not Sun Exposed (Suprapubic)
40 Skin – Sun Exposed (Lower leg)
41 Liver
42 Pancreas
43 Lung
44 Esophagus – Mucosa
45 Esophagus – Gastroesophageal Junction
46 Adipose – Visceral (Omentum)
47 Subcutaneous
48 Splanchnopleuric mesoderm
49 Esophagus – Muscularis
50 Heart tube
51 Brain – Hypothalamus
52 Pituitary
53 Ventral
54 Dorsal
55 Adipose – Subcutaneous
56 Breast – Mammary Tissue
57 Artery – Coronary
58 Artery – Tibial
59 Heart – Atrial Appendage
60 Heart – Left Ventricle
61 Basal ganglia
62 Brain – Amygdala
63 Brain – Hippocampus
64 Brain – Anterior cingulate cortex (BA24)
65 BA9 sample
66 Brain – Caudate (basal ganglia)
67 Brain – Nucleus accumbens (basal ganglia)
68 Brain – Putamen (basal ganglia)
69 Brain – Cortex
70 Brain – Frontal Cortex (BA9)

B

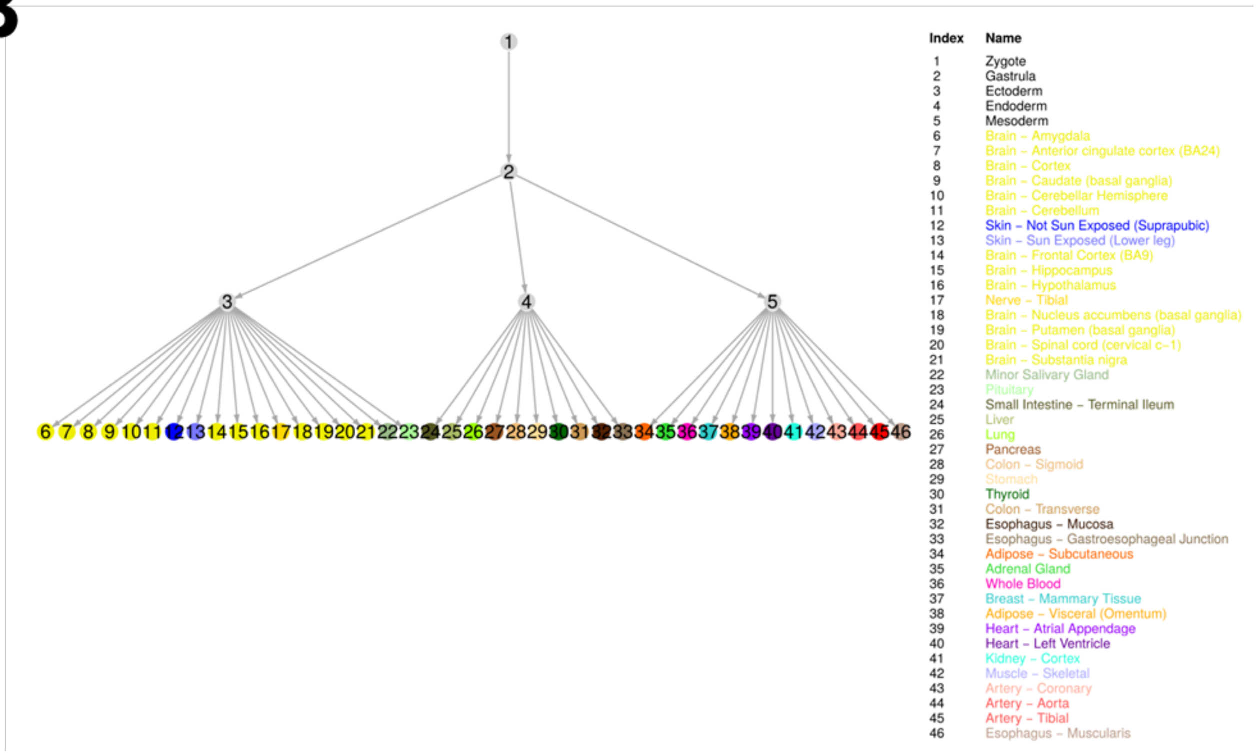

C

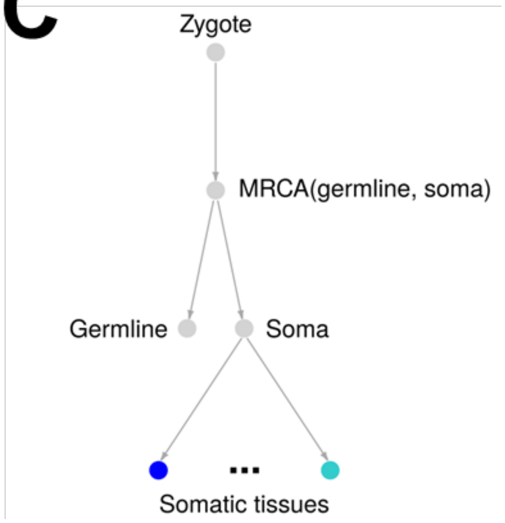

**Fig. S5. Annotated tissue trees.** (**A-C**) The root node is the zygote, primordial tissues are the internal nodes, and the GTEx tissues are the leaves. Each directed edge represents a developmental relationship such that the child node is a developmental descendant of the parent node. Each node is labeled with a number. A lookup table for converting node number to tissue name is shown. GTEx tissue nodes are colored using the GTEx coloring convention. (**A**) The full tissue tree. The full tissue tree has 70 nodes and 69 edges. (**B**) The germ layer tissue tree. The germ layer tree has 46 nodes and 45 edges. (**C**) Gonosomal tissue tree. For brevity, individual labelling of somatic tissues has been omitted.

To reduce the potential impact of incorrectly modelling development as an arborescence — but at the cost of decreasing the spatial-temporal resolution — we defined a simplified tree with only 4 internal nodes (gastrula and the three germ layers) (referred to as the germ layer tree) (**Fig. S5B**). Here, we started with the full tree and recursively merged internal nodes until only the gastrula and germ layer internal nodes were left. All downstream analyses were performed on both the full and germ layer trees.

###### *1.5.2 Challenges in PZM phylogenetic reconstruction*

There are several challenges to address when reconstructing PZM phylogenies with the GTEx study design. The challenges stem from three types of systematic differential power:

1. *Differences in detection power across the body*: since no donors had all GTEx tissues sequenced, a PZM may be undetected in a tissue simply because the tissue was not sequenced. Additionally, a PZM may be undetected in a tissue because the PZM was not present in the tissue biopsy and/or sequenced library.
2. *Differences in detection power across the genome*: since PZMs are identified from RNA-seq, a PZM may be undetected in a tissue because the gene was not expressed highly enough. Such effects are exacerbated for low VAF variants which require higher expression levels to be detected.
3. *Differences in detection power across the developmental tissue tree*: due to the incomplete and variable tissue profiling structure of the donors, specific subsets of tissues can be commonly or rarely profiled together in the cohort. Thus, different edges will have

different power to detect the mutation on the edge. Additionally, since earlier mutations are expected to 1) have higher VAF and 2) be present in more tissues than later mutations, earlier edges may have higher power than later edges in the tree. On the other hand, later mutations may be easier to detect despite being in fewer tissues because the tissues they're detected in may have more similar transcriptomes. For example, a PZM in a heart-specific gene that occurred during the formation of the heart tube may be easily detected since its expression may be high in all descendant tissues (i.e., left ventricle and atrial appendage).

We addressed these complex challenges by designing a method to control for differential power and uncertainty in the data when mapping the mutations.

##### *1.5.3 Algorithm for reconstructing PZM phylogenies*

We developed an algorithm, LachesisMap, to reconstruct the phylogenetic history of multi-tissue PZMs. The algorithm jointly analyzes all multi-tissue PZMs and accounts for missing data as well as differential PZM detection power due to differences in VAF, expression level, and tissue profiling in the dataset.

LachesisMap takes as input 1) a developmental tissue tree with edges $E = \{e_1, \ldots, e_N\}$ and 2) metadata on the $M$ multi-tissue variants ($A = \{a_1, \ldots, a_M\}$). The algorithm outputs a list of tree **edge weights** ($W = \{w_1, \ldots, w_N\}$) that represent the estimated probability of a PZM occurring in that spatiotemporal window of development.

Briefly, in step 0, the algorithm maps each multi-tissue PZM to a set of edges in the tree using parsimony and uncertainty rules. Next, in step 1, the edge weights are initialized. In step 2, the edge weights are updated by simultaneously analyzing all multi-tissue PZMs and incorporating mutation detection power. Step 2 is repeated until the edge weights converge or the maximum number of iterations is reached. The method is described in more detail below and is visualized in **Fig. S6**.

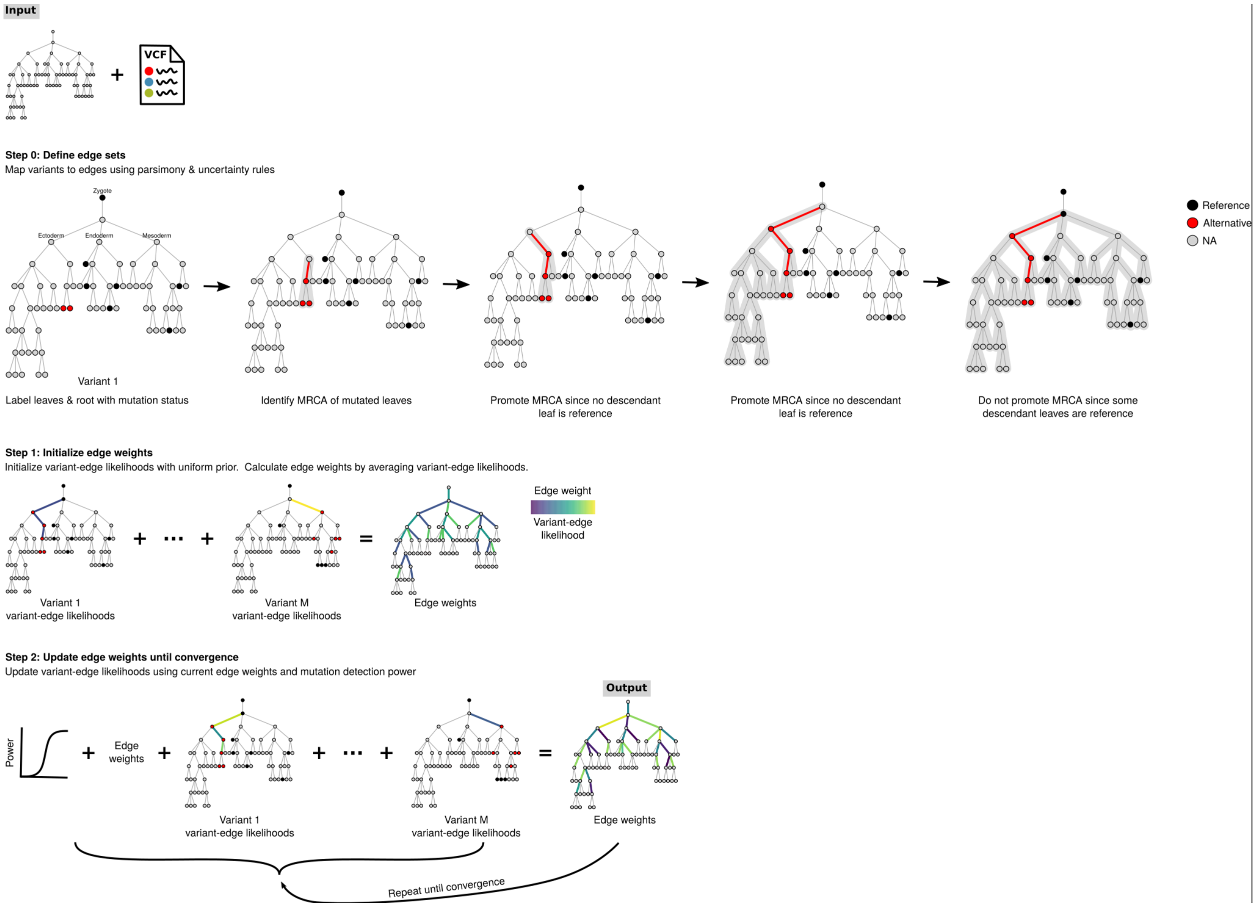

**Fig. S6. Algorithm for reconstructing PZM phylogenies.** MRCA = most recent common ancestor. NA = not applicable. VCF = variant call format.

In order to improve our estimation of edge weights, we aggregated PZMs from all donors prior to running the algorithm. This simplification assumes that mutation dynamics are similar across donors. For datasets with large mutation burdens (e.g., from WGS datasets), it may be possible to estimate donor-specific edge weights.

###### 1.5.3.1 Step 0: Define the $U_j$ edge sets

In the first step of the algorithm, each multi-tissue PZM is mapped to a set of possible edges where a mutation could have occurred using parsimony and uncertainty rules.

For each multi-tissue mutation $a_j$, label the leaves and root with their mutation status. The mutation status of a leaf is either **reference** (PZM not detected in tissue) **alternative** (PZM detected in tissue) or **NA** (tissue was not profiled or coverage was too low (< 10× coverage)). By definition, the root is labelled as reference.

Let $U_j$ be the set of edge(s) that are predicted to contain the origin of the variant $a_j$. To Initialize $U_j$, identify the most recent common ancestor (MRCA) node of all leaf nodes (i.e., GTEx tissues) containing the variant $a_j$. Add the incident edge of the MRCA node to $U_j$. Update the MRCA node to the parent of the MRCA node. Repeat the previous steps of adding edges and updating the MRCA. Terminate when the MRCA node is the root node (i.e., the zygote) or the MRCA node has one or more descendant leaves that were reference.

Step 0 highlights one of the ways LachesisMap incorporates missing data: leaf nodes with unknown mutation status have the effect of spreading their uncertainty towards the root of the tree, i.e., they have the potential to add more edges to $U_j$.

###### 1.5.3.2 Step 1: Initialize the $W$ edge weights

Define the **variant-edge likelihood**, $p_{ij}$, as the likelihood that variant $a_j$ occured on edge $e_i$. Let $n_j = |U_j|$. For each variant $a_j$, initialize the variant-edge likelihoods with a uniform distribution:

set $p_{ij} = 1/n_j$ for each edge in $U_j$. After all variants have been initialized, normalize the tree edge weights to 1: $w_i = \frac{\sum_j^M p_{ij}}{M}$.

###### 1.5.3.3 Step 2: Update the $W$ edge weights until convergence

In this step, the method analyzes all multi-tissue PZMs simultaneously to update the weights, taking into account differential PZM detection power due to differences in VAF, expression level, and tissue profiling in the dataset.

For each iteration, for each multi-tissue variant $a_j$, for each $e_i$ in $U_j$, update the variant-edge likelihoods $p_{ij}$: if $n_j = 1$, set $p_{ij} = 1$, otherwise, set $p_{ij} = w_i$. Next, adjust these variant-edge likelihoods for incomplete power. Here, we made the simplifying assumption that the observed variant-edge likelihood was proportional to the expected likelihood via the following equation: $observed\_likelihood = expected\_likelihood * mutation\_detection\_power$. Calculate the mutation detection power to detect the variant $a_j$ at edge $e_i$ given the multi-tissue variant's median VAF and median coverage using the tree power functions (see section tree mutation detection power). Update the variant-edge likelihood $p_{ij}$ to $\frac{p_{ij}}{Power(e_i, VAF_i, coverage_i)}$.

After updating all variant-edge likelihoods for all $M$ variants, normalize the weights to 1: $w_i = \frac{\sum_j^M p_{ij}}{M}$.

Repeat step 2 until $W$ converges or the maximum number of iterations is reached.

Step 2 integrates information across mutations. For example, if a particular mutation has a lot of mapping uncertainty, other mutations with more certainty will help direct which edges should have their uncertainty decreased. This feature is another way the algorithm incorporates missing data and other ascertainment biases in the data.

###### 1.5.3.4 Tree mutation detection power

The power to detect a mutation on a tree depends on the VAF, coverage, profiled tissues, and the tree topology. We estimated the power to detect a mutation on each edge of a tree as a function of these variables.

We defined the **tissue profiling vector** as the set of tissues that are profiled in a given GTEx donor and defined the **mutation vector** as the set of tissue mutation statuses. A mutation status is either reference or alternative. Of note, since we are simulating mutations, all tissues, regardless if they're profiled, will have a mutation status. We made the simplifying assumption that a mutation that occurs on a given tree edge will be present in all descendant tissues (see section Justification for not requiring monophyletic relationships during PZM mapping). Without this assumption (i.e., allowing incomplete penetrance of the mutation), it was computationally intractable to simulate all possible mutations for the large number of GTEx tissues. (To relax this assumption, one could model mutation penetrance across the nodes as a random variable, e.g., each child node has a 80% chance of inheriting the mutation.) Each edge uniquely specifies a mutation vector.

To simulate a mutation for a given edge $e_j$ and VAF, we randomly selected a tissue profiling vector from the distribution of tissue profiling vectors used in GTEx and then randomly selected GTEx samples to fit that profiling vector. Next we randomly selected a position in the allowable transcriptome. For each tissue sample, we simulated the alternative read coverage as a random binomial variable with $p$ = VAF and $N$ = the sample's coverage at that genomic position. We defined a multi-tissue mutation's coverage as the median coverage of tissues with the mutation and then discretized the coverage into one of four coverage bins.

We next ran the variant detection algorithm on the simulated multi-tissue mutations and mapped the mutation on the tree to identify the edge(s) where the mutation was predicted to have occurred (i.e., $U_j$). We conservatively defined a mutation to be predicted correctly if $U_j = e_j$. We defined the **power to detect a tree mutation** as the fraction of simulations that were predicted correctly for a given VAF, coverage bin, and mutation vector.

For each edge in a given tree and VAF, we simulated ~1.6M multi-tissue mutations (all chr22 positions in the allowable transcriptome), and calculated the tree mutation detection power for the full and germ layer trees. To examine how GTEx tissue profiling affected mutation detection, we also simulated mutations using tissue profiling vectors where all tissues were profiled.

###### 1.5.3.5 Justification for not requiring monophyletic relationships during PZM mapping

Using parsimony, our model assumes that if a mutation was detected in multiple terminal tissues, then the mutation occurred in a common ancestor edge ($U_j$). However, the model does not assume the converse, i.e., if a mutation occurred in a common ancestor edge, then the mutation was detected in all terminal tissues (i.e., a monophyletic group).

There are several biological and technical reasons why a mutation may be predicted to occur in edges $U_j$ but is unobserved in all descendent tissues. First, the internal and terminal nodes of the tree may have been seeded with cells from the parent node that did not contain the mutation. Second, the mutation may have been later eliminated in the lineage that gave rise to that tissue (e.g., from gene conversion, mitotic recombination, apoptosis, etc.(*36*)). Third, the mutation may be present in the tissue, but absent in the subset of cells that were sequenced. Lastly, the mutation may be present in the tissue and the cells that were sequenced, but still unobserved due to limited power to detect the mutation.

The lack of monophyletic relationships in PZM phylogenetic reconstruction has been previously reported. For example, Behjati *et al.* identified putative embryonic mutations predicted to occur before gastrulation but were absent from some of the profiled adult tissues(*37*). In Martincorena *et al*., skin tissue biopsies that were taken < 5 mm apart did not always contain the same PZMs(*6*).

We quantified the effect of mutation detection power to explain why a prenatal PZM may be missing from some of the descendent tissues. For this analysis, we focused on gonosomal PZMs, a subset of prenatal PZMs. (Recall that gonosomal PZMs are PZMs detected in both the germline and soma.) We used gonosomal PZMs for several reasons: since gonosomal PZMs are predicted to occur very early (within approximately the first 10 cell divisions of life (*38*)), these multi-tissue mutations are expected to be detected many tissues and have high VAFs and thus are likely to have 1) high power to detect the mutation in each tissue and 2) high power to be mapped correctly onto a tissue tree.

We first defined a **gonosomal tissue tree** which included nodes for the germline (as proxied by testis (see section Germ cell PZMs can be detected in bulk male gonads)) and the MRCA of the germline and soma (referred to as **MRCA(germline, soma)**) (**Fig. S5C**).

Similar to the previous tree mutation power simulations, we simulated mutations on the Zygote → MRCA(germline, soma) edge (i.e., gonosomal PZMs) at a variety of VAFs and coverages. Only samples from male testis donors were used. By definition, these gonosomal PZMs are present in all profiled tissues of the donor. For each simulated gonosomal PZM, we calculated the fraction of profiled tissues where the mutation was detected (referred to as the **tissue detection rate**) (**Fig. S7**). Power curves were fit to sigmoid functions using the Levenberg-Marquardt nonlinear least-squares algorithm from the minpack.lm R package (v 1.2-1) (https://CRAN.R-project.org/package=minpack.lm).

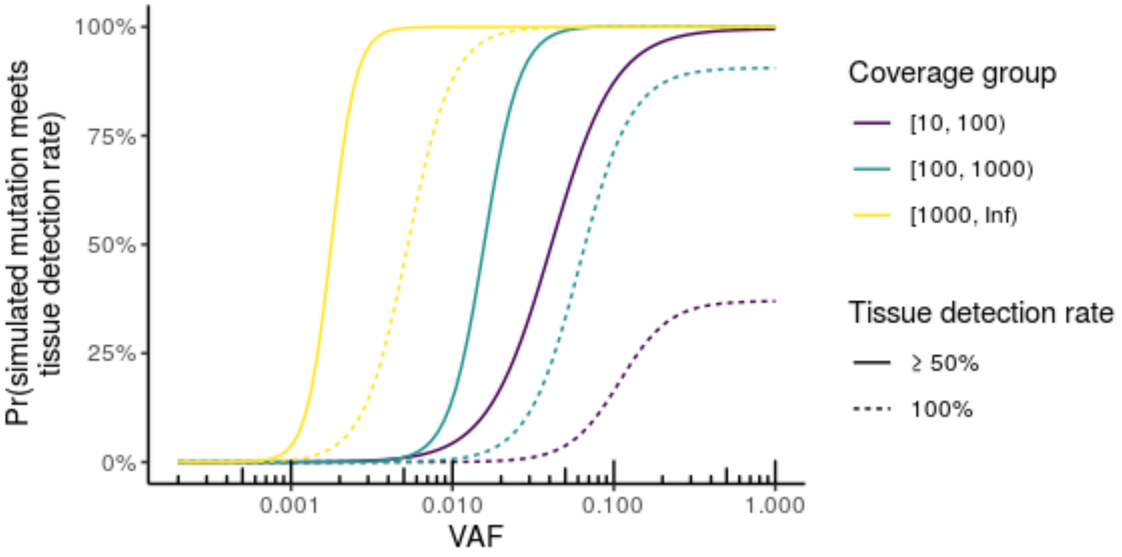

**Fig. S7. Power to detect simulated gonosomal PZMs in all tissues is variable.** Detecting a simulated gonosomal PZM in a given fraction of tissues depends on the simulated VAF and coverage (defined as the median coverage across all tissues). For a given coverage, as the simulated gonosomal PZM VAF increased, the power to detect mutations at a given minimum tissue detection rate increased. For a given simulated VAF, as the coverage increased, the power to detect mutations at a given minimum tissue detection rate also increased. Of note, power curves do not all saturate at 100%.

Using the observed gonosomal PZMs (median VAF = 0.037; mode coverage group = [100, 1000)), we predict that only 19% of gonosomal PZMs will have the mutation detected in all profiled tissues. This value jumps to 97% if the constraint is relaxed to at least 50% of profiled tissues to have the mutation detected (**Fig. S7**). Since mutation detection power is higher for gonosomal PZMs than all prenatal PZMs, these estimates likely represent an upper bound for the tissue detection rate in prenatal PZMs. Thus, failure to detect prenatal PZMs in all descendant tissues is likely very large and further justifies why we did not require PZM monophyly during PZM mapping.

##### *1.6 PZM phylogeny validation*

To test if prenatal PZMs were the result of shared developmental history rather than experimental error or multiple independent mutations, we randomly shuffled the mutation status labels of the mutation catalog 20,000 times, mapped the randomized PZMs to the tree, and calculated the distribution of expected edge weights under the null hypothesis. We used a multinomial goodness-of-fit test to test if the observed edge weight vector was likely sampled from the distribution of edge weight vectors from random mutations. Due to the large sample size, we used Monte-Carlo simulations to estimate the $P$-value using the XNomial R package (v1.0.4) (https://cran.r-project.org/package=XNomial). Since a fraction of a mutation can be mapped to an edge, both observed and expected mutation counts were rounded to the nearest integer. Edges with edge mapping power < 5% were excluded from the hypothesis tests since these edge weight estimates may be noisy.

To perform the *post-hoc* tests for testing which individual edges had significantly different burdens than the null model, we used permutation tests. For a given edge $e$ and edge weight $w$, the test statistic was defined as the absolute difference between $w$ and the median of all random $e$ edge weights. The $P$-value was defined as the percent of test statistics that were as or more extreme than the observed test statistic. $P$-values were corrected for multiple tests using the Benjamini-Hochberg procedure. q-values $\leq 0.05$ were defined as significant.

##### *1.7 Genetic variation datasets used for functional annotation*

To compare the different genetic variation datasets fairly, we strove to mimic the ascertainment biases in the PZM dataset in these other datasets. To do this, we only retained mutations that overlapped the allowable transcriptome (see Allowable transcriptome definition) unless specified otherwise.

###### *1.7.1 Inherited de novo mutations*

*De novo* mutations were pulled from denovo-db (v1.6.1), a curated database of ~628,000 *de novo* mutations from 53 studies(*39*). We used mutations from both the Simons Simplex Collection (SSC) and non-SSC subsets of denovo-db. Mutations were filtered for single-nucleotide mutations, lifted over from hg19 to hg38, and filtered for mutations overlapping the allowable transcriptome. As suggested by the authors, *de novos* from autism studies with poor validation were dropped and *de novos* from monozygotic twins and duplicate individuals were deduplicated. Mutations whose reference base in hg19 did not match the reference base in hg38 were excluded. Approximately 50,000 *de novo* mutations were retained after filtering. Mutations were partitioned into two mutually exclusive groups using the PrimaryPhenotype metadata field in denovo-db: 1) mutations observed in control subjects (defined as ***de novo* controls**) and 2) mutations observed in subjects with a disease (defined as ***de novo* cases**). Subjects with unknown disease status were excluded.

The open-access tier of simple somatic mutation datasets were used. The open-access tier excludes a small fraction of somatic mutations that overlap with the patient's germline variant. Since the fraction is small, the data censoring should only have a limited impact on the analyses. Mutations were filtered for single-nucleotide mutations that overlapped the allowable transcriptome. For computational ease but without loss of generality, cancer types with more than 180,000 mutations after filtering were randomly downsampled without replacement to 180,000 mutations. Only SKCM met this criteria. The number of cancer somatic mutations after filtering varied from ~25,000 to 180,000.

##### *1.7.2 Inherited germline variants*

Inherited germline variants were pulled from gnomAD (v3.0), a comprehensive database of germline genetic variation from ~125,000 exomes and ~15,000 genomes(*40*). Due to the large size of the dataset, we limited our analysis to variants on chr22. Variants were filtered for single-nucleotide variants that overlapped a modified allowable transcriptome. This modified allowable transcriptome did not have the GTEx germline variations removed. To reflect population allele frequencies in GTEx (where the majority of donors have self-reported European American ancestry) we used population allele frequencies from gnomAD's Non-Finnish European (NFE) population. Only variants with non-zero NFE allele frequencies were retained. Approximately 359,000 germline variants remained after filtering.

##### *1.7.3 Cancer somatic mutations*

Cancer somatic mutations in The Cancer Genome Atlas (TCGA) were pulled from ICGC release 28, a comprehensive genomic database of many cancer types(*41*). We selected six cancer types to represent the genetic diversity among cancer types. We selected cancer types whose tissue of origin was profiled in GTEx. Additionally, we selected cancers such that all three germ layers were represented by their tissue of origins and had varying degrees of somatic mutation burden. The following cancer types were selected: breast cancer (BRCA), brain glioblastoma multiforme

(GMB), liver hepatocellular carcinoma (LIHC), lung adenocarcinoma (LUAD), pancreatic cancer (PAAD), and skin cutaneous melanoma (SKCM).

##### *1.7.4 Simulated random mutations*

The point mutation rate can vary considerably across regions of mammalian genomes (reviewed in (*42*)). Therefore, we strove to model these nonuniformities in the simulated mutation datasets. Since there are many different assumptions and parameters one can use to simulate mutations, we choose two independent modelling approaches to tackle this problem. We simulated one dataset of random mutations based on human pseudogene substitutions and another dataset based on primate substitutions. The details are described below. The framework used to create the primate-based dataset was also used to train Combined Annotation-Dependent Depletion (CADD), a method later used to draw inferences about this and other datasets. Having two independent methods was one way we mitigated the circular logic of using CADD for both data simulation and evaluation.

###### 1.7.4.1 Mutations simulated from human pseudogenes

We developed a model of DNA sequence evolution based on human pseudogenes. We used a variant of the General Time-Reversible (GTR) model that specified a substitution rate parameter for each mutated nucleotide and their 5’ adjacent nucleotide context (N = (4*3)*4 = 48 rate parameters). These context-dependent substitution rate parameters were derived from >1,700 human ribosomal protein pseudogene sequences comprising >700,000 bases(*42*, *43*). Approximately 50,000 mutations were simulated in the allowable transcriptome.

###### 1.7.4.2 Mutations simulated from primate genome alignments

We next developed a model of DNA sequence evolution based on primate genome alignments. We used CADD's mutation simulator program to generate this dataset(*44*). Similar to the pseudogene-based simulated dataset, mutations were simulated from a GRT-like model that included a separate substitution rate parameter for CpGs and local (100kb) adjustment of mutation rates (N = 5*3 = 15 rate parameters for each region). Mutations were filtered for single-nucleotide mutations that overlapped the allowable transcriptome. Approximately 43,000 mutations were retained after filtering. Since the simulator generated mutations in hg19, we did not liftover the coordinates to hg38 in order to preserve the nucleotide context and local mutation rate from which the mutations were generated.

##### *1.8 PZM deleteriousness*

We used CADD (v1.6) to predict the deleteriousness of mutations(*44*). Since methods for predicting deleteriousness of non-coding mutations are less mature than protein-coding mutations, we only used protein-coding mutations for the deleteriousness analyses. Additionally, mutations on chrM and chrY were excluded as these are not annotated by CADD. If a mutation was mapped to multiple isoforms, we selected the most deleterious prediction. While this may overestimate the deleteriousness of the datasets, since this approach was used for all datasets, the *relative difference* in deleteriousness between datasets should remain unbiased. To avoid potential issues with model misspecification, we annotated datasets using CADD models that were trained on the same reference genome as the original dataset. The simulated random dataset based on primate alignments and the cancer datasets were annotated with CADD model GRCh37-v1.6; the remaining were annotated with CADD model GRCh38-v1.6.

We used two approaches to compare mutation deleteriousness between datasets. In the first approach, we plotted the complementary cumulative density function (CCDF) of PHRED-scaled CADD scores for each dataset, i.e., $Pr(X \geq x)$. Since we were most interested in differences in the proportion of deleterious mutations which have large scores, CCDFs were an appropriate method to use because they highlight differences in the right tails of distributions. The greater the area under the CCDF, the higher the proportion of deleterious mutations in the dataset. While the CCDF method is simple to interpret, it does not provide a way to find statistically significant differences in deleteriousness nor does it control for technical covariates. Therefore, we developed a second method to account for these limitations.

For the second approach, we compared the odds of detecting a deleterious mutation in one dataset compared to another dataset while controlling for technical covariates using logistic regression. A mutation was defined as **deleterious** if the PHRED-scaled CADD score $\geq 20$. The CADD authors recommend cutoffs between 10-20 for defining deleterious mutations. Results were generally similar for more stringent cutoffs. The data was fit using the model $is\_deleterious \sim dataset + mutation\_detection\_power$ and logistic regression. The odds of detecting a deleterious mutation in one dataset versus another was derived from the fitted model coefficients.

$mutation\_detection\_power$ was included since PZM mutation deleteriousness may be coupled to VAF and gene expression level. Since we had already filtered all datasets to the same allowable transcriptome, which genes the mutations were found in was likely not a technical confounder so gene was not added to the model. The relative position of the mutation along the coding sequence was also examined as a potential confounder. Within a dataset, for the CADD score cutoff of interest (≥ 20), scores were generally similar in the first half of the gene body compared to the last half. Therefore, the srelative position along the coding sequence was not added to the model.

For PZM datasets, $mutation\_detection\_power$ was derived from power simulations (see Step 1 PZM detection power). For the remaining datasets, since the variant VAFs were generally orders of magnitude larger than PZMs VAFs (e.g., germline variants have VAFs ~50% or ~100%), we made the simplifying assumption that $mutation\_detection\_power$ was 100% for all variants.

Based on the CCDFs (**Fig. S18C**), the difference in deleteriousness across datasets was strongest for higher CADD scores. Therefore, a cutoff of 25 was used for this analysis instead of the default CADD score cutoff of 20.

###### *1.8.1 PZM deleteriousness validation*

The same mutation deleteriousness analysis pipeline was used for estimating PZM deleteriousness in the independent PZM datasets except for the following minor change. When calculating odds ratios, we dropped $mutation\_detection\_power$ from the model. This is because mutation detection power was unknown for the validation PZM datasets and could not be easily estimated.

For GTEx-based datasets, only PZMs from GTEx pass QC tissues were used. For datasets with multiple tissues, putative prenatal PZMs were defined as PZMs found in two or more tissues from the same donor. Putative postnatal PZMs were defined as PZMs found in a single tissue in a donor. For TwinsUK, lymphoblast cell lines (LCLs) and blood were treated as a single tissue, e.g., if a PZM was detected in just LCL and blood, the PZM was predicted to be postnatal. (Cell line data in the Yizhak *et al.* and García-Nieto *et al.* datasets was already excluded by the original authors.)

The following sections describe how the PZM validation datasets were processed.

###### 1.8.1.1 TwinsUK dataset

###### 1.8.1.1.1 Samples

We used the subset of TwinsUK with RNA-seq profiling data. This data was generated from the Biomarkers of Ageing using whole Transcriptome Sequencing (EuroBATS) project. Library preparation, sequencing and quality control are described in detail in (*45*). Briefly, RNA-seq samples were prepared using a polyA, unstranded RNA-seq protocol (Illumina TruSeq) and sequenced on Illumina HiSeq instruments (HiSeq 2000) with 49bp paired-end reads to a median coverage of ~36M total reads.

We converted BAMs to FASTQ format using Picard Tool's SamToFastq (v2.9.4) (https://broadinstitute.github.io/picard/). Fastqs were then processed through the same alignment and postprocessing steps as GTEx samples. PZM variant calling of TwinsUK samples was performed in a similar manner as GTEx except that RIN was not used to remove low quality samples as this information was not readily available. After our quality control, there were 2,382 samples from 811 donors from 4 tissue and cell types (unexposed skin (derived from a punch biopsy near the umbilicus), subcutaneous adipose (derived from the same punch biopsy), peripheral blood, and LCLs (derived from Epstein-Barr virus (EBV) transformation of B-lymphocytes in peripheral blood).

###### 1.8.1.1.2 Germline variant calling

Low read depth whole-genome sequencing was performed for a subset of TwinsUK donors as part of UK10K(*46*). Library preparation, sequencing, quality control, alignment, and germline variant calling are described in detail in (*46*). Briefly, Illumina paired-end DNA libraries were generated from peripheral blood mononuclear cells and were sequenced on Illumina HiSeqs with 100bp paired-end reads to an average coverage of 7X. Reads were aligned to GRCh37 using BWA (v0.5.9-r16). SNVs and indels were called using samtools/bcftools (v0.1.18-r579).

An additional subset of TwinsUK donors were re-sequenced to higher depth (> 30× coverage) (*47*). Sample preparation, sequencing and analysis is discussed in detail in (*48*). Briefly, DNA was extracted from serum and sequenced on Illumina HiSeq X instruments using 2x150 bp paired-end reads. Reads were mapped to GRCh38 using Isaac Genome Alignment Software and variants were called using ISIS Analysis Software (v.2.5.26.13)(*48*).

In total, germline variant calls were available for 405 donors. Using the definition that inherited germline variants are the same in monozygotic twin pairs, germline variant calls were ostensibly available for 532 donors.

###### 1.8.1.1.3 PZM variant calling

We ran LachesisDetect on the TwinsUK samples to detect PZMs. An additional strand bias filter was implemented that removed mutations whose alternative allele-containing alignments were biased to a genome strand (Fisher's exact test). Strand bias *P*-values were corrected using the Benjamini-Hochberg procedure. Mutations with strand bias q-values ≤ 0.05 were retained. This additional filter removed ~1.3% of the mutations. Approximately 1,700 PZMs were detected.

###### 1.8.1.2 Yizhak et al. dataset

PZMs were taken from Table S3 of (*49*). PZMs were detected in GTEx v7 data. RNA-MuTect was used to call PZMs from bulk RNA-seq data. Using the columns of Table S3, VAF was defined as Alternate. Count / (Alternate. Count + Reference. Count).

###### *1.8.1.3* García-Nieto *et al.* dataset

PZMs were taken from Table S3 of (*50*). PZMs were detected in GTEx v7 data. A custom method was used to call PZMs from bulk RNA-seq data. Using the columns of Table S3, VAF was defined as alt_count/coverage.

###### *1.8.1.3.1* Brazhnik *et al.* dataset

VCFs of the PZMs were obtained from the authors of (*51*) (personal communication). PZMs were detected in DNA from single-cell human liver hepatocytes, liver stem cells, and organoids derived from them. Putative PZMs were called using VarScan2, MutTect2, and HapolotypeCaller. A PZM was considered detected if it was detected by three variant callers.

##### *1.9 Selection*

We used dN/dS to estimate selection pressure for each mutation dataset. dN/dS measures the relative balance between adaptive and functionally constrained changes in protein-coding genes. dN/dS is frequently used to estimate selection in species evolution(*52*, *53*) and is becoming increasingly more common in cancer biology(*28*) and mosaicism fields(*6*, *54*).

dN/dS was calculated using the R package dndscv (v0.0.1.0)(*28*) for missense and nonsense mutations. The dN/dS of splicing mutations was not evaluated because exon junctions were excluded during variant calling due to the possibility of higher false discovery rates. dndscv provides three substitution models: 1) the simplest model with N = 2 rate parameters (transitions and transversions); 2) an intermediate model with N = 12 rate parameters (one for each combination of the six base substitutions on either the transcribed or non-transcribed strand); and 3) a complex model with N = 192 rate parameters (one for each combination of the six base substitutions, transcription strand, and the sixteen possible bases immediately 5' and 3' to the mutated base). We experimented with all three models. Based on the models' Akaike information criterion, potential for overfitting on the dataset sizes, and the importance of capturing context-dependent heterogeneity in mutation rates across the transcribed exome, we selected the intermediate 12-rate model (sm="12r_3w") for PZM datasets and the complex 192-rate model (sm="192r_3w") for the non-PZM datasets. dndscv was run without reference covariates (cv=NULL) since this parameter did not impact on the global dN/dS estimates. Datasets were analyzed using the reference assembly originally used to call mutations (refdb="hg19" for the simulated random dataset based on primate alignments and the cancer datasets and rebdb="RefCDS_human_GRCh38.p12.rda" for all other datasets). RefCDS_human_GRCh38.p12.rda was downloaded from https://github.com/im3sanger/dndscv_data/blob/master/data/RefCDS_human_GRCh38.p12.rda on 8/18/2020.

For improved accuracy, when donor-level information was available, we removed contiguous substitutions, e.g., dinucleotide substitutions, within a donor prior to calculating dN/dS. This is because complex substitutions can be better modeled with alternative models than those used for single-base substitutions(*28*). This filter removed 0-2% of mutations from each dataset.

##### *1.9.1 Definition of cancer genes*

Table S3 Martincorena *et al.*(*28*) was used to define the set of cancer genes. These 369 high-confidence cancer genes were compiled using COSMIC classic genes and genes significantly mutated in pan-cancer studies. The gene list was filtered to the allowable transcriptome. This filter removed < 1% of the genes. The set of non-cancer genes was defined as the set difference of all genes in the allowable transcriptome minus the cancer genes in the allowable transcriptome.

Due to the large size of the germline variant dataset, only variants on chr22 were analyzed. The cancer and non-cancer gene sets were also filtered to chr22 for the germline variant dataset.

#### ***1.10 Germ cell PZM analyses***

##### *1.10.1 PZM variant filtering*

In a prerequisite analysis, we observed that donors who were not genotyped had slightly more gonosomal mutations than genotyped donors after accounting for other technical and biological covariates. Additionally, testis PZMs from non-genotyped donors had slightly larger VAFs than genotyped donors. These results suggest that mutations from non-genotyped testis donors may have a higher contamination of inherited germline variants. To avoid systematic bias among the donors and to minimize contamination from false positive inherited germline variants, we excluded non-genotyped testis donors from germ cell PZM analyses.

*1.10.2 Sperm sequencing experiments*

**1.10.2.1 Study design**

An overview of the sperm PZM study design is shown in **Fig. S8**. Ejaculated sperm and venous blood were collected from a European American. Sperm samples had normal sperm density, sperm motility and morphology.

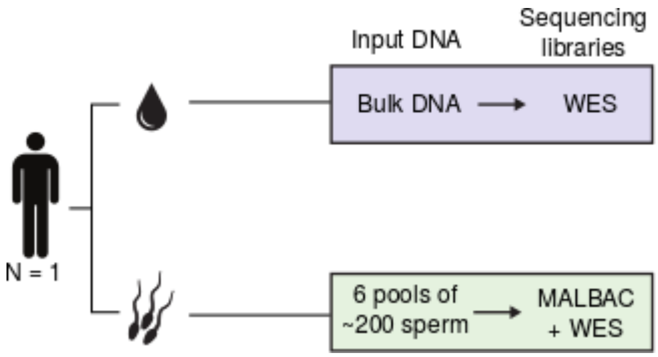

**Fig. S8. Sperm PZM study design.** Genomic DNA was extracted from blood of a male donor and subjected to standard exome sequencing. Genomic DNA was extracted from six pools of sperm from the same donor with approximately 200 sperm in each pool. These six DNA samples were amplified using MALBAC followed by standard exome sequencing library preparation and sequencing. WES = whole exome sequencing. MALBAC = Multiple annealing and looping–based amplification cycles.

**1.10.2.2 Sperm staining and FACS**

Fresh ejaculates were diluted 1:1 with sperm medium (EmbryMax HTF, Millipore) and incubated for 30 minutes at 36°C. After this incubation period, 1 ml of the diluted sperm sample was stained using the LIVE/DEAD Sperm Viability Kit (Invitrogen). 5 µl of a 1:50 dilution in sperm medium of SYBR-14 stock solution was added to the diluted sperm sample and the mixture was incubated for 10 minutes at 36°C. Next, 5 µl of the propidium iodide (PI) stock solution was added to mixture and incubated for another 10 minutes at 36°C.

Sperm samples were then selectively sorted via fluorescence-activated cell sorting (FACS) into 96 well plates (~200 sperm cells per well) and 5 ml Falcon tubes based on their staining, i.e., positive for SYBR-14 and PI (i.e., dead cells) or positive for SYBR-14 (i.e., live cells). The live/dead ratio of the sperm cells was recorded at the start of the FACS session as well as at the end of the session.

###### 1.10.2.3 DNA extraction from sorted sperm cells

DNA was extracted from three pools of live sperm and three pools of dead sperm for a total of six pools. While live/dead status was used during sample preparation, due to the limited number of PZMs detected in each group and the large overlap of PZMs detected in live and dead sperm groups, the groups were combined during analysis.

DNA was extracted using a lysis protocol as initially described in (*55*). Briefly, a stock solution of a lysis buffer was added to the PBS containing the sorted sperm cells resulting in mixture that had a final concentration of 1 M Tris·Cl, pH 8.0, 3M NaCl, 0.5 M EDTA and 20% SDS. To this mixture, DTT and proteinase K were added to a final concentration of 40 mM DTT and 10 mg/ml proteinase K and incubated overnight at 50°C. The following day, 2 volumes of ice-cold EtOH were added to the sample and a precipitation was performed for 2 hours at -20°C. Next, the DNA was pelleted by centrifugation for 20 minutes at 15,500xg at room temperature. The supernatant was removed and the DNA washed with 75% EtOH. The DNA was pelleted by centrifugation for 10 minutes at 15,500xg at room temperature. Finally, the supernatant was removed and the pellet was air-dried and dissolved in 50 µl dH20.

###### 1.10.2.4 Whole genome amplification and exome sequencing library preparation

We used MALBAC amplification(*56*) to prepare up to 1.5 µg of DNA from each pool of sperm using a kit according to the manufacturer's protocol (Yikon Genomics). Exome library preparation was performed according to the manufacturer's protocol (Illumina Nextera Rapid Capture Exome kit, Illumina) using 50 ng of pre-amplified MALBAC reactions or DNA extracted from blood. Size distribution of the enriched libraries was assessed by analyzing 1 µl of each library on an Agilent 2100 Bioanalyzer. The resulting six MALBAC libraries and standard blood library were sequenced by Illumina sequencing with 2x101 bp paired-end reads (sperm: 14M - 24M mapped reads; blood: 87M mapped reads).

###### 1.10.2.5 Alignment and mutation calling

The quality of the raw sequencing data was assessed with FastQC (http://www.bioinformatics.babraham.ac.uk/projects/fastqc/). The first 20 bp of the reads were trimmed using TrimGalore (v0.4.5) (https://www.bioinformatics.babraham.ac.uk/projects/trim_galore/). Resulting reads were processed according to the GATK best practices and were mapped to the hg19 reference genome using BWA-MEM (v0.7.13)(*23*) and Picard Tools (v2.8.3) (http://broadinstitute.github.io/picard/). To achieve a high quality mutation call set, only properly mapped reads (flag 0x2) and alignments with $NM \leq 3$ were retained. Single-nucleotide PZMs were called with MuTect (v1.1.4)(*57*). Each sperm sample was run against the blood sample as a reference to identify sperm-specific mutations.

###### 1.10.2.6 Validation of sperm PZMs

To validate sperm PZMs, amplicon sequencing of 144 regions was performed using Custom TargetGxOne amplicon primers and library preparation by GENEWIZ (www.genewiz.com). Libraries were sequenced on Illumina MiSeq with 2x250 bp paired-end reads. Reads were aligned using BWA and Picard Tools. Data at mutated loci was collected from the resulting BAM files with SAMtools mpileup(*1*). Variant calling was performed using VarScan (v2.4.2)(*58*). A PZM was defined as validated if the identical nucleotide change was detected at the same position as predicted by MuTect and the nucleotide change was not observed in blood. Only validated PZMs were used for downstream analyses (N = 93 PZMs). Validated PZMs were lifted over to hg38 and filtered to the allowable transcriptome (N = 84 PZMs).

##### *1.11 Calculation of mutation rates during gametogenesis*

We used a similar method as Rahbari *et al.*(*38*) to calculate mutation rates during gametogenesis. Gametogenesis in males can be partitioned into three stages: prior to primordial germ cell (PGC) specification, after PGC specification, and after puberty.

To calculate the mutation rate after puberty, we used the following linear model to fit the number of germ cell-specific PZMs in a donor using Poisson regression:

$$\#\ germ\ cell\ specific\ mutations \sim AGE + sample\_mutation\_detection\_power \qquad \text{1-8}$$

where $AGE$ was the donor's age and $sample\_mutation\_detection\_power$ was the average mutation detection power of simulated mutations in the donor's testis sample. The $AGE$ model coefficient (in units of # of germ cell-specific mutations / year) was divided by 23 (the approximate number of cell divisions per year)(*38*) to get the # of germ cell-specific mutations / division.

We next calculated the post-PGC mutation rate. There are approximately 24 cell divisions between post PGC specification and puberty(*38*). Thus the number of germ cell-specific mutations that are observed at puberty represent the mutations that occurred during this stage of gametogenesis. Using the previous model, we estimated the number of germ cell-specific mutations at puberty using $AGE = 14$ (an approximate age for male puberty) and $sample\_mutation\_detection\_power = Median(sample\_mutation\_detection\_power\ testis\ samples\ )$. The mutation burden was divided by 24 cell divisions to calculate the mutation rate.

To calculate the pre-PGC mutation rate, we fit the number of gonosomal PZMs in a donor to a Poisson distribution. The expected number of gonosomal PZMs in a donor was divided by 10 cell divisions (the approximate number of cell divisions before PGC specification)(*38*) to calculate the mutation rate.

##### *1.11.1 Germline surrogate tissue analysis*

We fit the following general linear mixed effects logistic regression model to predict whether a gonosomal PZM was detected in a somatic tissue given information about the gonosomal PZM in the germ cell and the somatic tissue.

$$\begin{aligned} is\_detected.non\_testis\_tissue \sim \text{tissue} + \log_{10} VAF.testis\_tissue \\ + \log_{10}(coverage.non\_testis\_tissue\ + 1)\ +\ power\ + AGE \\ + (1|donor\_mutation) \end{aligned} \tag{1-9}$$

where $donor\_mutation$ was modeled as a random effect and $power$ was the power to detect a mutation of $VAF = VAF.testis\_tissue$ and $coverage = coverage.non\_testis\_tissue$ in the somatic tissue.

The model was fit using the glmer function from the R package lme4 (v1.1-26)(*59*) and used option nAGQ=9 to improve the accuracy of the numerical integration.

Importantly, the model controlled for technical covariates such as expression level in the somatic tissue of interest. The predictions of the model, i.e., the probability of detecting a gonosomal PZM in a somatic tissue, are the cumulative effect of both the interesting, biological covariates and the nuisance, technical covariates of the study design. In order to focus on just the biological features, we analyzed the model coefficients, i.e., the odds of detecting a gonosomal PZM in one tissue versus another. Since blood is arguably the most common biospecimen collected for genetic studies and genetic counseling, we compared all tissues to blood.

#### 2 Supplementary results

The cross-sample calling in step 3 allowed us to identify variants that had lower total coverage and lower alternative allele coverage than those identified in step 1 (both *P*-values < 2.2E-308, Mann-Whitney *U* test) (**Fig. S9**). These newly detected PZMs also tended to have higher VAFs (*P*-value = 6.4E-268, Mann-Whitney *U* test) (**Fig. S1D**). As expected, step 3 increased the prevalence of multi-tissue PZMs (*P*-value = 3.2E-314, McNemar's Chi-square test with continuity correction) and increased the number of samples containing the multi-tissue PZMs by 56% on average. By including a random set of matched GTEx samples in step 2, we estimated the FDR to be 9.3%.

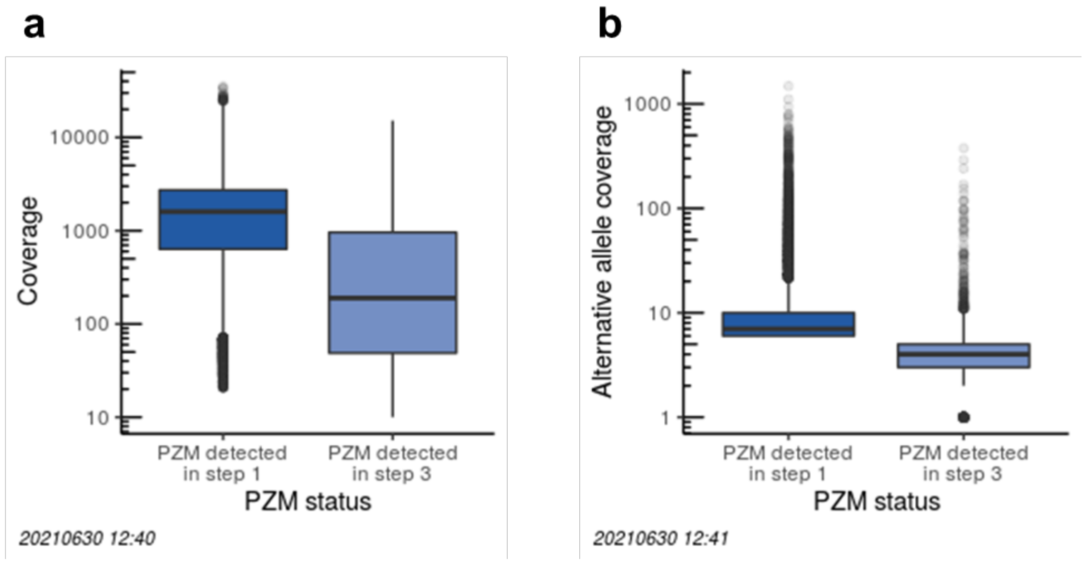

**Fig. S9. Distribution of step 1 and step 3 PZM statistics.** Distribution of total coverage (**A**) and alternative coverage (**B**) for PZMs detected in step 1 and step 3. Top and bottom of the boxes denote first and third quartiles, respectively; horizontal black lines denote medians; whiskers denote 1.5× the interquartile range; and outliers are plotted with transparency to disambiguate overlapping values.

##### *2.1 PZMs in hypermutated samples are likely false positives*

A small subset of normal GTEx samples (~5%) had an extraordinarily high PZM burden. Hypermutation is observed in cancer at relatively low rates (~17% of adult cancers) and can result from both extrinsic and intrinsic mutagenic exposures, e.g., UV light(*60*). We hypothesized that the GTEx hypermutated samples may be a novel form of hypermutation in normal tissues. We used experimental validation data to determine if these PZMs represented a technical artifact or a biological phenomenon.

Across the four experimental validation datasets, we attempted to validate 1,737 putative PZMs across 17 hypermutated samples. Validation data was informative for 1,509 PZMs. The average FDR was 98%. Thus, the overwhelming majority of PZMs in hypermutated samples are likely false positives. As a result, the observed hypermutated phenotype is likely a technical artifact. Given that the validation data was generated from tissue biopsies different from the ones used for the original mutation calling, it is possible we may have been underpowered to detect PZMs in the validation sample (e.g., due to tissue mosaicism). However, since non-hypermutated samples had a much lower FDR, this possibility is not very likely.

#### *2.2 PZM features recapitulate known biology*

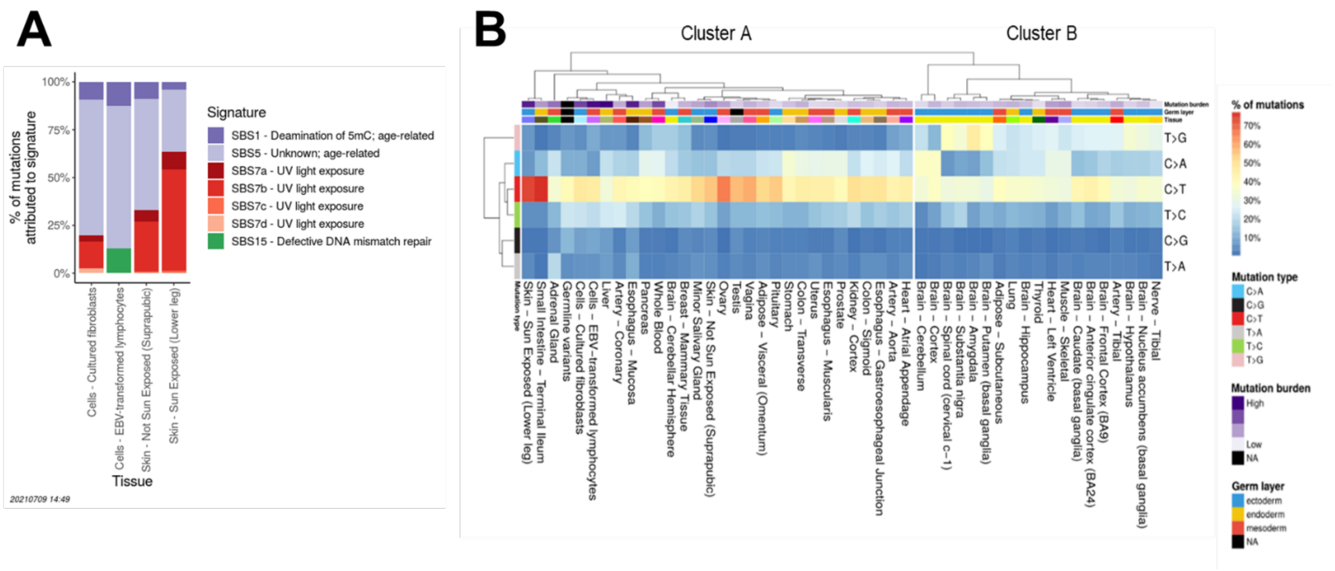

**Fig. S10. PZM mutation spectra.** (**A**) Percent of mutations attributed to mutation signatures across tissues. For all tissues except sun-exposed skin, most mutations are predicted to be from age-related mutagenic processes (purple shades). All skin-related tissue/cell types have evidence of mutations from UV light exposure (red shades), with sun-exposed skin having the greatest percentage. Only tissues with reconstruction accuracy ≥ 95% are shown. SBS = substitution mutation signature. (**B**) PZM mutation spectra clustered by tissue and mutation type.

We compared the mutation burden in sun-exposed skin versus unexposed skin as a function of self-reported ancestry. As expected, European Americans had a higher mutation burden in sun-exposed skin compared to unexposed skin (*P*-value < 2E-16, Mann Whitney *U* test) whereas there was no detectable difference in African Americans. Additionally, European Americans had a higher mutation burden in sun-exposed skin than African Americans, but no difference in burden in unexposed skin (*P*-values = 5E-12, not significant (NS), respectively, Mann Whitney *U* test) (**Fig. S11A**). Sun-exposed skin had a greater fraction of both C>T and CC>TT PZMs than non-skin tissues (**Fig. S11B** and **C**). As darker skin contains more photoprotective melanin(*61*) and UV mutagenesis results in C>T and CC>TT mutations(*62*), these results support the hypothesis that the PZMs were caused by UV damage. Together, they add further evidence that LachesisDetect produces high-quality mutation predictions.

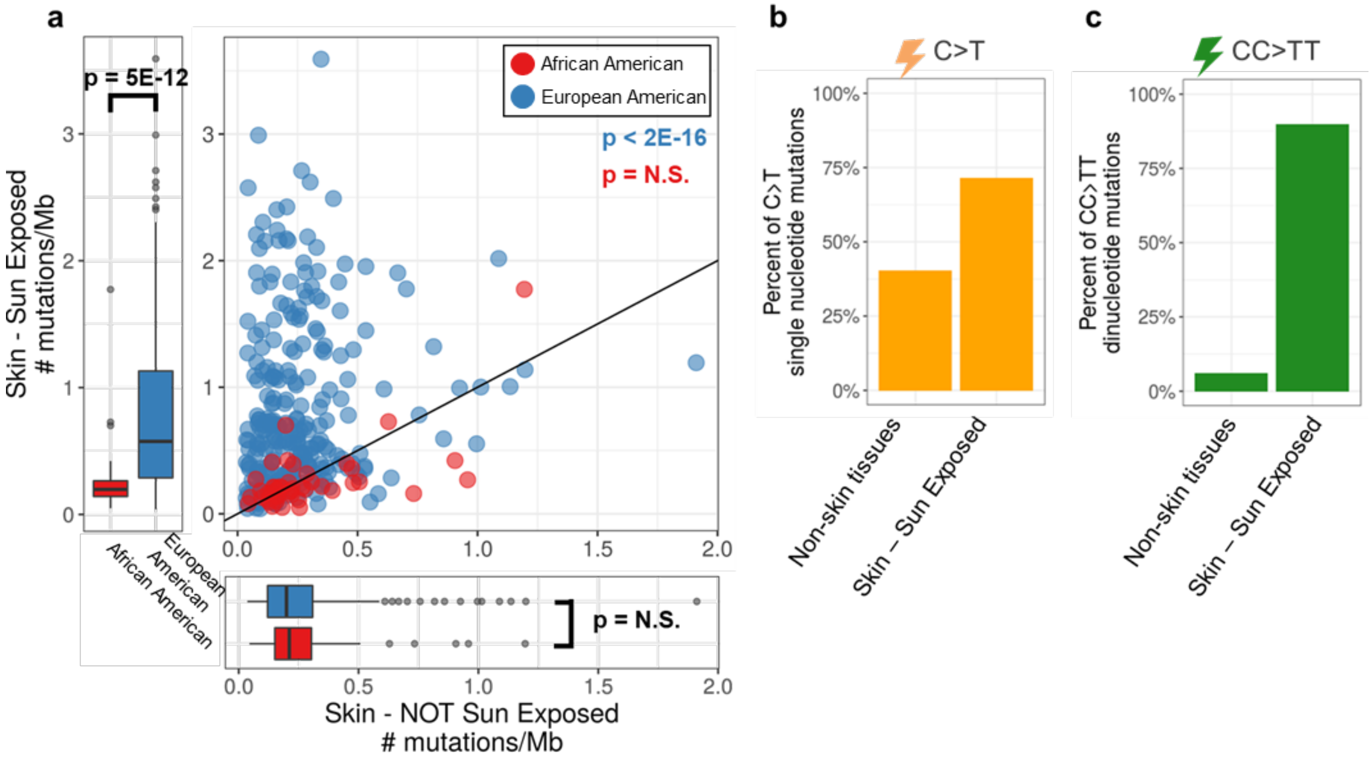

**Fig. S11. Skin PZMs recapitulate known biology.** (**A**) Joint distribution of PZM mutation burden in sun-exposed skin versus unexposed skin as a function of self-reported ancestry. Y = X line denotes equal burden across the two skin regions. Paired data used for the joint distribution (scatterplot) whereas all data (paired and unpaired) used for the univariate distributions (boxplots). Top and bottom of the boxes denote first and third quartiles, respectively; horizontal black lines denote median; whiskers denote 1.5× the interquartile range; and outliers are plotted with transparency to disambiguate overlapping values. (**B**) Fraction of single-nucleotide mutations that are C>T in sun-exposed and non-skin tissues. (**C**) Fraction of dinucleotide mutations that are CC>TT in sun-exposed and non-skin tissues. N.S. = not significant.

Another well supported mosaicism phenomenon is CHIP(*63*). CHIP PZMs were detected in blood and EBV-transformed lymphoblasts at expected rates (see Prevalence of expressed CHIP mutations)(*64*). This result provides additional evidence that the PZM call set is of high quality and has moderate sensitivity. To our knowledge, this is the first time CHIP mutations have been detected at the RNA level and thus suggests these mutations may have a direct functional role in clonal growth.

##### *2.3 Preservation method does not likely affect mutation burden and spectra*

During biospecimen collection, GTEx used different tissue preservation methods. Eleven of the thirteen brain regions were processed from fresh frozen tissue whereas the remaining two brain regions and all non-brain tissues (N = 35) were processed from PAXgene preserved tissue (**Fig. S12A**). This less than ideal choice of systematic differences in sample collection was likely driven by necessity as frozen brain samples were processed by a designated brain bank and the remaining samples were processed by GTEx.

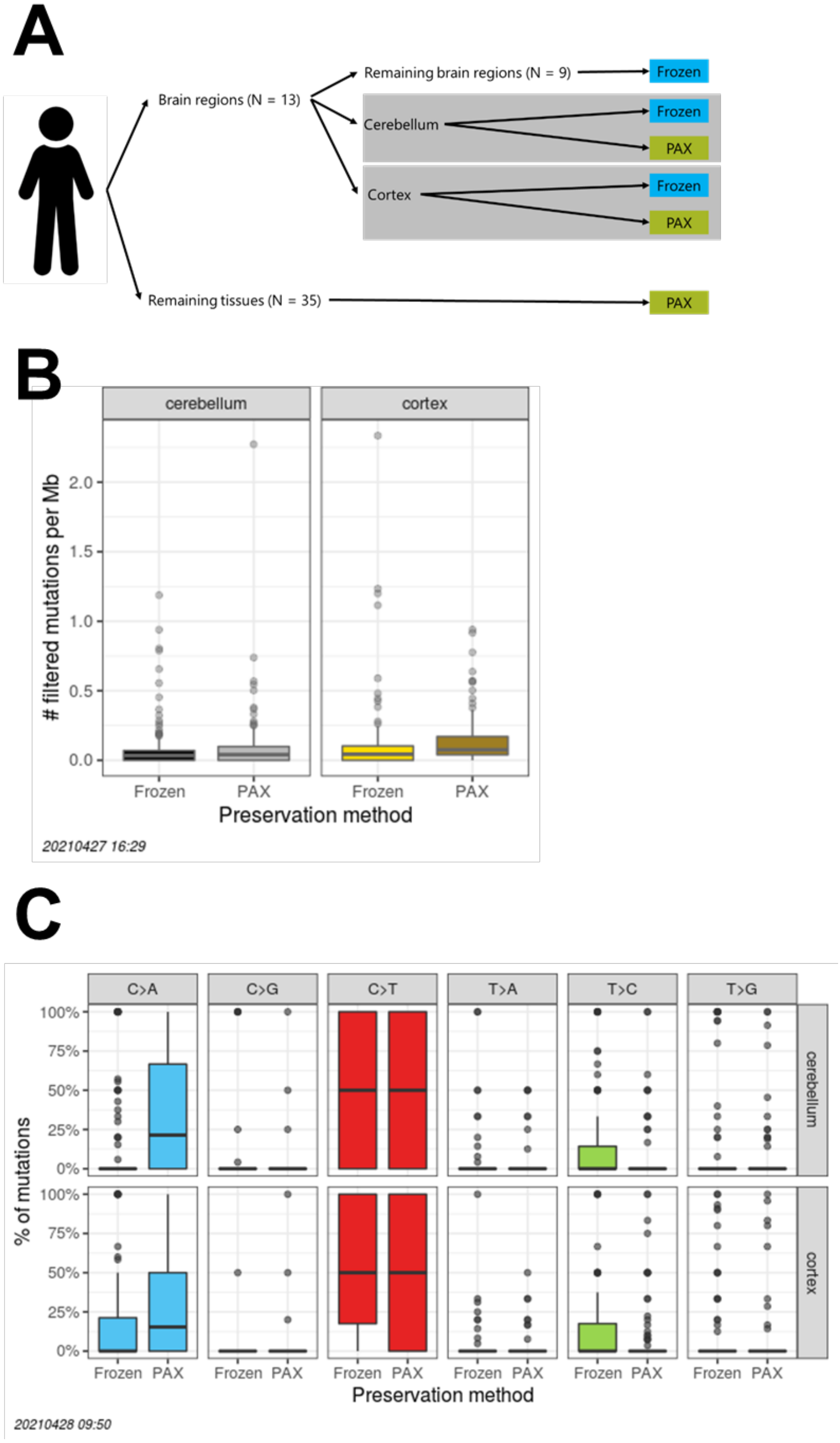

**Fig. S12. Preservation method does not likely affect mutation burden and spectra.** (**A**) Overview of the GTEx preservation schema. (**B**) Normalized mutation burden in paired frozen and PAXgene preserved cerebellum (**left**) and cortex (**right**) brain regions. Tissues are colored in their canonical GTEx tissue colors. (**C**) Mutation spectra in paired frozen and PAXgene preserved cerebellum (**top**) cortex (**bottom**) brain regions. Top and bottom of the boxes denote first and third quartiles, respectively; horizontal black lines denote median; whiskers denote 1.5× the interquartile range; and outliers are plotted with transparency to disambiguate overlapping values.

While fresh frozen is the gold standard for tissue preservation, alternatives like PAXgene are commonly used. PAXgene is a fixation method used to simultaneously preserve morphology and biomolecules. Importantly, PAXgene does not involve crosslinking or other chemical alterations to biomolecules(*65*). Several papers have found that PAXgene preserves the integrity of nucleic acids and produces results similar to fresh frozen samples (e.g., high correlation in expression(*66*) and high correlation in methylation (*67*)). To our knowledge, there have been no studies that have compared variant calls in fresh frozen and PAXgene RNA-seq data. The closest study was Högnäs *et al.* where they compared experimental error rates in targeted DNA sequencing data(*68*). No differences in mismatch rate or the distribution of incorrect basecalls types were detected between samples that were fresh frozen and samples that were preserved with PAXgene. This suggests the fidelity of DNA mutation calling from PAXgene preserved RNA may be high.

Given the lack of precedent of calling PZMs from PAXgene preserved tissue, we hypothesized that the differences in preservation method among tissue types might drive differences in mutation burden and spectra. We focused our analyses on the cerebellum and cortex samples because these brain metaregions each contained a pair of fresh frozen and PAXgene samples (N = 152 paired cerebellum samples; N = 115 paired cortex samples). These paired samples were taken from the same brain metaregion but unfortunately were most likely not from the same biopsy/core.

For the cerebellum metaregion, no difference in normalized mutation burden was detected between PAXgene and fresh frozen tissues (*P*-value = 0.44, Wilcoxon signed rank test). For the cortex metaregion, PAXgene samples had slightly larger mutation burdens (0.08 PZMs/Mb vs. 0.03 PZMs/Mb, *P*-value = 0.03, Wilcoxon signed rank test) (**Fig. S12B**).

We next compared the mutation spectra between paired fresh frozen and PAXgene preserved tissues. Mutation spectra was associated with preservation method for both cerebellum and cortex (*P*-values = 2.6E-6 and 1.7E-7, respectively, Pearson's Chi-square test). *Post-hoc* tests on the specific mutation types showed that C>A burden was higher in PAXgene tissues than frozen tissues for both regions after Bonferonni correction. The T>C burden was lower in PAXgene tissues for cerebellum (**Fig. S12C**). While statistically significant, we note that the effect sizes are small, e.g., in cerebellum, the median fraction of T>C mutations is still 0 in both frozen and PAXgene samples. Due to the relatively small number of mutations observed in a sample, this analysis may not have had adequate power to detect all differences.

To increase the power of detecting differences between frozen and PAXgene samples, we expanded the analysis to include all tissues in GTEx. This was done to increase the sample size and increase the dynamic range in mutation burden (total and by individual mutation type). However, this approach has the limitation of not being a well-controlled study, i.e., the vast majority of tissues do not have matched frozen and PAXgene tissues. We attempted to account for this by adding relevant covariates into a generalized linear mixed-effects model. For each of the six mutation type burdens and the total mutation burden (N = 7 models), we used Poisson regression to fit the following model:

$$Y \sim preservation\_method + tissue + RIN + AGE + SEX + self\_reported\_ancestry + genotype\_data + transcriptome\_size + sample\_mutation\_power + (1|BATCH) + (1|SUBJECT) \quad (2\text{-}1)$$

where $Y$ was the mutation burden (total or specific mutation type) and $BATCH$ and $SUBJECT$ were modeled as random effects.

We fit the data again with **equation 2-1**, only this time, we dropped the $preservation\_method$ covariate. Models were fit using R package lme4 (v1.1-26)(*59*). Next, we determined if including the preservation method improved the model fit by comparing the Akaike information criterion (AIC) of the nested models. For all seven models, adding $preservation\_method$ did not substantially change the AIC (maximum absolute change in AIC = 2.5%). Therefore, PAXgene does not appear to affect mutation burden and spectra of PZMs.

In summary, while there are systematic differences in preservation method across the GTEx tissues, preservation method does not appear to have a large effect on the mutation burden and spectra of PZMs detected. The small differences detected in the matched brain regions were not replicated when all tissues were examined. This suggests there may be some subtle differences. A large, well-controlled study is needed to determine the extent of these differences.

##### *2.4 Evidence suggesting multi-tissue PZMs occurred prenatally*

Akin to cancer evolution and under the assumption of neutral selection, a PZM that occurs earlier in development will likely be in more tissues and at a higher VAF than a PZM that occurs later. We found a significant positive correlation between VAF and the fraction of the donor's tissues that had the multi-tissue mutation detected (Spearman's $\rho = 0.34$, $P$-value = 9.7E-56, Spearman's rank correlation test, **Fig. S13A**). These results suggest that the multi-tissue mutations occurred prenatally.

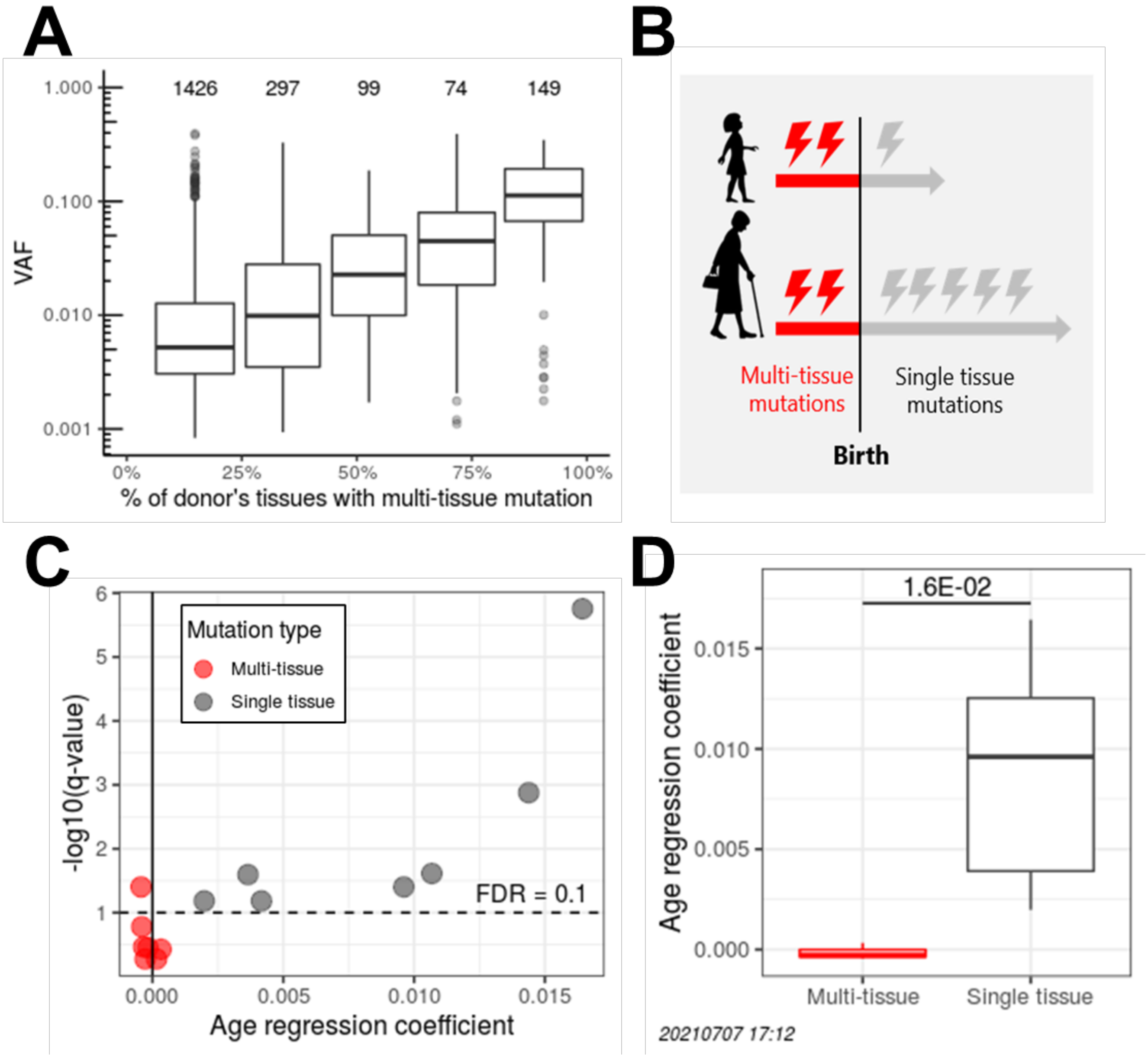

**Fig. S13. Multi-tissue PZMs exhibit prenatal properties.** (**A**) VAF is significantly associated with the fraction of tissues containing multi-tissue PZMs. To mitigate the effect of reduced mutation detection power for tissues with low coverage, the number of tissues in a donor was adjusted to only include tissues with at least 10× coverage at the mutation locus. VAF was defined as the median VAF across all tissues with the mutation. Number of multi-tissue mutations in each boxplot is annotated above the whiskers. (**B**) The number of multi-tissue mutations (a proxy of prenatal PZMs) is expected to be independent of age whereas the number of single tissue mutations (a proxy of postnatal PZMs) is expected to be dependent on age. (C) Prenatal PZM burden is generally not associated with donor age. Significance versus effect size of age regression coefficients by mutation class. Horizontal dashed line at $-\log_{10} 0.1 = 1$ denotes an FDR of 10%. Only tissues with significant single tissue age coefficients and their matched multi-tissue age coefficients are shown (e.g., the significant lung single tissue coefficient and the matching [non-significant] lung multi-tissue age coefficient are plotted). These tissues have an age association with single tissue burden and thus likely have the highest power to detect an age association with multi-tissue burden. (**D**) Distribution of age regression coefficients for multi-tissue and single tissue burden models from (C). *P*-value from Wilcoxon signed-rank test. Top and bottom of boxes denote first and third quartiles, respectively; horizontal black lines denote medians; whiskers denote 1.5× the interquartile range; and outliers are plotted with transparency to disambiguate overlapping values.

Another feature of prenatal mutations is that the number of prenatal mutations in a tissue should be independent of the donor's age since all donors spent approximately the same time *in utero* (**Fig. S13B.**). To show we have power to detect an age association, we also modelled the number of single tissue mutations (a proxy for postnatal mutations) in a tissue. For each tissue $t$ and donor $i$, we modeled the multi-tissue burden as:

$$\frac{\#\ of\ multi\text{-}tissue\ mutations_{t,i}}{\#\ tissues\ profiled\ in\ donor\ i} = \beta_{0,t} + \beta_{1,t}AGE_i + \beta_{2,t}ancestry_i + \beta_{3,t}SEX_i + \beta_{4,t}is\_genotyped_i + \beta_{5,t}donor\_transcriptome\_size_i + \beta_{6,t}sample\_transcriptome\_size_{t,i} + \beta_{7,t}BATCH_{t,i} + \beta_{8,t}RIN_{t,i} + \beta_{9,t}mutation\_detection\_power_{t,i} + \varepsilon_{t,i} \quad (2\text{-}2)$$

A similar model was used for single tissue mutations except that the predictor variable was changed to $\frac{\#\ of\ single\ tissue\ mutations_{t,i}}{\#\ tissues\ profiled\ in\ donor\ i}$ and $SEX$ was dropped for sex-specific tissues.

Consistent with our predictions, age was not significantly associated with multi-tissue mutation burden for the majority of tissues but was significantly associated with single tissue mutation burden for a large number of tissues (**Fig. S13C**). Additionally, the multi-tissue age regression coefficients ($\beta_{1,t}$) were significantly smaller than the single tissue age regression coefficients (*P*-value = 0.016, Wilcoxon signed-rank test) (**Fig. S13D**). Together, these results suggest that multi-tissue mutations are likely prenatal mutations since they have little to no dependence on the donor's age.

###### 2.4.1.1 Prenatal PZMs mapped to earlier timepoints have larger VAFs than later timepoints.

Assuming neutral selection, a PZM that occurs earlier in development will likely have a higher VAF than a PZM that occurs later. Using the reconstructed phylogenies described below, we calculated the distribution of VAFs mapped to each edge of the germ layer tree. The VAFs of mutations that mapped earlier in the tree (parent edge) were larger than the VAFs of mutations that mapped later in the tree (child edges) (*P*-value range 4.5E-41 - 3.8E-5, Mann–Whitney *U* test; **Fig. S14**). These results suggest that the reconstructed phylogenies are correct.

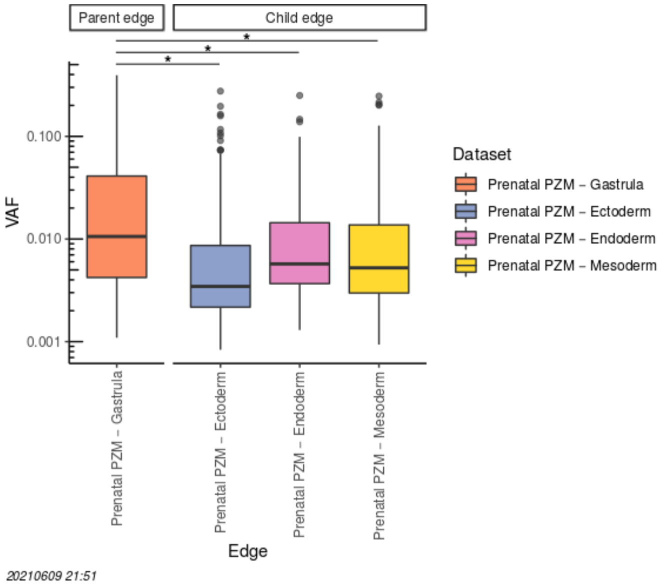

**Fig. S14. PZMs mapped to earlier timepoints have larger VAFs than later timepoints.** Edges are colored by edge name and are sorted parent edge to child edges. "*" denotes statistical significance in VAF between parent and child edges (Mann–Whitney U test). Top and bottom of the boxes denote first and third quartiles, respectively; horizontal black line denotes median; whiskers denote 1.5× the interquartile range; and outliers are plotted with transparency to disambiguate overlapping values.

Additionally, the tree mutation detection power is generally high for mutations with sufficient coverage suggesting that the method is able to map mutations well (**Fig. S15)**.

##### *2.5 PZM phylogeny reconstruction power*

We estimated the power to reconstruct PZM phylogenies via simulation. PZM phylogeny reconstruction power was dependent on the tree topology, profiled tissues, coverage and VAF. Power was greatest when coverage was high and VAF was high (**Fig. S15**). The high variability in power across the tree provides additional evidence that mutation burdens along tree edges should be adjusted for power.

Given that only a subset of GTEx tissues were profiled in a given donor, to examine how GTEx tissue profiling affected mutation mapping power, we also simulated mutations using tissue profiling vectors where all tissues were profiled. These power curves represent the upper bound for detecting mutations in RNA-seq in the GTEx tissues and illustrate that low mapping power was primarily driven by only profiling a subset of tissues in each donor rather than differences in tissue transcriptomes. The power curves may also be a useful reference when designing future experiments to detect prenatal PZMs.

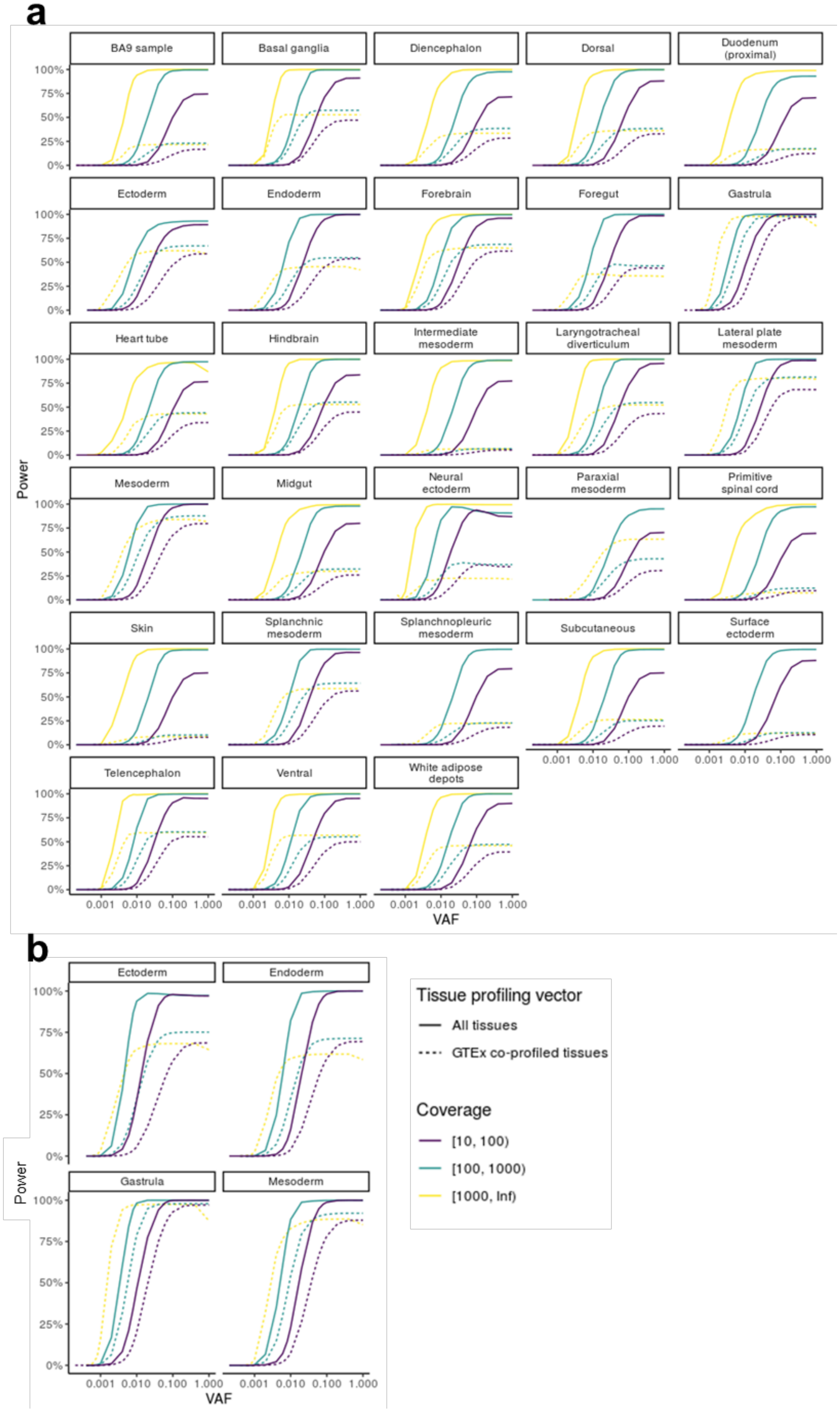
a
BA9 sample
Basal ganglia
Diencephalon
Dorsal
Duodenum (proximal)
Ectoderm
Endoderm
Forebrain
Foregut
Gastrula
Heart tube
Hindbrain
Intermediate mesoderm
Laryngotracheal diverticulum
Lateral plate mesoderm
Mesoderm
Midgut
Neural ectoderm
Paraxial mesoderm
Primitive spinal cord
Skin
Splanchnic mesoderm
Splanchnopleuric mesoderm
Subcutaneous
Surface ectoderm
Telencephalon
Ventral
White adipose depots
Power
VAF
b
Ectoderm
Endoderm
Gastrula
Mesoderm
Power
VAF
Tissue profiling vector
All tissues
GTEx co-profiled tissues
Coverage
[10, 100)
[100, 1000)
[1000, Inf)

**Fig. S15. Edge mutation mapping power is variable.** Edge mutation mapping power depends on tree topology (full tree in **a**, germ layer tree in **b**), edge (facet), tissue co-profiling strategy (line type), coverage (line color), and VAF (x-axis). Edges are referenced by the incident node name. Some facets do not have the highest coverage bin plotted because zero such mutations were encountered during simulation.

##### *2.6 PZM phylogeny validation*

The following lines of evidence suggest that the reconstructed phylogenies are likely correct.

###### *2.6.1 Mapped prenatal PZMs are explained by shared developmental history rather than random sharing patterns*

We confirmed that the observed mapping of the prenatal PZMs could not be explained by random mapping of mutations. If the prenatal PZMs truly did not share developmental histories (and thus were the result from experimental error or multiple independent mutation events), then we would expect the observed edge weights to be no different than those generated from randomized prenatal PZMs catalogs.

The ensemble of observed edge weights was significantly different from random (*P*-value = 2.2E-308, multinomial goodness-of-fit test) and the majority of individual edge weights (56%, 14/25) were significantly different from random after Benjamini-Hochberg correction (permutation tests) (**Fig. S16A**). These results suggest that the multi-tissue PZMs are developmentally acquired mutations and furthermore, the observed edge weights likely estimate the true mutational burdens during development. It is important to note that these are estimates for mutations *that are detectable in adulthood* — the data does not allow for extrapolating to all developmentally acquired mutations since some fraction is likely lost through cell death, revertant mosaicism, etc.

A

B

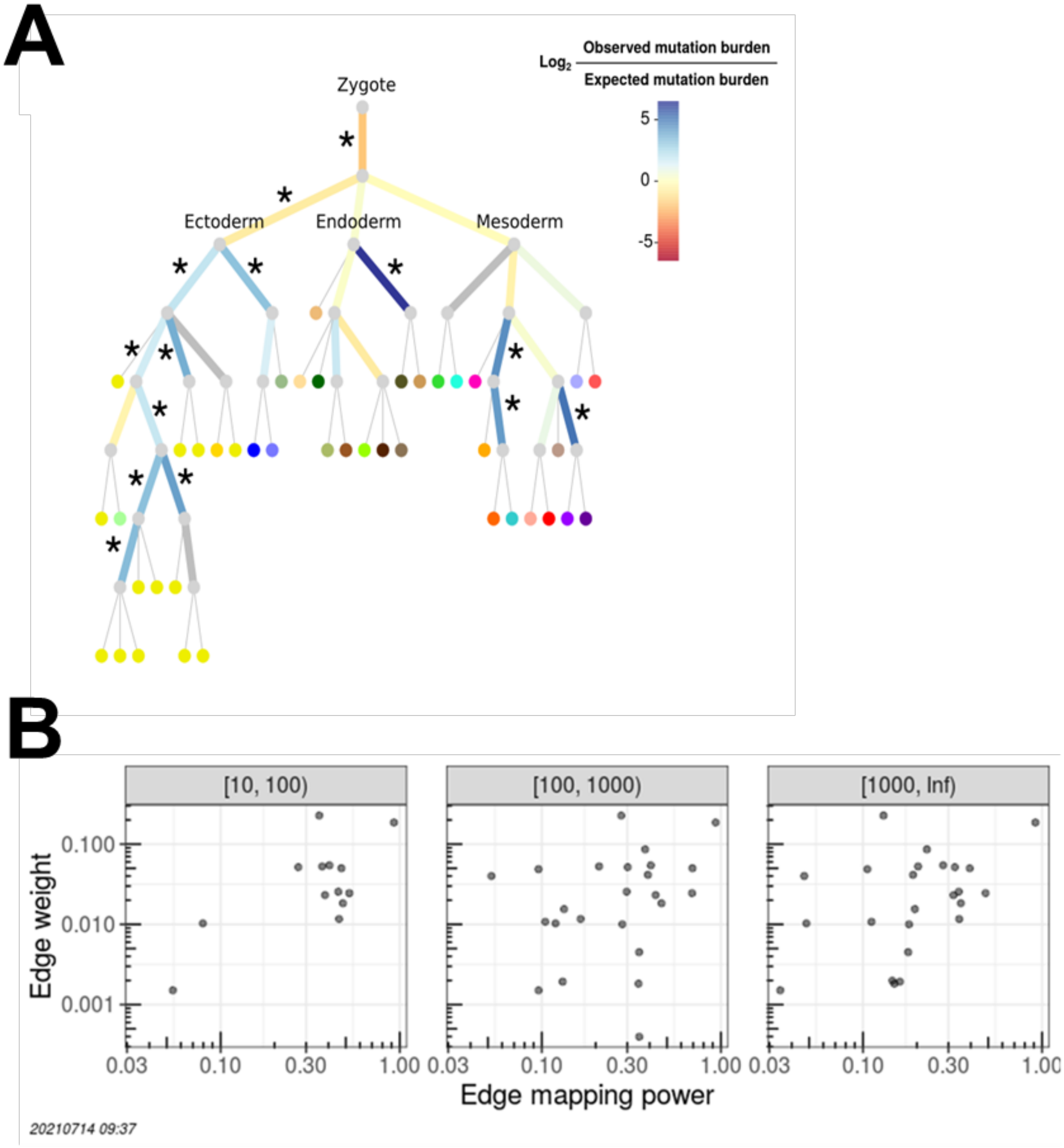

C

| Coverage level | [10, 100) | [100, 1000) | [1000, Inf) |
|---|---|---|---|
| # of edges | 11 | 21 | 21 |
| *P*-value (Spearman's correlation) | 0.37 | 0.09 | 0.12 |

**Fig. S16. Edge weights are nonrandom and not explained by differential power.** (A) Majority of edge weights are significantly different than expected by chance. Edge color represents the $\log_2$ ratio of observed to expected mutation burden. Edges that are significantly different from random are annotated by "*" (q-value $\leq 0.05$). Thick gray edges are edges with limited mutation detection power. See **Fig. S5A** for the full set of vertex labels. GTEx tissue vertices (leaves) are colored using the GTEx coloring convention. (**B**) Observed mutation edge weights are independent of edge mapping power. Since edge mapping power is dependent on coverage, the prenatal mutations were partitioned into three coverage bins. The observed mutation edge weight versus the edge mapping power for mutations with low (**left**), medium (**middle**), and high (**right**) coverage for each edge. (**C**) Results from testing for a correlation between the observed edge weights and the edge mapping power for each coverage level (Spearman's rank correlation test). Only edges with at least 10% edge mapping power were used since edge weights with low edge mapping power may be noisy. The number of edges refers to the number of edges that had the minimum power that were used for the correlation calculation. To calculate an edge mapping power for each edge and coverage level, we first calculated the median VAF of PZMs mapped to that edge and coverage level. Using the edge mapping power curves, we then estimated the power to detect PZMs of that median VAF.

##### *2.6.2 Observed mutation edge weights are independent of edge mapping power*

Since edge mapping power is variable across the tree (see PZM phylogeny reconstruction power), it is important to determine if the observed edge weights can be explained by differential edge mapping power. For both the full tissue tree and germ layer tree, there was no significant correlation between edge weight and edge mapping power at any coverage level (**Fig. S16B** and **C**). These results imply that edge weights are independent of ascertainment bias.

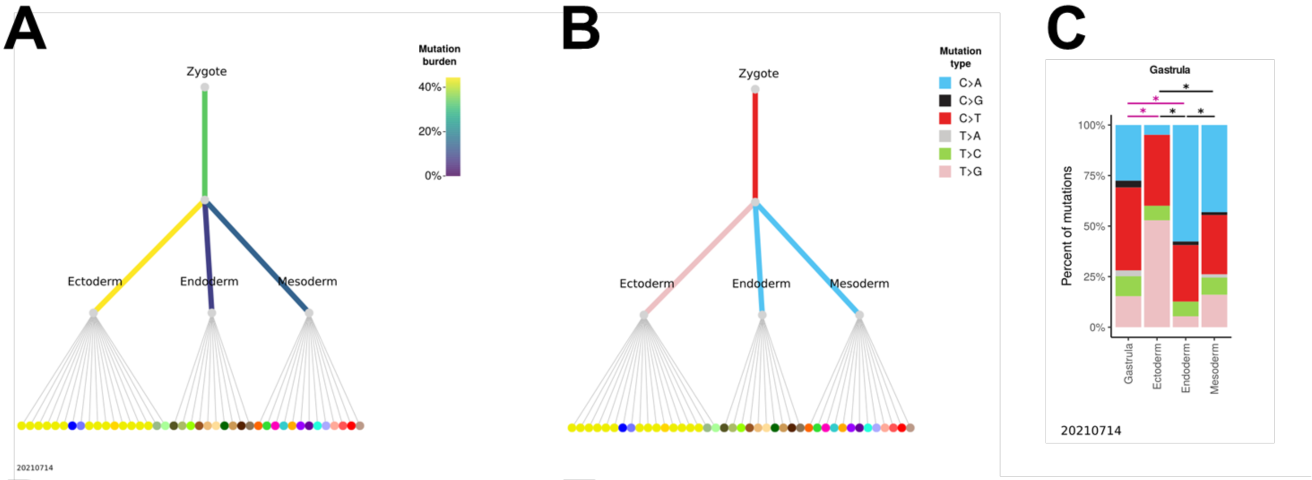

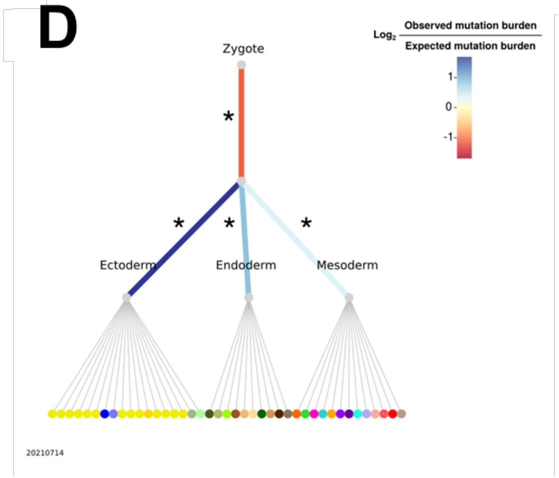

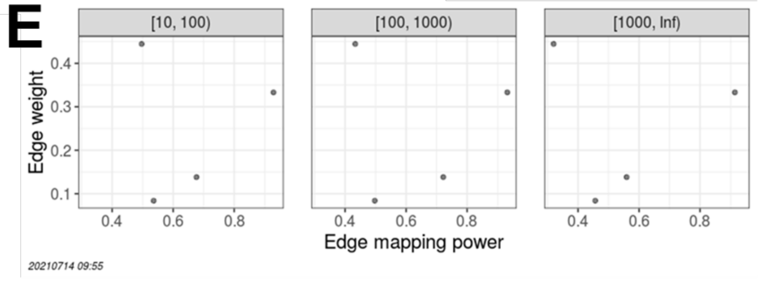

**F**

| Coverage level | [10, 100) | [100, 1000) | [1000, Inf) |
|---|---|---|---|
| # of edges | 4 | 4 | 4 |
| *P*-value (Spearman's correlation) | 0.92 | 0.92 | 0.92 |

**Fig. S17. Characterization of PZMs mapped to germ layer tree.** (**A**) Prenatal PZM mutation burden. Edge color represents the percent of prenatal PZMs mapped to that period in development. (**B**) The predominant mutation type. Edge color represents the predominant mutation type of mutations mapped to that edge. (**C**) Local variation in mutation spectra across developmental space and time. Facet represents the mutation spectra observed in a parent edge (leftmost barplot) and children edges. Statistically significant differences in mutation spectra are annotated with "*". (**D**) All edge weights are significantly different than expected by chance. Edge color represents the $\log_2$ ratio of observed to expected mutation burden. Edges that are significantly different from random are annotated by "*" (q-value $\leq 0.05$). See **Fig. S5B** for the full set of vertex labels. GTEx tissue vertices (leaves) are colored using the GTEx coloring convention. (**E**) Observed mutation edge weights are independent of edge mapping power. The observed mutation edge weight versus the edge mapping power for mutations with low (**left**), medium (**middle**), and high (**right**) coverage for each edge. (**F**) Results from testing for a correlation between the observed edge weights and the edge mapping power for each coverage level (Spearman's rank correlation test). The number of edges refers to the number of edges that had the minimum power that were used for the correlation calculation.

#### *2.7 Relationship between VAF and deleteriousness score*

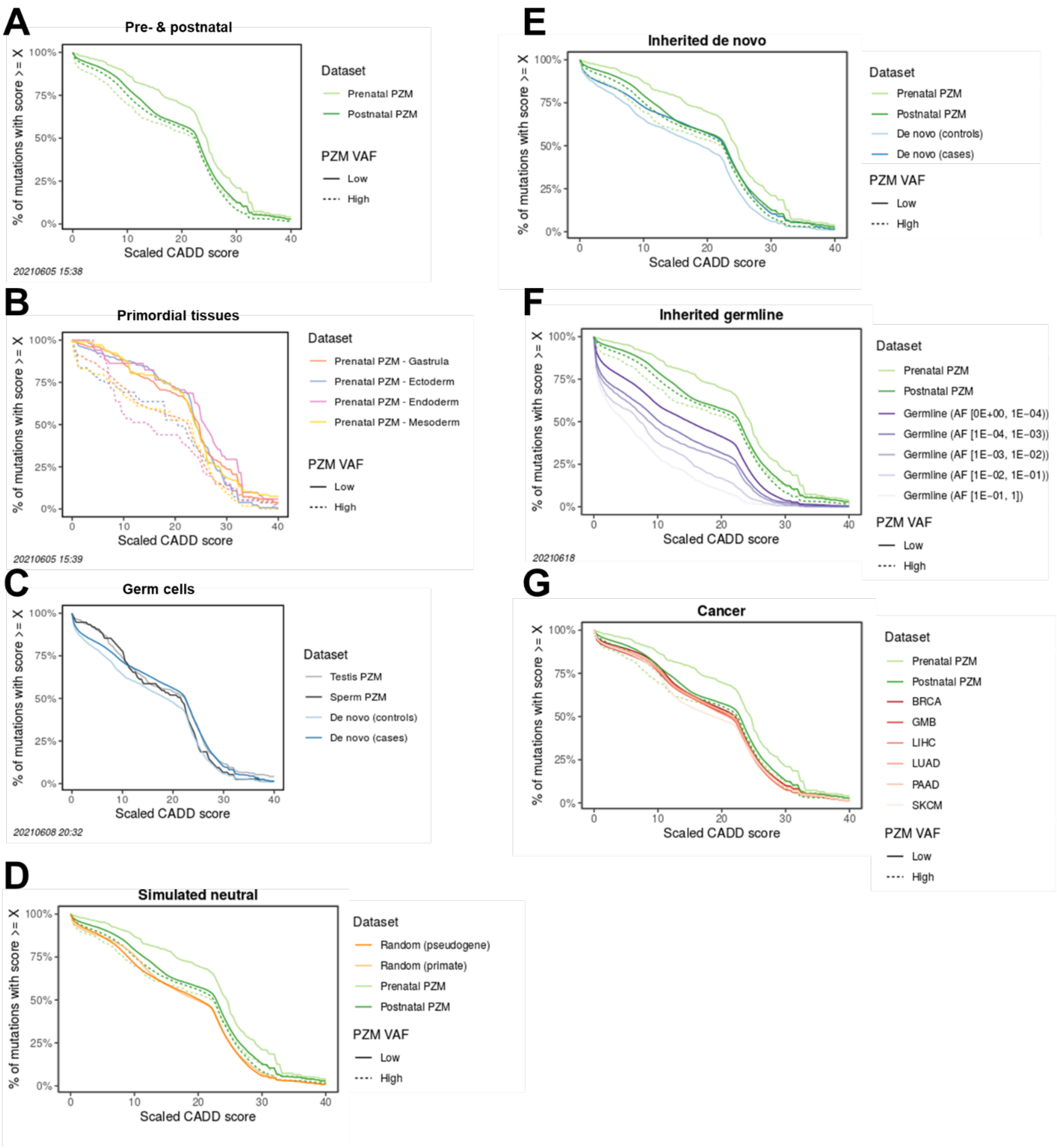

**Fig. S18. Low VAF prenatal PZMs are the most deleterious class of PZMs.** (**A-G**) PHRED-scaled CADD score CCDFs as a function of VAF, space, time, and classes of genetic variation. Distributions with larger areas under the curve have a larger fraction of deleterious mutations.

(**A**) Prenatal and postnatal PZMs stratified by VAF. (**B**) Prenatal PZMs by primordial tissue and VAF. (**C**) Mutations throughout the germ cell life cycle. (**D**) PZMs versus simulated neutral mutations, (**E**) inherited de novo mutations, (**F**) inherited germline variant, and (**G**) somatic mutations in cancer. To increase readability of the comparisons, the same PZM data is plotted in A and D-G.

##### *2.8 The high fraction of deleterious PZM mutations is recapitulated in multiple validation datasets*

To determine if the PZM deleteriousness results were generalizable, we compared the deleteriousness of GTEx PZMs with PZMs from four other datasets (TwinsUK (unpublished work), García-Nieto *et al.*(*50*), Yizhak *et al.*(*49*), and Brazhnik *et al.*(*51*)). Importantly, these validation datasets used a variety of different data sources, nucleic acid types, and variant calling algorithms (see

| Tissue | Color |
|---|---|
| Adipose - Subcutaneous | |
| Adipose - Visceral (Omentum) | |
| Adrenal Gland | |
| Artery - Aorta | |
| Artery - Coronary | |
| Artery - Tibial | |
| Brain - Amygdala | |
| Brain - Anterior cingulate cortex (BA24) | |
| Brain - Caudate (basal ganglia) | |
| Brain - Cerebellar Hemisphere | |
| Brain - Cerebellum | |
| Brain - Cortex | |
| Brain - Frontal Cortex (BA9) | |
| Brain - Hippocampus | |
| Brain - Hypothalamus | |
| Brain - Nucleus accumbens (basal ganglia) | |
| Brain - Putamen (basal ganglia) | |
| Brain - Spinal cord (cervical c-1) | |
| Brain - Substantia nigra | |
| Breast - Mammary Tissue | |
| Cells - EBV-transformed lymphocytes | |
| Cells - Cultured fibroblasts | |
| Colon - Sigmoid | |
| Colon - Transverse | |
| Esophagus - Gastroesophageal Junction | |
| Esophagus - Mucosa | |
| Esophagus - Muscularis | |
| Heart - Atrial Appendage | |
| Heart - Left Ventricle | |
| Kidney - Cortex | |
| Liver | |
| Lung | |
| Minor Salivary Gland | |
| Muscle - Skeletal | |
| Nerve - Tibial | |
| Ovary | |
| Pancreas | |
| Pituitary | |
| Prostate | |
| Skin - Not Sun Exposed (Suprapubic) | |
| Skin - Sun Exposed (Lower leg) | |
| Small Intestine - Terminal Ileum | |
| Stomach | |
| Testis | |
| Thyroid | |
| Uterus | |
| Vagina | |
| Whole Blood | |

**Table S9. U.S. cancer incidence rates related to tissues with significant differences in mutation burden among ancestry groups.**

**Table S10. Summary of datasets used to validate deleteriousness patterns and whether results were validated**.NA = Not applicable.

| | **This study** | **TwinsUK** | **García-Nieto *et al.*** | **Yizhak *et al.*** | **Brazhnik *et al.*** |
|---|---|---|---|---|---|
| **Reference** | NA | Unpublished | (*50*) | (*49*) | (*51*) |
| **Uses RNA** | Yes | Yes | Yes | Yes | No |
| **Uses GTEx data** | Yes (v8) | No | Yes (v7) | Yes (v7) | No |
| **Variant calling method** | Lachesis | Lachesis | Custom method | RNA-MuTect | VarScan2, MuTect2, & HaplotypeCaller |
| **Results validated** | NA | Yes | No | Yes | Yes |

**Table S11. List of CHIP mutations detected in GTEx samples.**Genomic coordinates are relative to GRCh38 and are 0-based with exclusive end coordinates. GTEx recurrence is defined as the number of times the mutation was detected in GTEx blood and LCL samples. COSMIC recurrence is defined as the number of times the mutation was detected in haematopoietic and lymphoid tissue cancer samples. COSMIC recurrence percentile is defined as the percent of recurrent COSMIC mutations with the same or lower recurrence as the mutation.

| chr | start | end | ref_allele | alt_allele | gene_name | HGVSG | HGVSC | HGVSP | cosmic_mutation_id | tissue | recurrence.gtex | recurrence.cosmic | recurrence_percentile.cosmic |
|---|---|---|---|---|---|---|---|---|---|---|---|---|---|
| chr3 | 38141149 | 38141150 | T | C | MYD88 | 3:g.38141150T>C | ENST00000417037.6:c.818T>C | ENSP00000401399.2:p.Leu273Pro | COSV57169334 | Cells - EBV-transformed lymphocytes | 1 | 1969 | 0.99998 |
| chr15 | 90088701 | 90088702 | C | T | IDH2 | 15:g.90088702C>T | ENST00000330062.7:c.419G>A | ENSP00000331897.3:p.Arg140Gln | COSV57468751 | Cells - EBV-transformed lymphocytes | 1 | 815 | 0.99991 |
| chr17 | 42322442 | 42322443 | T | A | STAT3 | 17:g.42322443T>A | ENST00000264657.9:c.1940A>T | ENSP00000264657.4:p.Asn647Ile | COSV52882818 | Cells - EBV-transformed lymphocytes | 1 | 18 | 0.99630 |
| chr12 | 92145426 | 92145427 | G | A | BTG1 | 12:g.92145427G>A | ENST00000256015.4:c.109C>T | ENSP00000256015.3:p.Leu37= | COSV55458204 | Cells - EBV-transformed lymphocytes | 2 | 6 | 0.98268 |
| chr15 | 44711580 | 44711581 | T | C | B2M | 15:g.44711581T>C | ENST00000558401.5:c.35T>C | ENSP00000452780.1:p.Leu12Pro | COSV62563197 | Cells - EBV-transformed lymphocytes | 1 | 6 | 0.98268 |
| chr17 | 42322400 | 42322401 | T | A | STAT3 | 17:g.42322401T>A | ENST00000264657.9:c.1982A>T | ENSP00000264657.4:p.Asp661Val | COSV52886283 | Whole Blood | 1 | 6 | 0.98268 |

for summary). A detailed description of the validation results are in the following sections. Briefly, our findings regarding the frequency of deleterious PZM mutations were supported by three of the four validation datasets (**Fig. S20**, **Fig. S21**, **Fig. S22**, **Fig. S19**). Deleteriousness patterns observed in García-Nieto *et al.* were dissimilar to all other datasets. We suspect this may be due to germline variant contamination and not biological differences in deleteriousness.

Therefore, we have high confidence that the deleteriousness observations described herein may reflect global properties of PZMs rather than technical artifacts.

##### *2.8.1 Brazhnik et al. validation dataset*

To establish that the CADD score distributions derived from RNA-seq PZM calls were similar to PZMs inferred from DNA-seq, we compared the deleteriousness of our liver PZMs with PZMs from DNA-seq generated by Brazhnik *et al.*(*51*) Brazhnik *et al.* detected PZMs from single human hepatocytes, liver stem cells (LSCs), and organoids derived from LSCs. Importantly, these PZMs were detected from samples independent of GTEx, were detected in DNA, and used a different mutation calling method and thus, represents an independent validation dataset. Hepatocytes are an appropriate validation cell type because bulk liver is primarily composed of hepatocytes(*69*). Thus the comparison between single-cell hepatocytes and bulk liver data will likely not be confounded by cell composition issues.

PZMs were detected from 62 DNA single-cell multiple displacement amplification libraries across 12 donors. After filtering PZMs to the allowable transcriptome, there were 414 PZMs of which the majority were from hepatocytes (88% (363/414)). Since all samples were generated from the same tissue, we did not try to predict prenatal PZMs. Additionally, since PZMs were generated from single-cell data rather than bulk data, we did not try to predict the VAF of the PZM in bulk tissue.

Hepatocyte PZMs were as deleterious as GTEx postnatal liver PZMs (odds-ratio = 0.84, FDR-corrected q-value = 0.21). LSC PZMs were also as deleterious as postnatal liver PZMs (odds-ratio = 1.8, FDR-corrected q-value = 0.09). Surprisingly, LSC PZMs may be less deleterious than hepatocyte PZMs; however this result was not significant after multiple test correction (odds-ratio = 0.46, *P*-value = 0.014, q-value = 0.08) (**Fig. S19**). The lack of a significant difference may be due to small sample size. This observation raises an intriguing hypothesis that stem cells may be under stronger selection than differentiated cells to constrain the functional impact of PZMs.

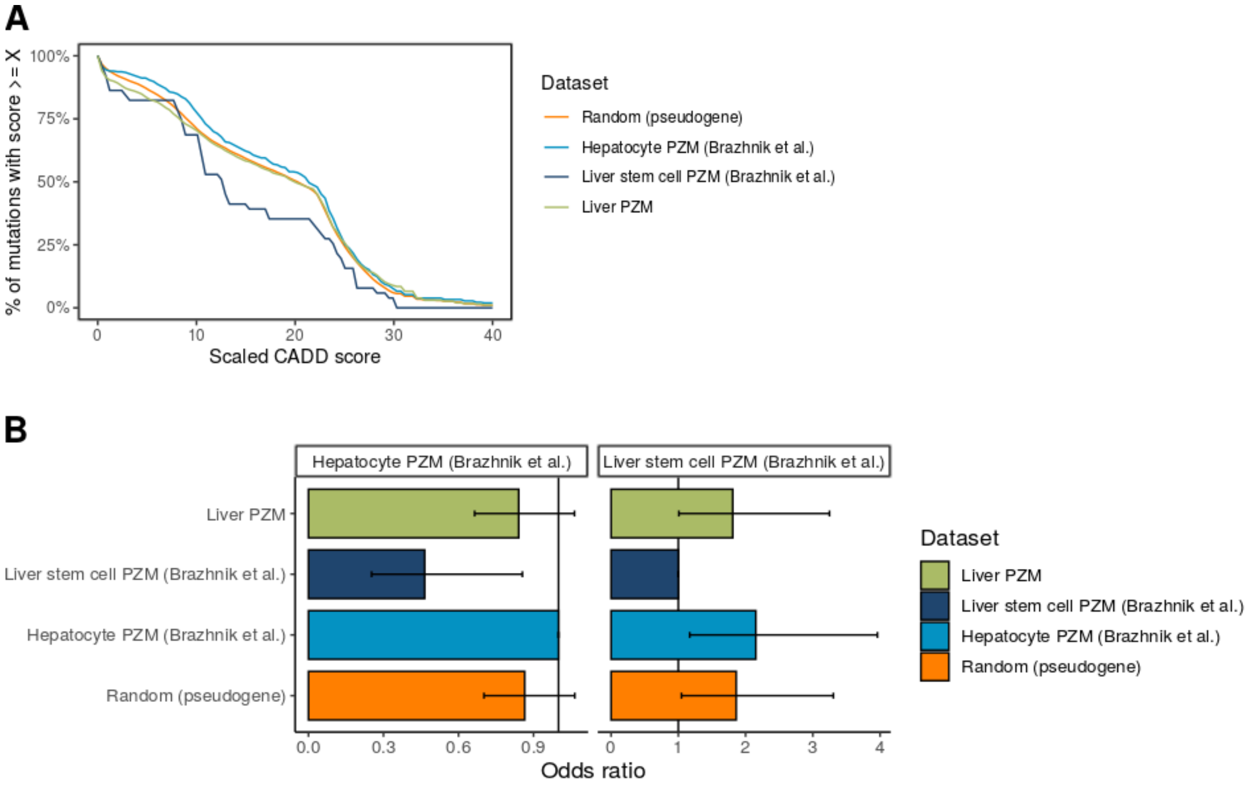

**Fig. S19. PZM deleteriousness results replicated in Brazhnik *et al.* data.** (**A**) PHRED-scaled CADD score CCDFs of GTEx PZMs and Brazhnik *et al*. PZMs. Random neutral CADD CCDF shown for context. (**B**) Odds of detecting deleterious mutations in a given dataset compared to Brazhnik *et al.* hepatocyte PZMs (**left**) and LSC PZMs (**right**). Vertical line at odds ratio = 1 indicates no difference in odds of detecting deleterious mutations in a given dataset compared to reference dataset. Error bars represent 95% CIs.

##### *2.8.2 TwinsUK validation dataset*

As another form of validation, we ran our variant calling method on a novel, independent dataset, TwinsUK(*47*). The TwinsUK dataset consisted of data from 856 twins wherein each twin had ~3 tissues profiled with RNA-seq. In total, there were ~2,500 RNA-seq samples. After filtering PZMs to the allowable transcriptome, there were 1,507 PZMs of which 1% (18/1,507) were predicted to be prenatal PZMs. (Of note, compared to GTEx, TwinsUK had lower sequencing depth and fewer tissues profiled per donor. As a result, the sensitivity to detect low VAFs and prenatal PZMs was most likely reduced.)

Encouragingly, the deleteriousness of TwinsUK PZMs was not statistically different than the GTEx PZMs, suggesting that the GTEx results may reflect global properties of PZMs rather than technical artifacts (odds ratio of GTEx prenatal PZMs over TwinsUK prenatal PZMs = 1.14, FDR-corrected q-value = 0.84 (GTEx low VAF) and odds ratio = 0.57, q-value = 0.37 (GTEx high VAF); odds ratio of GTEx postnatal PZMs over TwinsUK postnatal PZMs = 0.98, q-value = 0.84) (**Fig. S20**).

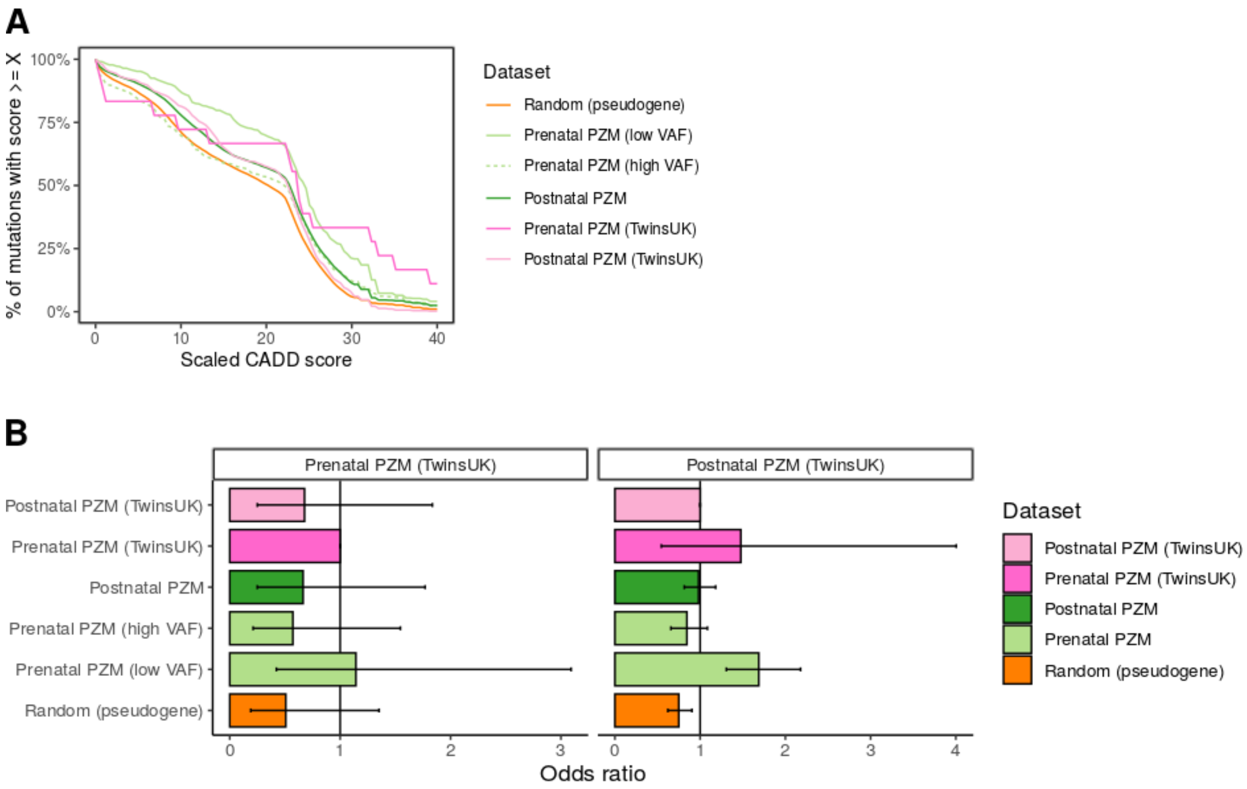

**Fig. S20. PZM deleteriousness results are replicated in TwinsUK data.** (**A**) PHRED-scaled CADD score CCDFs of GTEx PZMs and TwinsUK PZMs. TwinsUK PZMs were not partitioned by VAF due to the small sample size and decreased sensitivity to detect low VAF PZMs. Random neutral CADD CCDF shown for context. (**B**) Odds of detecting deleterious mutations in a given dataset compared to TwinsUK prenatal PZMs (**left**) and postnatal PZMs (**right**). Vertical line at odds ratio = 1 indicates no difference in odds of detecting deleterious mutations in a given dataset compared to reference dataset. Error bars represent 95% CIs.

##### *2.8.3 Yizhak et al. validation dataset*

We next compared the deleteriousness of PZMs with PZMs from Yizhak *et al.*(*49*) These PZMs were generated from an earlier version of GTEx. Thus, this validation dataset is not entirely independent from the one in this study. However, a different mutation calling method was used to call PZMs from RNA-seq. PZMs were detected from ~6,700 RNA-seq samples across 488 donors and 29 tissue types. After filtering PZMs to the GTEx pass QC tissues and allowable transcriptome, there were 4,454 PZMs, of which we predicted 0.8% (36/4,454) to be prenatal. All VAFs were larger than the low VAF cutoff and thus were defined as high VAF PZMs.

Yizhak *et al.* high VAF prenatal PZMs were as deleterious as prenatal high VAF PZMs (odds-ratio = 1.8, FDR-corrected q-value = 0.13). Yizhak *et al.* postnatal PZMs were slightly less deleterious than postnatal PZMs (odds-ratio = 1.2, q-value = 1.5E-8) (**Fig. S21A** and **B**). However, this difference may be due to differences in tissue composition between the different GTEx versions. Therefore, we compared deleteriousness of postnatal PZMs by tissue.

Due to the lower dataset size, we only compared tissues with at least 100 PZMs in Yizhak *et al.* Additionally, we used the same tissue grouping as Yizhak *et al.*, e.g., skin includes GTEx tissues Skin - Not Sun Exposed (Suprapubic) and Skin - Sun Exposed (Lower leg). Yizhak *et al.* tissue-specific postnatal PZMs were as deleterious as tissue-specific postnatal PZMs for 80% (8/10) of tissues. The exceptions were brain (odds-ratio = 0.62, q-value = 5.9E-3) and skin (odds-ratio =

0.87, q-value = 0.04) (**Fig. S21C** and **D**). The differences may be due to differences in tissue composition across the two datasets since the tissues represent multiple GTEx tissues.

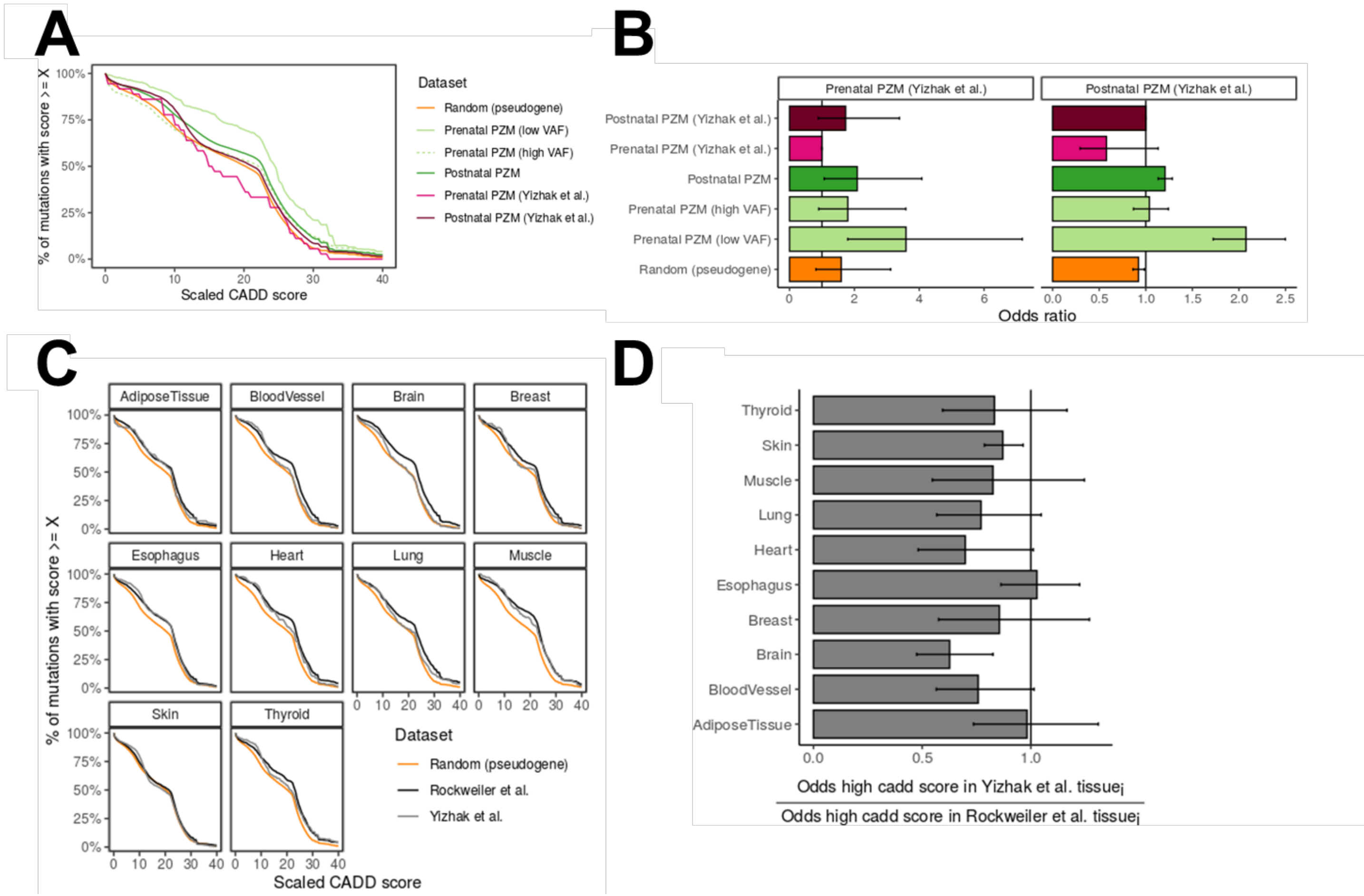

**Fig. S21. Global and tissue-specific PZM deleteriousness results replicated in Yizhak *et al.* data** (**A**) PHRED-scaled CADD score CCDFs of GTEx PZMs and Yizhak *et al.* Random neutral CADD CCDF shown for context. (**B**) Odds of detecting deleterious mutations in a given dataset compared to Yizhak *et al.* [high VAF] prenatal PZMs (**left**) and postnatal PZMs (**right**). (**C**) PHRED-scaled CADD score CCDFs of GTEx postnatal PZMs and Yizhak *et al.* postnatal PZMs by tissue. Random neutral CADD CCDF shown for context. (**D**) Odds of detecting deleterious mutations in a given tissue in Yizhak *et al.* relative to the odds of detecting deleterious mutations in the same tissue in this study. Vertical lines at odds ratio = 1 indicates no difference in odds of detecting deleterious mutations in a given dataset compared to reference dataset. Error bars represent 95% CIs.

##### *2.8.4 García-Nieto et al. validation dataset*

We next compared the deleteriousness of PZMs with PZMs from García-Nieto *et al.*(*50*) Like Yizhak *et al.*, these PZMs were generated from an earlier version of GTEx and thus, this validation dataset is not entirely independent from the one in this study. However, a different mutation calling method was used to call PZMs from RNA-seq. PZMs were detected from ~7,600 RNA-seq samples across 547 donors and 36 tissue types. After filtering PZMs to the GTEx pass QC tissues and allowable transcriptome, there were 282,561 PZMs, of which we predicted 9.6% (27,251/282,561) to be prenatal. All VAFs were larger than the low VAF cutoff and thus were defined as high VAF PZMs.

Surprisingly, García-Nieto *et al.* prenatal and postnatal PZMs were as deleterious as common standing germline variation (population allele frequency = 1-10%, odds-ratio = 1.2, FDR-corrected q-value = 0.17) and did not validate the patterns observed with the PZMs presented here and the other validation studies (**Fig. S22**). Given the low fraction of deleterious mutations, their similarity to germline mutations, and their dissimilarity to the other four datasets (the PZMs presented here, TwinsUK, Yizhak *et al.*, and Brazhniek *et al.*), we suspect this dataset may be contaminated with common germline variants from the donors.

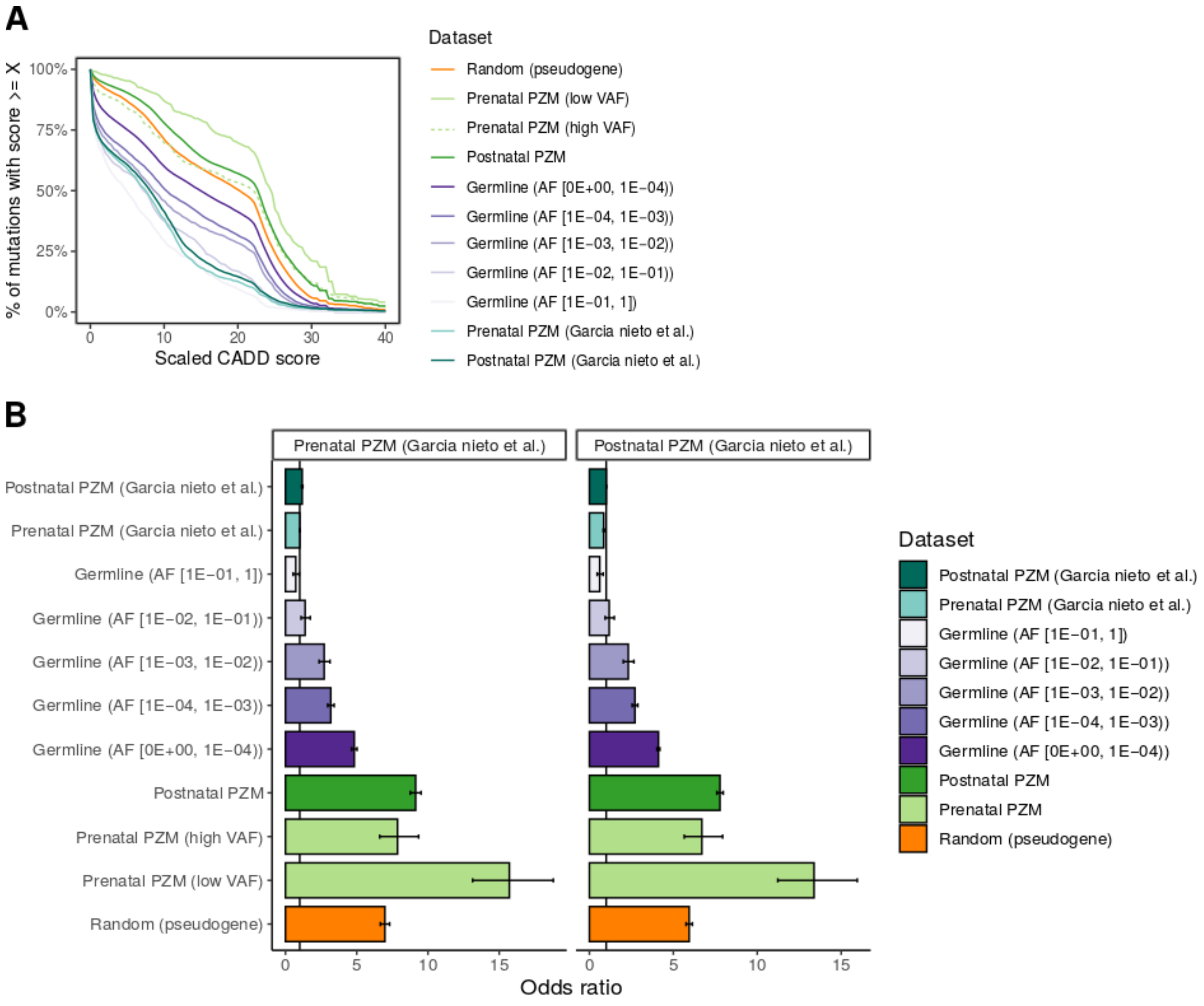

**Fig. S22. PZM deleteriousness results not replicated in García-Nieto *et al.* data.**

(**A**) PHRED-scaled CADD score CCDFs of GTEx PZMs and García-Nieto *et al*. PZMs. Random neutral CADD CCDF and germline variation CCDFs shown for context. (**B**) Odds of detecting deleterious mutations in a given dataset compared to García-Nieto *et al.* prenatal [high VAF] PZMs (left) and postnatal PZMs (**right**). Vertical line at odds ratio = 1 indicates no difference in odds of detecting deleterious mutations in a given dataset compared to reference dataset. Error bars represent 95% CIs. AF = population allele frequency.

***Prevalence of expressed CHIP mutations in GTEx***

Clonal growth of hematopoietic cells (regardless of the underlying cause or disease state) is known as clonal hematopoiesis. Clonal hematopoiesis of indeterminate potential (CHIP) is a clinical subclass in which cancer-associated variants (most often in *ASXL1*, *DNMT3A*, and *TET2*) are detected in the blood of individuals without apparent hematologic malignancies(*63*). Since larger clones are more likely to be clinically meaningful, the field generally requires CHIP mutations to have a VAF, a measure of clone size, to be ≥ 0.02(*63*). While CHIP mutations are rare within a donor, CHIP appears to be a common phenomenon across donors. Detection of CHIP increases with age: CHIP is detected in < 1% of individuals younger than 50 but is found in 10% of individuals older than 65(*64*).

##### *2.9.1 Definition of CHIP mutations*

We defined putative **CHIP mutations** as single point mutations that were observed ≥ 5× in haematopoietic and lymphoid cancer samples in COSMIC v92 (N = 2,289 variants at 2,076 genomic sites).

##### *2.9.2 Expressed CHIP mutations are observed at expected rates*

We asked whether CHIP could be detected in the GTEx donors. The motivation was threefold. First, to provide additional evidence about the validity of the variant calling method. Second, unlike other large cohort studies that use exome and targeted sequencing with limited coverage, RNA sequencing offers the possibility of thousands of fold coverage and thus the potential to detect very low VAFs. Such results would expand the field's knowledge about the prevalence of CHIP at sub-clinical VAFs. Lastly, detecting mutations in RNA versus the routinely used DNA, would offer new insight on the potential function of mutated genes.

To detect CHIP mutations in GTEx, we scanned the PZMs detected in whole blood (N = 746 samples) and EBV-transformed lymphocytes (N = 174 samples) for overlap with the set of putative CHIP mutations. Six unique CHIP mutations were detected in 7 samples (Table **S9. U.S.**

**cancer incidence rates related to tissues with significant differences in mutation burden among ancestry groups.**

**Table S10. Summary of datasets used to validate deleteriousness patterns and whether results were validated**.NA = Not applicable.

| | **This study** | **TwinsUK** | **García-Nieto *et al.*** | **Yizhak *et al.*** | **Brazhnik *et al.*** |
|---|---|---|---|---|---|
| **Reference** | NA | Unpublished | (*50*) | (*49*) | (*51*) |
| **Uses RNA** | Yes | Yes | Yes | Yes | No |
| **Uses GTEx data** | Yes (v8) | No | Yes (v7) | Yes (v7) | No |
| **Variant calling method** | Lachesis | Lachesis | Custom method | RNA-MuTect | VarScan2, MuTect2, & HaplotypeCaller |
| **Results validated** | NA | Yes | No | Yes | Yes |

**Table S11**). Two of the mutations (IDH2 R140Q and MYD88 L273P) are in the 99.99$^{th}$ percentile of recurrent mutations in haematopoietic and lymphoid cancers and have been shown to have gain-of-function properties(*70*, *71*). 0.1% (1/746) of whole blood donors and 3.5% (6/174) of EBV-transformed lymphocyte donors had a CHIP mutation. Of note, none of the CHIP-positive donors had a history of non-metastatic cancer. The observed CHIP prevalence in GTEx is similar to what we would expect given the age demographics of the cohort and published prevalence rates(*64*).

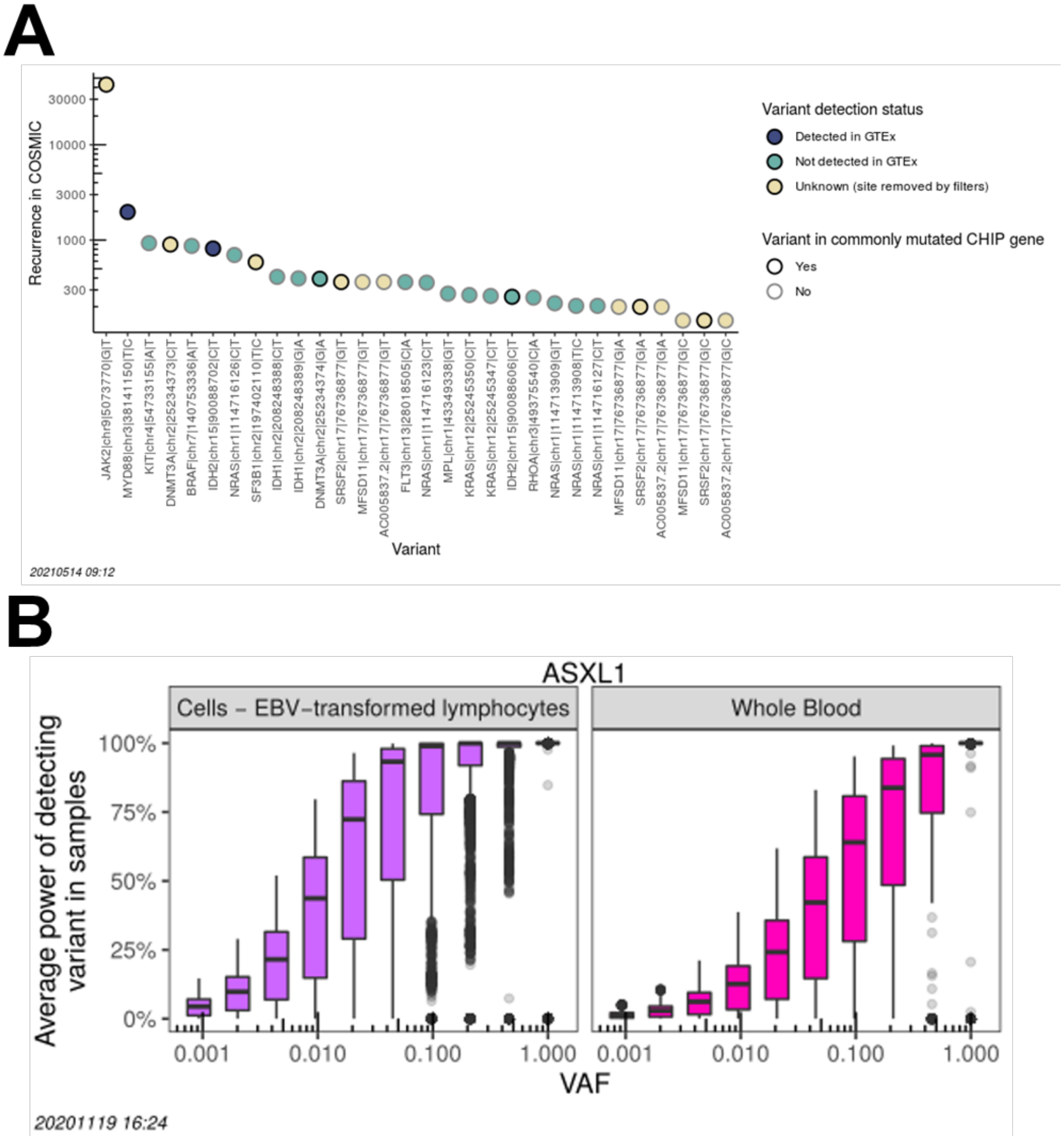

**Fig. S23. Filtering and low power reduced sensitivity to detect CHIP mutations.** (**A**) Several commonly observed mutations in haematopoietic and lymphoid cancers and CHIP studies were excluded in this study.For the top 30 recurrent mutations observed in haematopoietic and lymphoid cancers, we annotated whether the genomic position was removed by the mutation calling method (yellow) or retained (blue and teal) and plotted their recurrence in cancer from highest to lowest. Genes commonly mutated in CHIP studies are outlined in black; grey otherwise. (**B**) Mutation detection power in CHIP genes is low and is greater in EBV-transformed lymphocytes than whole blood. ASXL1 is shown as a representative CHIP gene. Distribution of average power to detect a variant in ASXL1 across all samples as a function of VAF in EBV-transformed lymphocytes (**left**) and whole blood (**right**). Each point in a boxplot represents the average power across all samples at a given position in the gene. Tissues colored in their canonical GTEx colors. Top and bottom of the boxes denote first and third quartiles, respectively; horizontal black lines denote median; whiskers denote 1.5× the interquartile range; and outliers are plotted with transparency to disambiguate overlapping values.

We strongly caution that the CHIP prevalence and CHIP burden are likely underestimates. Since we valued specificity over sensitivity during variant calling, the filters used were very aggressive. As a result, 67% (1,534/2,289) of the defined CHIP mutations were removed by one or more filters. This list includes DNMT3A R882H, a highly recurrent mutation observed in CHIP studies(*64*, *72*) and JAK2 V617F, the most recurrent mutation observed in haematopoietic and lymphoid cancers(*73*). A list of the most recurrent CHIP mutations and their filter status is visualized in **Fig. S23A**.

In addition to the constraints imposed by our variant calling method, we were also constrained by detection power stemming from the expression and background error rate. To calculate the PZM detection power for a given sample $s$, at position $i$, with PZM VAF $v$, we set up the hypothesis test as:

$$H_0: alt.coverage_{s,i} \sim Binom(p = error\ rate_i, N = total\ coverage_{s,i}) \quad \text{2-3}$$
$$H_a: alt.coverage_{s,i} \sim Binom(p = v, N = total\ coverage_{s,i})$$

and defined the statistical power as $1 - \Pr\ (Type\ II\ error)$. We next calculated the average power across all samples at a given position for each position in genes associated with CHIP. The detection power for a representative CHIP gene is shown in **Fig. S23B**. For many CHIP genes, detection power was very limited. This likely contributed to underestimating the true CHIP prevalence. Additionally, we are underpowered by biology and study design to detect several classes of CHIP mutations. Since mutations are detected at the RNA level, nonsense mutations which can contribute a large fraction of single nucleotide CHIP mutations (e.g., 27% in (*72*)) may be undetected due to nonsense-mediated decay. To avoid false positives from spurious alignments at splice junctions, we ignored all mutations near splice junctions. Lastly, we did not call postzygotic indels.

##### *2.9.3 Elevated rates of CHIP mutations in EBV-transformed lymphocytes compared to blood is likely due to technical reasons*

The enrichment of detected CHIP mutations in EBV-transformed lymphocytes compared to whole blood may be the result of mutations occurring *in vitro* rather than *in vivo.* While we cannot rule out this possibility, we did observe that EBV-transformed lymphocytes samples had greater mutation detection power than whole blood samples at CHIP genes due to the differences in expression levels (**Fig. S23B**). Thus for technical reasons, we would expect EBV-transformed lymphocytes to be enriched for CHIP mutations compared to whole blood.

##### *2.9.4 Summary*

In summary, LachesisDetect was able to detect expressed somatic cancer drivers in individuals without apparent hematological malignancies. This result provides additional evidence that the mutation calling method has high specificity and moderate sensitivity. To our knowledge, this is the first time CHIP mutations have been detected at the RNA level and thus suggest expression of these mutations may have a functional role in clonal growth.

#### ***2.10 Characterization of germ cell PZMs***

##### *2.10.1 Germ cell PZMs can be detected in bulk male gonads*

We hypothesized that PZMs detected in bulk gonads could be used to study PZMs in germ cells. We used a bottom-up approach by first investigating if the transcriptomes of germ cells and bulk gonads were similar and then determining if the specific genes with PZMs were enriched for genes expressed in germ cells.

We first determined if germ cell PZMs could be detected in bulk female gonads. We tested if the bulk ovary transcriptome was similar to the oocyte transcriptome. We performed hierarchical clustering of published human mature oocyte transcriptomes(*74*) with GTEx tissue transcriptomes. The oocyte transcriptomes did not closely cluster with the ovary transcriptome, suggesting that bulk ovary expression is a poor proxy for female germ cell expression (**Fig. S24A**). This result is expected given that ovaries primarily consist of interstitial stroma, a heterogenous mixture of somatic cell types(*75*). We therefore concluded that PZMs in bulk ovary could not be confidently mapped to PZMs in female germ cells.

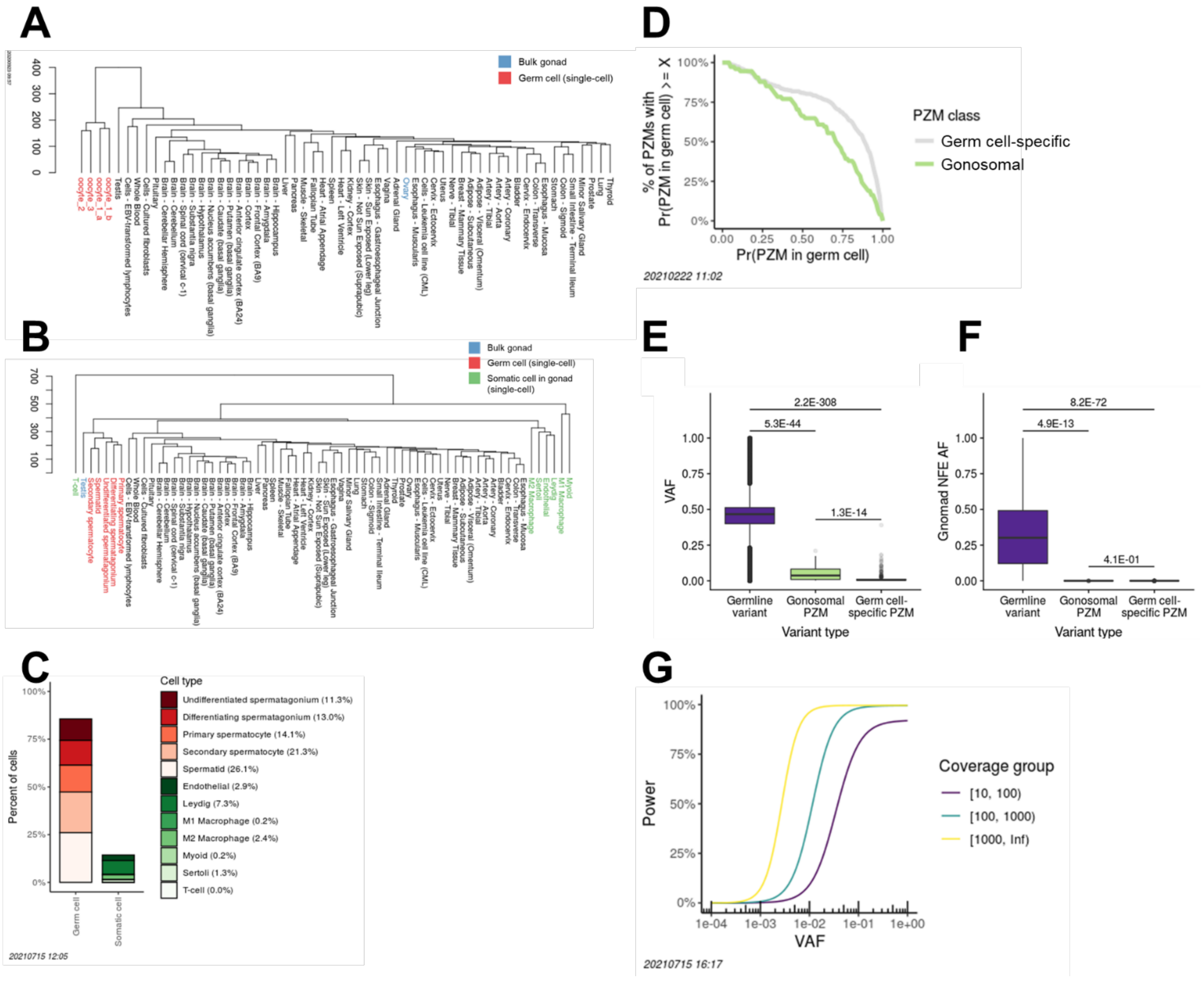

**Fig. S24. Germ cell PZMs can be detected in bulk male gonads.** (**A**) Bulk ovary and single-cell oocyte transcriptomes are dissimilar. Data from single-cell RNA-seq of oocytes (red) is from (*74*). Oocyte data consists of three biological replicates (oocyte_1, oocyte_2, and oocyte_3). Oocyte_1 had two technical replicates (oocyte_1_a and oocyte_1_b). While oocytes form a clade, oocytes and bulk ovary (blue) do not, suggesting that oocyte and ovary transcriptomes are dissimilar. (**B**) Bulk testis and male germ cell transcriptomes are highly similar. Testis single-cell RNA-seq data from Mahyari, *et al*. Germ cell (red) and somatic testicular cell types (green) were defined from differential expression analysis in Mahyari, *et al*. Testis (blue) and germ cell transcriptomes form a clade and are more similar to each other than any other tissue or testicular somatic cell type transcriptome. (**C**) Majority of cells in adult testis are germ cells. Data from Mahyari, *et al*. Average percent of cell types in testis across six normal adult males. Germ cell types are marked in shades of reds; somatic cell types are marked in shades of greens. Percent of each cell type listed in legend. (**D**) CCDF of the probability that PZMs originated from germ cells. The majority of germ cell-specific (gray) and gonosomal (green) PZMs are more likely to have come from germ cells than somatic cells. Gonosomal PZMs have a lower probability than germ cell-specific PZMs because they are ascertained from transcripts broadly expressed across the body and thus are less likely to be germ cell-specific. (**E-F**) Gonosomal and germ cell-specific PZMs have signatures distinct from germline variants. (**E**) VAF distribution of donor germline variants, gonosomal PZMs, and germ cell-specific PZMs. All VAFs are measured from RNA-seq data. (**F**) Distribution of minor allele frequencies of germline variants, gonosomal PZMs, and germ cell-specific PZMs. *P*-values from Mann Whitney *U* tests. To reduce noise in estimates, only donor germline variants with at least 10× coverage were used. For computational ease, only donor germline variants on chr22 were used. Top and bottom of the boxes denote first and third quartiles, respectively; horizontal black lines denote median; whiskers denote 1.5× the interquartile range; and outliers are plotted with transparency to disambiguate overlapping values. (**G**) Gonosomal PZM detection power as function of VAF and expression level. Gonosomal PZMs were simulated in male testis donors across a wide range of VAFs. The percent of simulated variants that were correctly recovered as gonosomal (present in testis and at least one other tissue) was recorded. Coverage of the gonosomal PZM was defined as the median coverage across all sampled tissues in the donor at the particular genomic position. Since coverage is an important component of power, simulated variants were partitioned in three coverage groups. For some coverage groups, gonosomal PZM power curves do not saturate to 100% power due to constraints from differences among tissue transcriptomes and the GTEx tissue sampling structure. NFE AF = allele frequency in non-Finish Europeans.

Next, we determined if germ cell PZMs could be detected in bulk male gonads. We tested if the bulk testis transcriptome was similar to male germ cell transcriptomes. We performed hierarchical clustering of human single-cell testicular biopsy transcriptomes with GTEx tissue transcriptomes (*76*). Mahyari *et al.* classified the single-cell testicular biopsy data into five germ cell types and seven somatic cell types using differential expression analysis (N = 12 total cell types). The bulk testis and germ cell transcriptomes formed a clade and were more similar to each other than to any other tissue or cell type, suggesting that bulk testis expression is a reasonable proxy for male germ cell expression (**Fig. S24B**). This result is expected as 85% of cells in a normal adult testis are germ cells (**Fig. S24C**).

We next determined if the PZMs were likely to have originated from genes expressed by germ cells rather than somatic cells. For each germ cell PZM, we defined the probability that the PZM was in a gene expressed in germ cells as the ratio of expression from germ cell types to the expression of all 12 cell types in the testis. Each cell type was weighted by its prevalence in the testis. The germ cell PZMs were partitioned into gonosomal (i.e., PZM detected in testis and at least one other tissue) and germ cell-specific (i.e., PZM detected only in testis) groups. Both groups were more likely to have originated from germ cells than somatic cells. On average, gonosomal PZMs were 64% likely to be expressed in germ cells and germ cell-specific PZMs were 76% likely to be expressed in germ cells (**Fig. S24D**). The lower probability in gonosomal PZMs is expected: since variants were detected from RNA-seq, putative gonosomal PZMs are likely to be in genes that are broadly expressed across the body whereas putative germ cell-specific PZMs may be broadly expressed or show testis-specific expression.

Together, these results suggest that the majority of PZMs detected from bulk testis RNA-seq are PZMs in germ cells. Thus, we can use such PZMs to measure properties about mutations in the male germ line.

##### *2.10.2 Gonosomal and germ cell-specific PZM class labels are likely correct*

As an extra level of precaution, we next assessed the validity of the gonosomal and germ cell-specific class labels by checking if each group manifested expected properties. The results are summarized in **Table S12**.

For observed gonosomal PZMs, we first determined if the predicted mutations were actually inherited germline variants. The RNA-seq cumulative VAF from all tissues with the gonosomal mutations was significantly lower than the RNA-seq cumulative VAF of germline variants in GTEx donors ($P$-value = 5.3E-44, Mann Whitney $U$ test, **Fig. S24E**). Additionally, gonosomal PZMs had lower population allele frequencies than the GTEx donors' inherited germline variants ($P$-value = 4.9E-13, Mann Whitney $U$ test, **Fig. S24F**). Together, these results suggest that even with complexities due to allele-specific expression, gonosomal PZMs are likely not germline variants.

Next we determined if the predicted gonosomal PZMs were actually germ cell-specific PZMs. This scenario would require false positive PZM calls in at least one other tissue. The FPR of the full set of PZMs was estimated to be 1%. The FPR for germ cell-specific PZMs is likely very similar. Therefore, most predicted gonosomal PZMs are likely not germ cell-specific PZMs.

Lastly, a predicted gonosomal PZM may actually be noise. This case is likely not common as experimental and computational FDRs were reasonably low.

After ruling out mislabelling opportunities, we then confirmed the gonosomal PZM burden had expected properties, namely, the burden was not associated with age. We fit the gonosomal burden with the following linear model:

$$gonosomal\_burden \sim AGE + num\_tissues + self\_reported\_ancestry + median\_RIN + median\_sample\_mutation\_power + \log_{10}(germ\ cell\text{-}specific\ burden + 1) \quad \text{2-4}$$

where $gonosomal\ burden$ was defined as the number of gonosomal PZMs in a donor normalized by the median transcriptome size of all samples in the donor. $median\_RIN$ was defined as the median RIN of all samples in a donor. $median\_sample\_mutation\_power$ was defined in a similar manner. $germ\ cell\text{–}\ specific\ burden$ was defined as the number of germ cell specific PMZs in a donor. Only male donors with genotype data available were used.

As expected, gonosomal PZM burden was not associated with donor age ($P$-value = 0.28). Together, this data suggests that putative gonosomal PZMs are correctly labelled.

We applied a similar framework for the predicted germ-cell specific PZMs. This class is also unlikely to be highly contaminated with germline variants. The predicted germ cell-specific PZM VAFs were significantly smaller than the RNA-seq VAFs of donor germline variants ($P$-value = 2.2E-308, Mann Whitney $U$ test, **Fig. S24E**) and the PZMs had significantly lower minor allele frequencies than donor germline variants ($P$-value = 8.2E-72, Mann Whitney $U$ test, **Fig. S24F**).

In order for a putative germ cell-specific PZM to actually be a gonosomal PZM, the mutation must have been a false negative mutation in all somatic tissues containing the PZM. This likely does not explain the majority of germ cell-specific PZMs since the power to detect gonosomal PZMs was reasonably high for moderately high VAFs (**Fig. S24G**). (Due to their timing, gonosomal PZMs are predicted to have relatively high VAFs.) At the median gonosomal PZM coverage ([100, 1000)) and median gonosomal PZM VAF (0.05), the power to detect gonosomal PZMs was 95%. (Note that these power estimates include the desired case (i.e., a gonosomal PZM is mislabeled as a germ cell-specific PZM) but also include the case when no PZM was detected in any somatic or germline tissue. Therefore, these power estimates will likely be underestimated when just the desired case is considered.)

A predicted germ cell-specific PZM may actually be noise. This case is likely not common as experimental and computational FDRs were reasonably low.

After ruling out potential false positive scenarios, we confirmed germ cell-specific PZMs had properties of true germ-cell specific PZMs. We expected that germ cell-specific PZMs would show an age dependence and would have lower VAFs than gonosomal PZMs.

We fit the germ cell-specific burden with the following linear model:

$$germ\_cell\text{–}specific\_burden \sim AGE + self\_reported\_ancestry + RIN + sample\_mutation\_power \quad (2\text{-}5)$$

where $germ\ cell\text{–}specific\ burden$ was defined as $\frac{\log_{10}(\#\ germ\ cell-specific\ mutations+1)}{transcriptome\ size}$. (This transformation produced the best fitting model of all models tested. Sample batch was not included as a random effect because very few batch levels were repeated and the models resulted in singular model fits suggesting overfitting.) Again, only male donors with genotype data available were used.  As expected, germ cell-specific PZM burden was positively associated with donor age (*P*-value = 0.03). Additionally, as expected, germ cell-specific PZMs had lower VAFs than gonosomal PZMs (*P*-value = 1.3E-14, Mann-Whitney *U* test, **Fig. S24E**). Collectively, these results suggest that the gonsomal and germ cell-specific PZM class labels are likely accurate; however there may still be some misclassification at an individual variant level.

#### *2.10.3 Distribution and deleteriousness of gonosomal and germ cell-specific PZM mutation burdens across male donors*

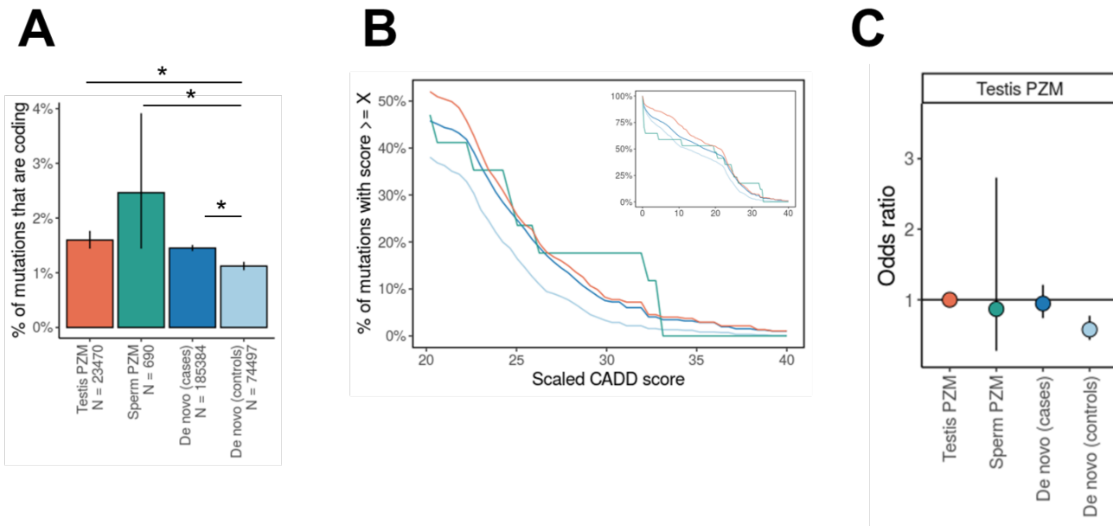

**Fig. S25. Independent germ cell datasets also show purging of deleterious mutations during the germ cell life cycle.** (**A**) Proportion of coding mutations by dataset. Horizontal lines represent results from testing if the proportion of coding mutations in a given dataset is different than *de novo* (controls). Statistically significant differences (*P*-value $\leq 0.05$) are annotated with "*". N is the number of single-nucleotide mutations in the dataset. (**B**) PHRED-scaled CADD score CCDFs of the independent germ cell datasets. Distributions with larger areas under the curve have a larger fraction of deleterious mutations. **Inset**: CCDF for CADD scores [0,40]; **main**: CCDF for high CADD scores [20,40]. (**C**) Relative odds of detecting deleterious mutations across germ cell datasets compared to independent testis PZM dataset. Datapoints colored by dataset. Horizontal black line at odds ratio = 1 denotes no difference in odds. Error bars denote 95% CIs. Independent testis PZM dataset from Moore *et al.*(*77*); independent sperm PZM dataset from Yang *et al.*(*78*)

#### *2.10.4 Study design and model assumptions did not affect germline surrogate tissue analyses*

##### 2.10.4.1 Donor genotype status does not affect prenatal PZM burden

An important confounder in the gonosomal PZM surrogate tissue analysis is that blood was used to define (and later remove) germline variants from the donors. These predicted germline variants might actually be high VAF PZMs that were incorrectly labelled as germline variants during germline variant calling on blood WGS. Thus, at first glance, the result that blood was the worst tissue to detect gonosomal PZMs may be explained by blood having a higher rate of gonosomal PZMs removed from the germline variant filter.

The GTEx study design offered a unique opportunity to examine the validity of this hypothesis: for 6% of GTEx donors, a non-blood tissue was used for genotyping and for 12% of donors, no genotyping data was available. (We note that while we excluded non-genotyped donors from all germline analyses, including non-genotyped donors for this analysis allowed us to investigate the strongest effect of genotyping on PZM variant calling.) To determine if using blood for germline variant calling affected the surrogate analysis, we compared various mutation metrics between donors who were genotyped using blood DNA, donors genotyped using non-blood DNA, and donors not genotyped. To increase the power of detecting a significant difference, we expanded the analysis to all prenatal PZMs rather than just the subset of gonosomal PZMs.

If germline variant filtering removed [high VAF] prenatal PZMs from blood, we would expect blood genotyped donors to have fewer prenatal PZMs detected in blood than non-blood genotyped donors and much fewer prenatal PZMs than non-genotyped donors. We fit the following model:

$$\frac{\#\ prenatal\ PZMs\ in\ blood + 1}{transcriptome\_size} \sim genotype\_source + RIN + AGE + SEX + self\_reported\_ancestry + sample\_mutation\_power + (1|BATCH) \qquad 2\text{-}6$$

where $genotype\_source$ was blood, not_blood, or not_genotyped. We fit the data again with **equation 2-6**, only this time, we dropped the $genotype\_source$ covariate. Models were fit using R package lme4 (v1.1-26). Next, we determined if including the genotype source improved the model fit by comparing the AIC of the nested models. Adding $genotype\_source$ did not substantially change the AIC (percent change in AIC = 0.1%). This suggests that the use of blood for germline variant filtering did not affect the prenatal PZM burden and thus poor detection of gonosomal PZMs in blood is not the result of overzealous germline variant filtering in blood.

###### 2.10.4.2 Step 2 of LachesisDetect does not reduce inter-tissue variation in VAF

We investigated if the similarity of tissue VAFs for a given gonosomal PZMs could be an artefact of LachesisDetect. An assumption of the mutation calling pipeline is that if a mutation is found in multiple tissues of the same donor, the tissue VAFs will be similar. Recall that in step 1 of LachesisDetect, the variant calling algorithm identifies mutations in a sample *independent* of all other samples. In step 2, the algorithm uses the aforementioned assumption to perform more sensitive variant calling.

We defined the inter-tissue variation in VAF of a given multi-tissue PZM as the variation in VAF across all of the donor's tissues with the PZM. To test if step 2 biased multi-tissue PZMs to have more similar VAFs, we compared the inter-tissue variation in VAF of multi-tissue PZMs detected in step 1 with multi-tissue PZMs detected in step 2. If step 2 artificially shrank the tissue VAFs to be more similar, we would expect a reduction in inter-tissue VAF variation after step 2.

To increase power, we used all prenatal PZMs rather than just the gonosomal PZM subset. We used the median absolute deviation (MAD) of VAFs normalized by the median VAF to estimate the variation in VAFs within a multi-tissue PZM. MAD was chosen to robustly measure variation due to the relatively small number of tissues in a multi-tissue PZM. We fit the following linear model to the data:

$$normalized\ MAD \sim PZM\_detection\_step + num\_tissues + PZM\_detection\_step * num\_tissues \qquad 2\text{-}7$$

The relationship between normalized MAD and the covariates is shown in **Fig. S26A**. The interaction term was significant, i.e., the relationship between normalized MAD and which step identified the PZM in multiple tissues was different when the number of tissues increased. When the number of tissues was small (< 8), there was no difference in normalized MAD between step 1 and step 2 multi-tissue PZMs. When the number of tissues was large (≥ 8), there was a significant difference; however, the sign was in the opposite direction from what we expected: step 2 PZMs had greater variation than step 1 PZMs (**Fig. S26B**). The majority of multi-tissue PZMs (94%) occurred in the former case.

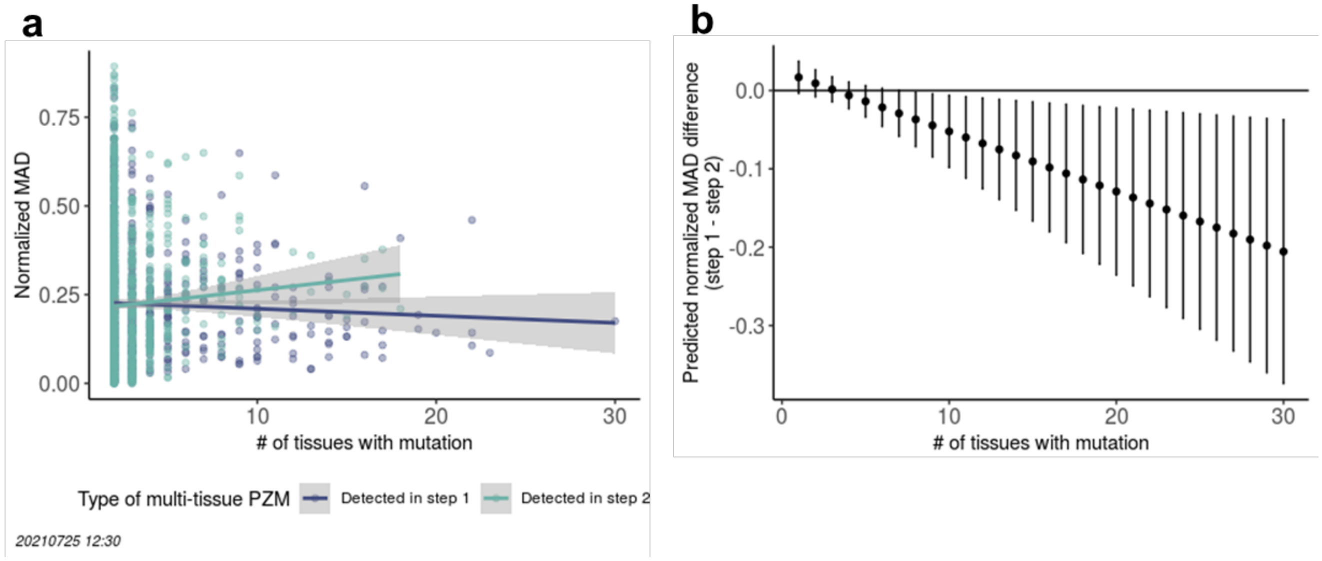

**Fig. S26. Step 2 of LachesisDetect does not reduce intra-tissue variation in VAF.** (**A**) Normalized MAD of multi-tissue PZMs as a function of the number of tissues containing the mutation and which step the multi-tissue evidence was detected in. (**B**) Predicted difference in normalized MAD between step 1 and step 2 multi-tissue PZMs as a function of the number of tissues containing the mutation. Error bars represent 95% CIs. MAD = median absolute deviation.

Together, this analysis shows that step 2 does not artificially decrease variation in VAF. The implications are two fold: first the assumption about VAF similarity is appropriate (MADs were generally small) and second, the similarity of germ cell and somatic tissue gonosomal PZM VAFs is likely a biological rather than technical result.

#### 3 Supplementary tables & files

**Table S1. List of filters applied to the PZM call set and the number of PZMs removed or added by each filter.**

| Filter | Variants removed | | Variants added | | Remaining # of variants |
|---|---|---|---|---|---|
| | # | % | # | % | |
| Input (# of sites with ≥ 2X alternative coverage) | | | | | 927,604,333 |
| Blacklist samples<br>- Low RIN<br>- GTEx blacklist tissue<br>- Transplanted GTEx tissue | 93,237,551 | 10.1% | | | 834,366,782 |
| Blacklist genomic regions<br>- Known RNA editing sites<br>- Immunoglobulin genes<br>- Variants near exon boundaries | 87,176,943 | 10.4% | | | 747,189,839 |
| Variants that overlap with GTEx germline SNVs and indels | 298,581,071 | 40.0% | | | 448,608,768 |
| Total coverage ≤ 20X | 18,803,931 | 4.2% | | | 429,804,837 |
| Putative germline variants | 1,331,121 | 0.3% | | | 428,473,716 |
| Alternative coverage ≤ 5X | 414,983,074 | 96.9% | | | 13,490,642 |
| Putative RNA editing sites | 2,095,884 | 15.5% | | | 11,394,758 |
| Sites with high nucleotide diversity | 7,140,260 | 62.7% | | | 4,254,498 |
| PZM FDR correction | 3,100,562 | 72.9% | | | 1,153,936 |
| Biased alignment metrics | 448,032 | 38.8% | | | 705,904 |
| In segmental duplication | 83,632 | 11.8% | | | 622,272 |
| In repeat masked region | 32,453 | 5.2% | | | 589,819 |
| Hypermutated samples | 455,466 | 77.2% | | | 134,353 |
| Call multi-tissue variants | 2,005 | 1.5% | 30,731 | 22.9% | 163,079 |
| Balance mutation burden in genotyped and non-genotyped donors | 33,289 | 20.4% | | | 129,790 |
| Large background error rate (pilot validation dataset) | 43,131 | 33.2% | | | 86,659 |
| Large background error rate (all validation datasets) | 30,074 | 34.7% | | | 56,585 |

**Table S2. List of sample filters and the number of samples removed by each filter.**

| Filter | Samples removed | | Remaining # of samples |
|---|---|---|---|
| | # | % | |
| Input | | | 17,382 |
| Low RIN<br>(RIN < 6) | 1,674 | 10% | 15,708 |
| GTEx tissues with poor overall quality (Bladder, Spleen, Cervix - Ectocervix, Cervix - Endocervix, Fallopian Tube, Kidney - Medulla) | 230 | 1% | 15,478 |
| Sample derived from a transplanted tissue | 0 | 0% | 15,478 |
| Hypermutated samples | 806 | 5% | 14,672 |

**Table S3. Experimental and in silico validation results.***Sheet Validation results summary:* Summary of results for experimental and in silico validation experiments for non-hypermutated and hypermutated samples.

- *Sheet Validation results:* Table of variant information (e.g., genomic coordinate and validation status) for the PZMs where validation was attempted across the four experimental validation datasets. Due to privacy, sample IDs were converted to deidentified sample IDs.
- *Sheet Valid. results - column defs:* Description of columns used in Sheet Validation results.
- *Sheet Amp.-seq pilot samples summary:* Number of non-hypermutated and hypermutated samples used in the amplicon-seq pilot validation dataset.
- *Sheet Amp.-seq pilot samples:* List of samples used in the amplicon-seq pilot validation dataset.
- *Sheet Amp.-seq large samples summary:* Number of non-hypermutated and hypermutated samples used in the amplicon-seq large validation dataset.
- *Sheet Amp.-seq large samples:* List of samples used in the amplicon-seq large validation dataset.
- *Sheet ENCODE samples summary:* Number of non-hypermutated and hypermutated samples used in the ENCODE validation datasets.
- *Sheet ENCODE large samples:* List of samples used in the ENCODE validation datasets.

**Table S4. List of regions amplified for the pilot amplicon-seq validation experiment.** Coordinates are in hg19.

**Table S5. List of regions amplified for the large-scale amplicon-seq validation experiment.** Coordinates are in hg38.

**Table S6. List of primer pairs used for the amplicon-seq large validation experiment.**

**Table S7. List of PZMs.** Spreadsheet has the following sheets:

- *Sheet List of mutations:* Table of PZM information (e.g., genomic coordinate and sequencing read coverage). Due to privacy, sample IDs and donor IDs were deidentified.
- *Sheet Column definitions:* Description of columns used in Sheet List of mutations.

**Table S8. GTEx coloring convention.** The same coloring convention used in GTEx was used(*15*).

| Tissue | Color |
|---|---|
| Adipose - Subcutaneous | |
| Adipose - Visceral (Omentum) | |
| Adrenal Gland | |
| Artery - Aorta | |
| Artery - Coronary | |
| Artery - Tibial | |
| Brain - Amygdala | |
| Brain - Anterior cingulate cortex (BA24) | |
| Brain - Caudate (basal ganglia) | |
| Brain - Cerebellar Hemisphere | |
| Brain - Cerebellum | |
| Brain - Cortex | |
| Brain - Frontal Cortex (BA9) | |
| Brain - Hippocampus | |
| Brain - Hypothalamus | |
| Brain - Nucleus accumbens (basal ganglia) | |
| Brain - Putamen (basal ganglia) | |
| Brain - Spinal cord (cervical c-1) | |
| Brain - Substantia nigra | |
| Breast - Mammary Tissue | |
| Cells - EBV-transformed lymphocytes | |
| Cells - Cultured fibroblasts | |
| Colon - Sigmoid | |
| Colon - Transverse | |
| Esophagus - Gastroesophageal Junction | |
| Esophagus - Mucosa | |
| Esophagus - Muscularis | |
| Heart - Atrial Appendage | |
| Heart - Left Ventricle | |
| Kidney - Cortex | |
| Liver | |
| Lung | |
| Minor Salivary Gland | |
| Muscle - Skeletal | |
| Nerve - Tibial | |
| Ovary | |
| Pancreas | |
| Pituitary | |
| Prostate | |
| Skin - Not Sun Exposed (Suprapubic) | |
| Skin - Sun Exposed (Lower leg) | |
| Small Intestine - Terminal Ileum | |
| Stomach | |
| Testis | |
| Thyroid | |
| Uterus | |
| Vagina | |
| Whole Blood | |

**Table S9. U.S. cancer incidence rates related to tissues with significant differences in mutation burden among ancestry groups.**

**Table S10. Summary of datasets used to validate deleteriousness patterns and whether results were validated**.NA = Not applicable.

| | **This study** | **TwinsUK** | **García-Nieto *et al.*** | **Yizhak *et al.*** | **Brazhnik *et al.*** |
|---|---|---|---|---|---|
| **Reference** | NA | Unpublished | (*50*) | (*49*) | (*51*) |
| **Uses RNA** | Yes | Yes | Yes | Yes | No |
| **Uses GTEx data** | Yes (v8) | No | Yes (v7) | Yes (v7) | No |
| **Variant calling method** | Lachesis | Lachesis | Custom method | RNA-MuTect | VarScan2, MuTect2, & HaplotypeCaller |
| **Results validated** | NA | Yes | No | Yes | Yes |

**Table S11. List of CHIP mutations detected in GTEx samples.**Genomic coordinates are relative to GRCh38 and are 0-based with exclusive end coordinates. GTEx recurrence is defined as the number of times the mutation was detected in GTEx blood and LCL samples. COSMIC recurrence is defined as the number of times the mutation was detected in haematopoietic and lymphoid tissue cancer samples. COSMIC recurrence percentile is defined as the percent of recurrent COSMIC mutations with the same or lower recurrence as the mutation.

| chr | start | end | ref_allele | alt_allele | gene_name | HGVSG | HGVSC | HGVSP | cosmic_mutation_id | tissue | recurrence.gtex | recurrence.cosmic | recurrence_percentile.cosmic |
|---|---|---|---|---|---|---|---|---|---|---|---|---|---|
| chr3 | 38141149 | 38141150 | T | C | MYD88 | 3:g.38141150T>C | ENST00000417037.6:c.818T>C | ENSP00000401399.2:p.Leu273Pro | COSV57169334 | Cells - EBV-transformed lymphocytes | 1 | 1969 | 0.99998 |
| chr15 | 90088701 | 90088702 | C | T | IDH2 | 15:g.90088702C>T | ENST00000330062.7:c.419G>A | ENSP00000331897.3:p.Arg140Gln | COSV57468751 | Cells - EBV-transformed lymphocytes | 1 | 815 | 0.99991 |
| chr17 | 42322442 | 42322443 | T | A | STAT3 | 17:g.42322443T>A | ENST00000264657.9:c.1940A>T | ENSP00000264657.4:p.Asn647Ile | COSV52882818 | Cells - EBV-transformed lymphocytes | 1 | 18 | 0.99630 |
| chr12 | 92145426 | 92145427 | G | A | BTG1 | 12:g.92145427G>A | ENST00000256015.4:c.109C>T | ENSP00000256015.3:p.Leu37= | COSV55458204 | Cells - EBV-transformed lymphocytes | 2 | 6 | 0.98268 |
| chr15 | 44711580 | 44711581 | T | C | B2M | 15:g.44711581T>C | ENST00000558401.5:c.35T>C | ENSP00000452780.1:p.Leu12Pro | COSV62563197 | Cells - EBV-transformed lymphocytes | 1 | 6 | 0.98268 |
| chr17 | 42322400 | 42322401 | T | A | STAT3 | 17:g.42322401T>A | ENST00000264657.9:c.1982A>T | ENSP00000264657.4:p.Asp661Val | COSV52886283 | Whole Blood | 1 | 6 | 0.98268 |

**Table S12. Confusion matrix for potential mislabelling of gonosomal and germ cell-specific PZMs.**Matrix cells represent all possible correct and incorrect labellings of predicted gonosomal and germ cell-specific PZMs. Cells that represent correctly labelled classes are shaded in green; incorrectly labelled classes are shaded in red. Cell text describes the expected patterns in the data if the scenario were true. Check marks denote the pattern was observed; X's denote the pattern was not observed. Since all of the correct labelling patterns were observed and none of the incorrect labelling patterns were observed, the gonosomal and germ cell-specific PZM datasets are likely correctly labeled.

| | | Predicted condition | |
|---|---|---|---|
| | | **Gonosomal PZM** | **Germ cell-specific PZM** |
| **True condition** | **Gonosomal PZM** | ✓ Burden not associated with age<br>✓ VAF < 50% | ✗ Gonosomal PZM detection power is low |
| | **Germ cell-specific PZM** | ✗ FPR is high | ✓ Burden associated with age<br>✓ VAF << 50%<br>✓ Germ cell-specific VAFs < gonosomal VAFs |
| | **Inherited germline variant** | ✗ PZM VAF similar to RNA-seq VAF of donor germline variants<br>✗ PZM MAF similar to MAF of donor germline variants | ✗ PZM VAF similar to RNA-seq VAF of donor germline variants<br>✗ PZM MAF similar to MAF of donor germline variants |
| | **Noise** | ✗ Experimental and computational validation is poor. | |

**Table S13. Results from testing for differences in germ cell mutation spectra.** Mutation *spectra* q-value = Benjamini-Hocherberg corrected *P*-value for testing if a given pair of datasets have the same mutation spectra (Chi-square test). Mutation *type* q-value = Benjamini-Hochberg corrected *P*-value for testing if the proportion of given mutation type is the same in a given pair of datasets (Chi-square test). Significant statistical results at an FDR of 5% are bolded.

| **Dataset 1** | **Dataset 2** | **Mutation spectra q-value** | **Mutation type q-value** | |
|---|---|---|---|---|
| De novo (case) | De novo (control) | **7.22E-03** | **C>A** | **4.28E-03** |
| De novo (case) | De novo (control) | | C>G | 4.25E-01 |
| De novo (case) | De novo (control) | | **C>T** | **7.50E-03** |
| De novo (case) | De novo (control) | | T>A | 9.79E-01 |
| De novo (case) | De novo (control) | | T>C | 7.26E-01 |
| De novo (case) | De novo (control) | | T>G | 4.90E-01 |
| De novo (case) | Germ cell-specific | **8.33E-04** | **C>A** | **2.73E-03** |
| De novo (case) | Germ cell-specific | | **C>G** | **2.73E-03** |
| De novo (case) | Germ cell-specific | | **C>T** | **7.50E-03** |
| De novo (case) | Germ cell-specific | | T>A | 7.61E-01 |
| De novo (case) | Germ cell-specific | | **T>C** | **2.73E-03** |
| De novo (case) | Germ cell-specific | | **T>G** | **2.25E-02** |
| De novo (case) | Gonosomal | **8.33E-04** | **C>A** | **2.73E-03** |
| De novo (case) | Gonosomal | | C>G | 5.51E-01 |
| De novo (case) | Gonosomal | | C>T | 8.62E-02 |

| | | | | |
|---|---|---|---|---|
| De novo (case) | Gonosomal | | T>A | 3.41E-01 |
| De novo (case) | Gonosomal | | T>C | 1.47E-01 |
| De novo (case) | Gonosomal | | T>G | 8.65E-01 |
| De novo (case) | Sperm | **8.33E-04** | C>A | 7.84E-01 |
| De novo (case) | Sperm | | C>G | 1.45E-01 |
| De novo (case) | Sperm | | C>T | 2.81E-01 |
| De novo (case) | Sperm | | T>A | **2.73E-03** |
| De novo (case) | Sperm | | T>C | 5.08E-01 |
| De novo (case) | Sperm | | T>G | 3.20E-01 |
| De novo (control) | Germ cell-specific | **8.33E-04** | **C>A** | **2.73E-03** |
| De novo (control) | Germ cell-specific | | **C>G** | **4.28E-03** |
| De novo (control) | Germ cell-specific | | **C>T** | **2.73E-03** |
| De novo (control) | Germ cell-specific | | T>A | 7.42E-01 |
| De novo (control) | Germ cell-specific | | **T>C** | **2.73E-03** |
| De novo (control) | Germ cell-specific | | **T>G** | **2.00E-02** |
| De novo (control) | Gonosomal | **8.33E-04** | **C>A** | **2.73E-03** |
| De novo (control) | Gonosomal | | C>G | 4.90E-01 |
| De novo (control) | Gonosomal | | C>T | 2.71E-01 |
| De novo (control) | Gonosomal | | T>A | 3.38E-01 |
| De novo (control) | Gonosomal | | T>C | 1.31E-01 |
| De novo (control) | Gonosomal | | T>G | 1.00E+00 |
| De novo (control) | Sperm | **8.33E-04** | C>A | 1.00E+00 |
| De novo (control) | Sperm | | C>G | 1.45E-01 |
| De novo (control) | Sperm | | C>T | 5.51E-01 |
| De novo (control) | Sperm | | T>A | **2.73E-03** |
| De novo (control) | Sperm | | T>C | 5.08E-01 |
| De novo (control) | Sperm | | T>G | 3.20E-01 |
| Gonosomal | Germ cell-specific | **1.85E-02** | C>A | 8.21E-02 |
| Gonosomal | Germ cell-specific | | C>G | 5.08E-01 |
| Gonosomal | Germ cell-specific | | **C>T** | **1.23E-02** |
| Gonosomal | Germ cell-specific | | T>A | 5.08E-01 |
| Gonosomal | Germ cell-specific | | T>C | 9.32E-01 |
| Gonosomal | Germ cell-specific | | T>G | 1.63E-01 |
| Gonosomal | Sperm | **5.00E-03** | **C>A** | **2.05E-02** |
| Gonosomal | Sperm | | C>G | 5.94E-01 |
| Gonosomal | Sperm | | C>T | 8.46E-01 |
| Gonosomal | Sperm | | T>A | 2.53E-01 |
| Gonosomal | Sperm | | T>C | 7.63E-02 |
| Gonosomal | Sperm | | T>G | 3.20E-01 |
| Sperm | Germ cell-specific | **1.43E-03** | C>A | 2.71E-01 |
| Sperm | Germ cell-specific | | C>G | 1.00E+00 |

| | | | | |
|---|---|---|---|---|
| Sperm | Germ cell-specific | | C>T | 5.43E-02 |
| Sperm | Germ cell-specific | | T>A | **2.73E-03** |
| Sperm | Germ cell-specific | | T>C | **4.28E-03** |
| Sperm | Germ cell-specific | | T>G | 1.00E+00 |

8. The GTEx Consortium, K. G. Ardlie, D. S. Deluca, A. V. Segre, T. J. Sullivan, T. R. Young, E. T. Gelfand, C. A. Trowbridge, J. B. Maller, T. Tukiainen, M. Lek, L. D. Ward, P. Kheradpour, B. Iriarte, Y. Meng, C. D. Palmer, T. Esko, W. Winckler, J. N. Hirschhorn, M. Kellis, D. G. MacArthur, G. Getz, A. A. Shabalin, G. Li, Y.-H. Zhou, A. B. Nobel, I. Rusyn, F. A. Wright, T. Lappalainen, P. G. Ferreira, H. Ongen, M. A. Rivas, A. Battle, S. Mostafavi, J. Monlong, M. Sammeth, M. Mele, F. Reverter, J. M. Goldmann, D. Koller, R. Guigo, M. I. McCarthy, E. T. Dermitzakis, E. R. Gamazon, H. K. Im, A. Konkashbaev, D. L. Nicolae, N. J. Cox, T. Flutre, X. Wen, M. Stephens, J. K. Pritchard, Z. Tu, B. Zhang, T. Huang, Q. Long, L. Lin, J. Yang, J. Zhu, J. Liu, A. Brown, B. Mestichelli, D. Tidwell, E. Lo, M. Salvatore, S. Shad, J. A. Thomas, J. T. Lonsdale, M. T. Moser, B. M. Gillard, E. Karasik, K. Ramsey, C. Choi, B. A. Foster, J. Syron, J. Fleming, H. Magazine, R. Hasz, G. D. Walters, J. P. Bridge, M. Miklos, S. Sullivan, L. K. Barker, H. M. Traino, M. Mosavel, L. A. Siminoff, D. R. Valley, D. C. Rohrer, S. D. Jewell, P. A. Branton, L. H. Sobin, M.

Barcus, L. Qi, J. McLean, P. Hariharan, K. S. Um, S. Wu, D. Tabor, C. Shive, A. M. Smith, S. A. Buia, A. H. Undale, K. L. Robinson, N. Roche, K. M. Valentino, A. Britton, R. Burges, D. Bradbury, K. W. Hambright, J. Seleski, G. E. Korzeniewski, K. Erickson, Y. Marcus, J. Tejada, M. Taherian, C. Lu, M. Basile, D. C. Mash, S. Volpi, J. P. Struewing, G. F. Temple, J. Boyer, D. Colantuoni, R. Little, S. Koester, L. J. Carithers, H. M. Moore, P. Guan, C. Compton, S. J. Sawyer, J. P. Demchok, J. B. Vaught, C. A. Rabiner, N. C. Lockhart, K. G. Ardlie, G. Getz, F. A. Wright, M. Kellis, S. Volpi, E. T. Dermitzakis, The Genotype-Tissue Expression (GTEx) pilot analysis: Multitissue gene regulation in humans. *Science*. **348**, 648–660 (2015).

18. Y. S. Ju, I. Martincorena, M. Gerstung, M. Petljak, L. B. Alexandrov, R. Rahbari, D. C. Wedge, H. R. Davies, M. Ramakrishna, A. Fullam, S. Martin, C. Alder, N. Patel, S. Gamble, S. O'Meara, D. D. Giri, T. Sauer, S. E. Pinder, C. A. Purdie, Å. Borg, H. Stunnenberg, M. van de Vijver, B. K. T. Tan, C. Caldas, A. Tutt, N. T. Ueno, L. J. van 't

Veer, J. W. M. Martens, C. Sotiriou, S. Knappskog, P. N. Span, S. R. Lakhani, J. E. Eyfjörd, A.-L. Børresen-Dale, A. Richardson, A. M. Thompson, A. Viari, M. E. Hurles, S. Nik-Zainal, P. J. Campbell, M. R. Stratton, Somatic mutations reveal asymmetric cellular dynamics in the early human embryo. *Nature*. **543**, 714–718 (2017).

29. The National Cancer Institute, The Biospecimen Research Database, (available at http://biospecimens.cancer.gov/brd).

30. J. E. Sulston, E. Schierenberg, J. G. White, J. N. Thomson, The embryonic cell lineage of the nematode Caenorhabditis elegans. *Dev. Biol.* **100**, 64–119 (1983).

31. R. Edgar, Y. Mazor, A. Rinon, J. Blumenthal, Y. Golan, E. Buzhor, I. Livnat, S. Ben-Ari, I. Lieder, A. Shitrit, Y. Gilboa, A. Ben-Yehudah, O. Edri, N. Shraga, Y. Bogoch, L. Leshansky, S. Aharoni, M. D. West, D. Warshawsky, R. Shtrichman, LifeMap Discovery$^{TM}$: the embryonic development, stem cells, and regenerative medicine research portal. *PLoS One*. **8**, e66629 (2013).

32. Editorial office Of journal "Morphologia," Langman's Medical Embryology. 14th Edition, 2018 Author: T.W. Sadler. *Morphologia*. **13**, 90–96 (2019).

33. T. H. N. Teshima, R. F. Ianez, C. M. Coutinho-Camillo, M. E. Buim, F. A. Soares, S. V. Lourenço, Development of human minor salivary glands: expression of mucins according to stage of morphogenesis. *J. Anat.* **219**, 410–417 (2011).

34. GI Motility online, (available at https://www.nature.com/gimo/).

40. K. J. Karczewski, L. C. Francioli, G. Tiao, B. B. Cummings, J. Alföldi, Q. Wang, R. L. Collins, K. M. Laricchia, A. Ganna, D. P. Birnbaum, L. D. Gauthier, H. Brand, M. Solomonson, N. A. Watts, D. Rhodes, M. Singer-Berk, E. M. England, E. G. Seaby, J. A. Kosmicki, R. K. Walters, K. Tashman, Y. Farjoun, E. Banks, T. Poterba, A. Wang, C. Seed, N. Whiffin, J. X. Chong, K. E. Samocha, E. Pierce-Hoffman, Z. Zappala, A. H. O'Donnell-Luria, E. V. Minikel, B. Weisburd, M. Lek, J. S. Ware, C. Vittal, I. M. Armean, L. Bergelson, K. Cibulskis, K. M. Connolly, M. Covarrubias, S. Donnelly, S. Ferriera, S. Gabriel, J. Gentry, N. Gupta, T. Jeandet, D. Kaplan, C. Llanwarne, R. Munshi, S. Novod, N. Petrillo, D. Roazen, V. Ruano-Rubio, A. Saltzman, M. Schleicher, J. Soto, K. Tibbetts, C. Tolonen, G. Wade, M. E. Talkowski, Genome Aggregation Database Consortium, B. M.

Neale, M. J. Daly, D. G. MacArthur, The mutational constraint spectrum quantified from variation in 141,456 humans. *Nature*. **581**, 434–443 (2020).

55. A. Weyrich, *Curr. Protoc. Mol. Biol.*, in press, doi:10.1002/0471142727.mb0213s98.

56. C. Zong, S. Lu, A. R. Chapman, X. S. Xie, Genome-wide detection of single-nucleotide and copy-number variations of a single human cell. *Science*. **338**, 1622–1626 (2012).

57. K. Cibulskis, M. S. Lawrence, S. L. Carter, A. Sivachenko, D. Jaffe, C. Sougnez, S. Gabriel, M. Meyerson, E. S. Lander, G. Getz, Sensitive detection of somatic point mutations in impure and heterogeneous cancer samples. *Nat. Biotechnol.* **31**, 213–219 (2013).

58. D. C. Koboldt, K. Chen, T. Wylie, D. E. Larson, M. D. McLellan, E. R. Mardis, G. M. Weinstock, R. K. Wilson, L. Ding, VarScan: variant detection in massively parallel sequencing of individual and pooled samples. *Bioinformatics*. **25**, 2283–2285 (2009).

59. D. Bates, M. Mächler, B. Bolker, S. Walker, Fitting Linear Mixed-Effects Models using lme4. *arXiv [stat.CO]* (2014), (available at http://arxiv.org/abs/1406.5823).

60. B. B. Campbell, N. Light, D. Fabrizio, M. Zatzman, F. Fuligni, R. de Borja, S. Davidson, M. Edwards, J. A. Elvin, K. P. Hodel, W. J. Zahurancik, Z. Suo, T. Lipman, K. Wimmer, C. P. Kratz, D. C. Bowers, T. W. Laetsch, G. P. Dunn, T. M. Johanns, M. R. Grimmer, I. V. Smirnov, V. Larouche, D. Samuel, A. Bronsema, M. Osborn, D. Stearns, P. Raman, K. A. Cole, P. B. Storm, M. Yalon, E. Opocher, G. Mason, G. A. Thomas, M. Sabel, B. George, D. S. Ziegler, S. Lindhorst, V. M. Issai, S. Constantini, H. Toledano, R. Elhasid, R. Farah, R. Dvir, P. Dirks, A. Huang, M. A. Galati, J. Chung, V. Ramaswamy, M. S. Irwin, M. Aronson, C. Durno, M. D. Taylor, G. Rechavi, J. M. Maris, E. Bouffet, C. Hawkins, J. F. Costello, M. S. Meyn, Z. F. Pursell, D. Malkin, U. Tabori, A. Shlien, Comprehensive analysis of hypermutation in human cancer. *Cell*. **171**, 1042-1056.e10 (2017).

69. P. Godoy, N. J. Hewitt, U. Albrecht, M. E. Andersen, N. Ansari, S. Bhattacharya, J. G. Bode, J. Bolleyn, C. Borner, J. Böttger, A. Braeuning, R. A. Budinsky, B. Burkhardt, N. R. Cameron, G. Camussi, C.-S. Cho, Y.-J. Choi, J. Craig Rowlands, U. Dahmen, G. Damm, O. Dirsch, M. T. Donato, J. Dong, S. Dooley, D. Drasdo, R. Eakins, K. S. Ferreira, V. Fonsato, J. Fraczek, R. Gebhardt, A. Gibson, M. Glanemann, C. E. P. Goldring, M. J. Gómez-Lechón, G. M. M. Groothuis, L. Gustavsson, C. Guyot, D. Hallifax, S. Hammad, A. Hayward, D. Häussinger, C. Hellerbrand, P. Hewitt, S. Hoehme, H.-G. Holzhütter, J. B. Houston, J. Hrach, K. Ito, H. Jaeschke, V. Keitel, J. M. Kelm, B. Kevin Park, C. Kordes, G. A. Kullak-Ublick, E. L. LeCluyse, P. Lu, J. Luebke-Wheeler, A. Lutz, D. J. Maltman, M. Matz-Soja, P. McMullen, I. Merfort, S. Messner, C. Meyer, J. Mwinyi, D. J. Naisbitt, A. K. Nussler, P. Olinga, F. Pampaloni, J. Pi, L. Pluta, S. A. Przyborski, A. Ramachandran, V. Rogiers, C. Rowe, C. Schelcher, K. Schmich, M. Schwarz, B. Singh, E. H. K. Stelzer, B. Stieger, R. Stöber, Y. Sugiyama, C. Tetta, W. E. Thasler, T. Vanhaecke, M. Vinken, T. S. Weiss, A. Widera, C. G. Woods, J. J. Xu, K. M. Yarborough, J. G. Hengstler, Recent advances in 2D and 3D in vitro systems using primary hepatocytes, alternative hepatocyte

sources and non-parenchymal liver cells and their use in investigating mechanisms of hepatotoxicity, cell signaling and ADME. *Arch. Toxicol.* **87**, 1315–1530 (2013).

77. L. Moore, A. Cagan, T. H. H. Coorens, M. D. C. Neville, R. Sanghvi, M. A. Sanders, T. R. W. Oliver, D. Leongamornlert, P. Ellis, A. Noorani, T. J. Mitchell, T. M. Butler, Y. Hooks, A. Y. Warren, M. Jorgensen, K. J. Dawson, A. Menzies, L. O'Neill, C. Latimer, M. Teng, R. van Boxtel, C. A. Iacobuzio-Donahue, I. Martincorena, R. Heer, P. J. Campbell, R. C.

Fitzgerald, M. R. Stratton, R. Rahbari, The mutational landscape of human somatic and germline cells. *Nature* (2021), doi:10.1038/s41586-021-03822-7.

78. X. Yang, M. W. Breuss, X. Xu, D. Antaki, K. N. James, V. Stanley, L. L. Ball, R. D. George, S. A. Wirth, B. Cao, A. Nguyen, J. McEvoy-Venneri, G. Chai, S. Nahas, L. Van Der Kraan, Y. Ding, J. Sebat, J. G. Gleeson, Developmental and temporal characteristics of clonal sperm mosaicism. *Cell*. **184**, 4772-4783.e15 (2021).
